# A conditional upper-bound design map for rhodopsin-augmented electro-microbial CO_2_-to-biomass conversion

**DOI:** 10.64898/2026.10.09.758023

**Authors:** Haris Saeed, Wei E. Huang, Aidong Yang

## Abstract

The value of rhodopsin proton pumps as light harvesters in electro-microbial CO_2_-to-biomass conversion depends on molecular photophysics, pathway energetics and reactor light delivery. We couple these scales for *Cupriavidus necator*, comparing Calvin–Benson–Bassham (CBB) and reductive glycine (rGlyP) assimilation. The model predicts a 17.9% dark rGlyP yield advantage, close to 16.5% measured in chemostats. At 568 nm, 100 PPFD and 45% membrane occupancy, light-derived ATP raises yield by 17% for rGlyP and 14% for CBB. Charging CO_2_ capture and formate electrosynthesis, a retinal-only reactor reaches 3.29% efficiency versus 2.56% in the dark. A carotenoid antenna raises this to 3.52% at the model-derived 70% transfer estimate and helps in 93.2% of uncertainty samples, but lowers efficiency at the thermodynamic limit by diverting photons from photovoltaics. Point-source CO_2_ cuts the relative light benefit from 38% to 8%. Rhodopsin is therefore most useful when upstream formate is energy-intensive, not as a general efficiency boost.

**Highlights:**

- Predicted 17.9% dark yield gain closely matches chemostat measurements
- Light-derived energy replaces respiratory formate use at similar rates
- A carotenoid antenna helps in 93.2% of matched uncertainty samples
- Point-source carbon dioxide cuts the relative light benefit to 8%

**eTOC Blurb:** Light-driven proton pumps could reduce the formate that carbon-fixing bacteria consume to make ATP. Saeed et al. link molecular, cellular and reactor constraints, showing how pathway choice, absorption bandwidth and carbon source determine the efficiency of rhodopsin-assisted CO_2_ conversion.

## Introduction

Hybrid artificial photosynthesis separates two tasks: photovoltaic electrocatalysis converts sunlight and CO_2_ into a soluble feedstock, and a microbe converts that feedstock into biomass or products^1^. The CO_2_ can come from air or from a concentrated industrial or biogenic stream; both feed the same downstream process but impose different capture duties and carbon-accounting claims^2^. Formate is an attractive intermediate because it carries both carbon and reducing power. NAD-dependent formate dehydrogenase can oxidise it directly to CO_2_ and NADH; in the dark, formatotrophs must send part of that NADH through respiration to make ATP. A retinal rhodopsin pump could replace that respiratory ATP with light-derived proton motive force ^3–5^. The process-level question is therefore not only whether rhodopsin-pumped protons can regenerate ATP and improve biomass yield, but whether the saved formate is worth more than the light and cellular resources used to obtain it.

That trade-off spans three scales: the retinal absorption peak controls absorption, quantum yield and photocycle kinetics, while a carotenoid antenna can widen the absorption band by transferring energy to retinal, as in natural xanthorhodopsins, so the retinal peak and antenna band together set the action spectrum ^6^. In the cell, the value of ATP depends on the carbon pathway: the reductive glycine pathway (rGlyP) incorporates formate directly, whereas the Calvin–Benson–Bassham (CBB) cycle first oxidises it to CO_2_ ^7–9^. In the reactor, self-shading limits how much light reaches the culture ^10,11^. Rhodopsin can enhance bacterial CO_2_ fixation^12,13^, and engineering rGlyP into *C. necator* has increased biomass yield from formate ^14^. These studies do not show which pathway gains more from pumping, or whether spectral engineering still helps when the antenna competes with photovoltaics for light.

We ask how much formate light-derived ATP saves, why rGlyP and CBB respond differently, whether retinal and antenna tuning improve the whole process, how the CO_2_ source changes the result, and what limits efficiency. The model links wavelength-resolved rhodopsin photophysics, GECKO-style enzyme-constrained metabolism^15^ and two-flux reactor optics. We compare the pathways at equal growth in a chemostat operated at dilution rate *D* = *µ*, so the effect of light is measured by formate savings rather than faster growth.

The model gives a *conditional upper-bound* design map, not a plant forecast. The optimisation includes CO_2_ capture, formate synthesis, reactor light transport, oxygen and formate transfer, and a membrane-allocation cap, but excludes mixing, temperature control, product separation, collector spacing, weather, downtime and unmodelled growth burden. For the main direct-air-capture case, biomass energy is divided by all sunlight incident on the collector, including the share used for CO_2_ capture and formate synthesis, with no credit for unused energy (Fig. 1b). We apply the same boundary to a 14 wt% point-source case; other efficiency measures are labelled as diagnostics (Table S10).

**Figure 1:**
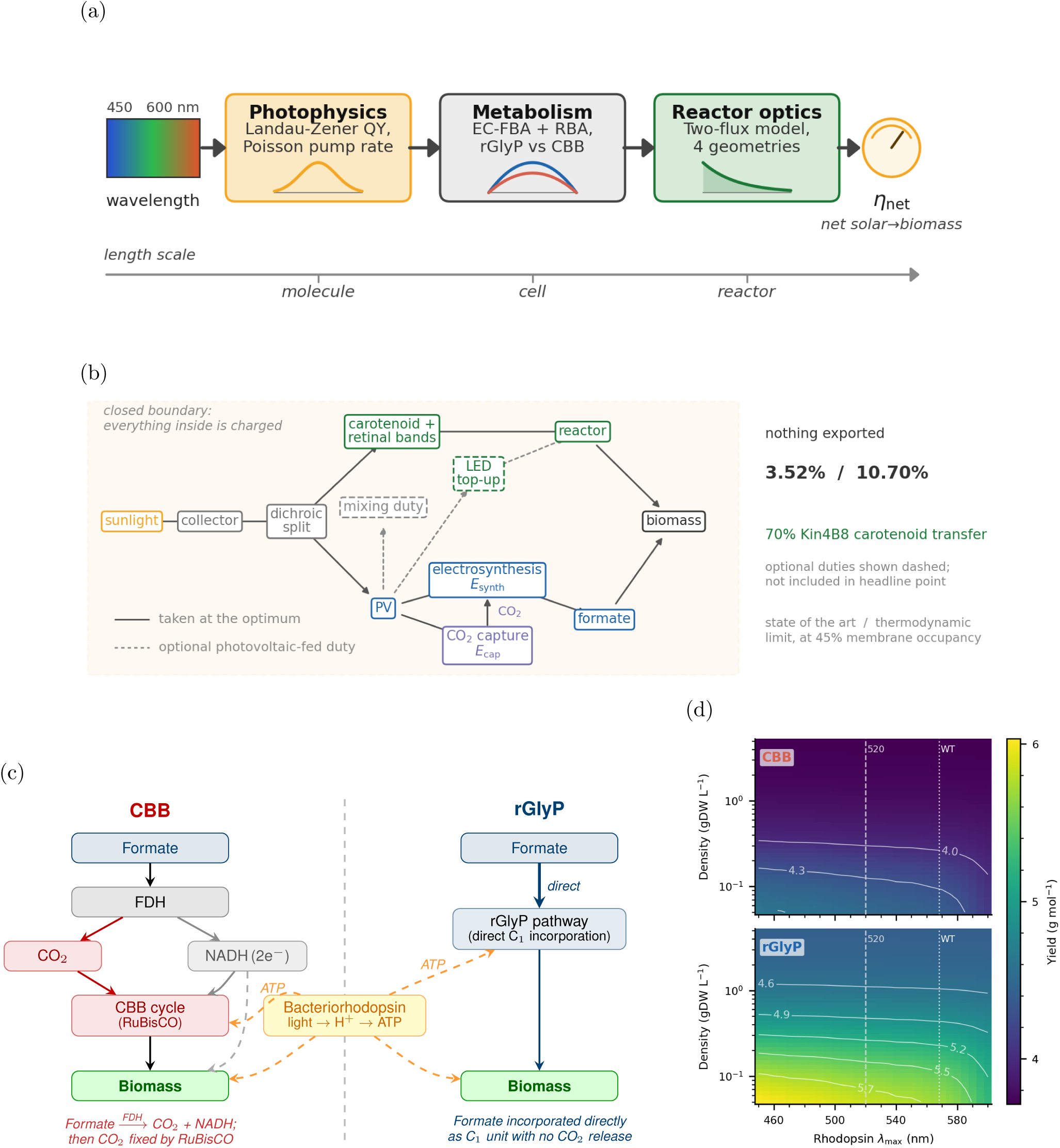
Multi-scale framework for rhodopsin-augmented formatotrophy. **(a)** Workflow linking photophysics, metabolism, reactor optics and *η*. **(b)** Closed energy boundary; values use 45% occupancy and the model-fit 70% Kin4B8 sensitivity, while dashed duties are excluded (Table S11). **(c)** CBB and rGlyP carbon routes coupled to rhodopsin-driven ATP synthesis. **(d)** Fixed-*µ* yield across *λ*_max_ and cell density at 100 PPFD (CBB upper, rGlyP lower); dotted, wild-type 568 nm; dashed, 520 nm reference.

Figure 1a connects the causal chain, while panel c shows where rGlyP and CBB differ. Quantum yield and photocycle kinetics convert local PPFD into a per-molecule pump rate; membrane-limited abundance converts that rate into proton flux; ATP synthase converts proton flux into ATP; and fixed-growth metabolism converts ATP into avoided formate oxidation. Because rhodopsin supplies neither carbon nor reducing power, any improvement must appear as less formate burned for energy, not as altered carbon assimilation.

Blue-shifted retinal absorption improves per-molecule pumping, but that advantage contracts as culture density rises because an optically thick culture already absorbs nearly all incident photons (Fig. 1d). Once photon energy, solar-band availability and reactor delivery are included, the molecular blue preference becomes a broad reactor-scale optimum that includes wild-type bacteriorhodopsin. An antenna can broaden this action spectrum while the retinal maximum is tuned within that broad optimum; the joint design succeeds only if the additional pumping repays the photovoltaic output displaced by collecting the antenna band. Per-cell and fixed-geometry efficiencies are used to diagnose molecular or optical mechanisms; all process comparisons use the closed, capture-charged boundary in Fig. 1b and Table S10. Details of the surface-hopping, master-equation, enzyme-constrained and two-flux calculations are given in Methods.

## Results

Unless stated otherwise, all comparisons use the fixed-growth formulation: biomass production is fixed at *µ* = 0.126 h*^−^*^1^, chosen as 70% of the measured 0.18 h*^−^*^1^ maximum formate-growth rate, and formate uptake is minimised. Growth is therefore imposed on the linear programme rather than emerging from it. The experimental counterpart is a chemostat at dilution rate *D* = *µ*. This matches productivity between pathways and makes formate saving the readout. Photon flux is reported on the basis appropriate to each scale: in the per-cell sections PPFD is the uniform flux every cell experiences, and once light propagation enters, surface and volume-averaged flux are named separately wherever both appear.

### Metabolic pathway architecture sets the dark culture baseline

rGlyP’s dark advantage comes from avoiding respiratory formate loss, not from incorporating more carbon into biomass: at the fixed-growth setpoint, CBB requires 34.1 rather than 28.9 mmol formate gDW*^−^*^1^ h*^−^*^1^ and generates 40.6 rather than 31.5 mmol respiratory ATP gDW*^−^*^1^ h*^−^*^1^. Carbon allocation to biomass is nearly unchanged; CBB instead oxidises more formate and consumes more O_2_ (Fig. 2a,b). NAD-dependent FDH produces NADH directly from formate, some of which passes through Complex I and the terminal oxidase in the dark to build the proton motive force used by ATP synthase; neither pathway makes ATP from formate by substrate-level phosphorylation, and CBB drives this route faster than rGlyP at matched growth (Fig. 2e,f). Near the predicted growth ceiling, the optimisation progressively selects the reconstruction’s quinone-linked FDH2 route, which bypasses Complex I. NADH16 then falls even while FDH2, terminal-oxidase and ATP-synthase rates continue to rise. Membrane-associated FDH activity has been measured in *C. necator*, but recent fitness data indicate that soluble FDH carries most formate-oxidation flux under the conditions tested^16,17^. The bend is therefore a conditional model prediction near the proteome limit, not an experimentally observed route switch. Rhodopsin later replaces part of the ATP supplied by these routes (Fig. 4c).

**Figure 2:**
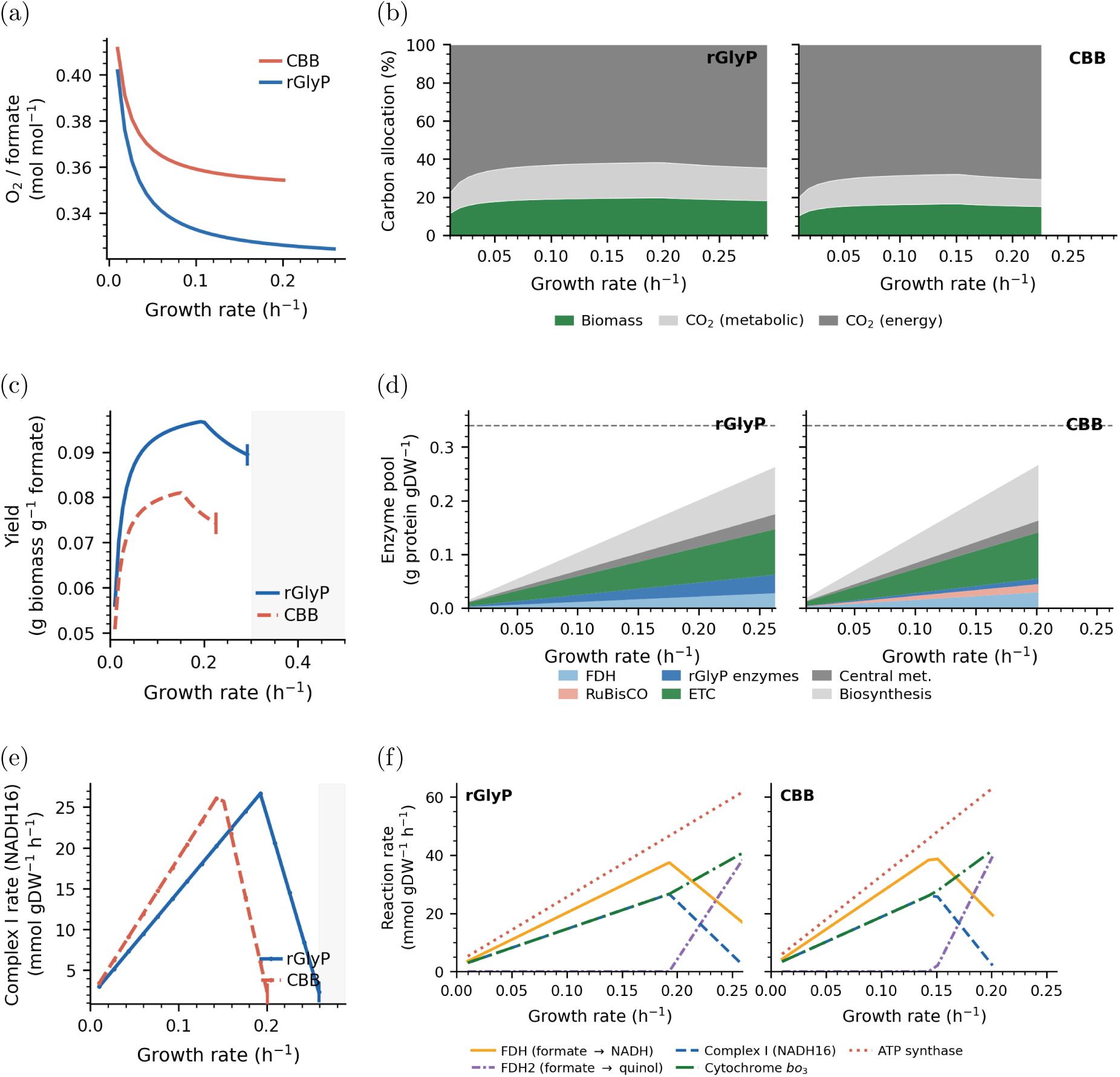
Dark formatotrophic baseline for rGlyP and CBB across growth rates. **(a)** O_2_:formate ratio. **(b)** Carbon allocation (rGlyP left, CBB right); CBB oxidises more formate through FDH. **(c)** Biomass yield; rGlyP remains higher across the feasible range. Maximum-growth endpoints are enzyme-cost-sensitive (Section S14). **(d)** Enzyme-pool allocation (rGlyP left, CBB right); dashed line, pool capacity. **(e)** Complex I (NADH16) flux. Its decline near the predicted growth ceiling marks the model-selected transfer to FDH2, not lower total respiration. **(f)** Rates of the key reactions connecting formate oxidation to ATP generation across the growth-rate sweep (rGlyP left, CBB right): NAD-dependent FDH, quinone-linked FDH2, Complex I, cytochrome *bo*_3_ oxidase and ATP synthase.

The model predicts 17.9% higher yield with rGlyP, close to the 16.5% measured by Dronsella et al. in chemostats ^14^. This agreement concerns the difference between pathways: absolute predicted yields are 3.5–4.7% low, and the parameters were not fitted to those data (Section S4). The yield and enzyme-pool profiles in Fig. 2c,d show that, across *µ* = 0.05–0.15 h*^−^*^1^, rGlyP uses 14.1–15.8% less formate and remains the lower-demand pathway under the tested enzyme-cost policies (Table S8; Section S14).

Because reducing-power shadow prices exceed the ATP shadow price in both pathways (Figure S12), a proton pump cannot remove the cofactor limitation or establish a higher growth ceiling; it can only replace ATP that would otherwise require respiratory formate oxidation.

### Rhodopsin proton pumping drives formate saving with pathway-dependent gain

At 100 PPFD, wild-type 568 nm and the same 45% membrane-occupancy cap used for the process reference, light-derived ATP reduces formate demand from 28.9 to 24.7 mmol gDW*^−^*^1^ h*^−^*^1^ for rGlyP and from 34.1 to 29.8 for CBB. Biomass yield rises 17% and 14%, respectively. These per-cell responses show formate preserved by pumping at a stated, membrane-feasible reference; they are not reactor optima (Table S26).

The larger rGlyP percentage does not mean that its pump is more tightly coupled to metabolism: one mmol of light-derived ATP saves about 0.57 mmol formate in either pathway because both use the same respiratory chain. rGlyP gains more in percentage terms only because it starts with lower formate demand.

Faradaic efficiency approaches the same carbon-only ceiling in both pathways, but rGlyP reaches it at lower irradiance because it has less respiratory ATP to replace (Fig. 3e). The Pareto fronts and fixed-formate growth and O_2_ responses are also consistent with light replacing respiration (Fig. 3a,b,c,d). The non-monotonic curve in panel f comes from a switch between growth- and non-growth-associated ATP demand, not a new pathway effect. rGlyP uses less formate at matched growth under every enzyme-constraint policy tested (Figure S13a).

**Figure 3:**
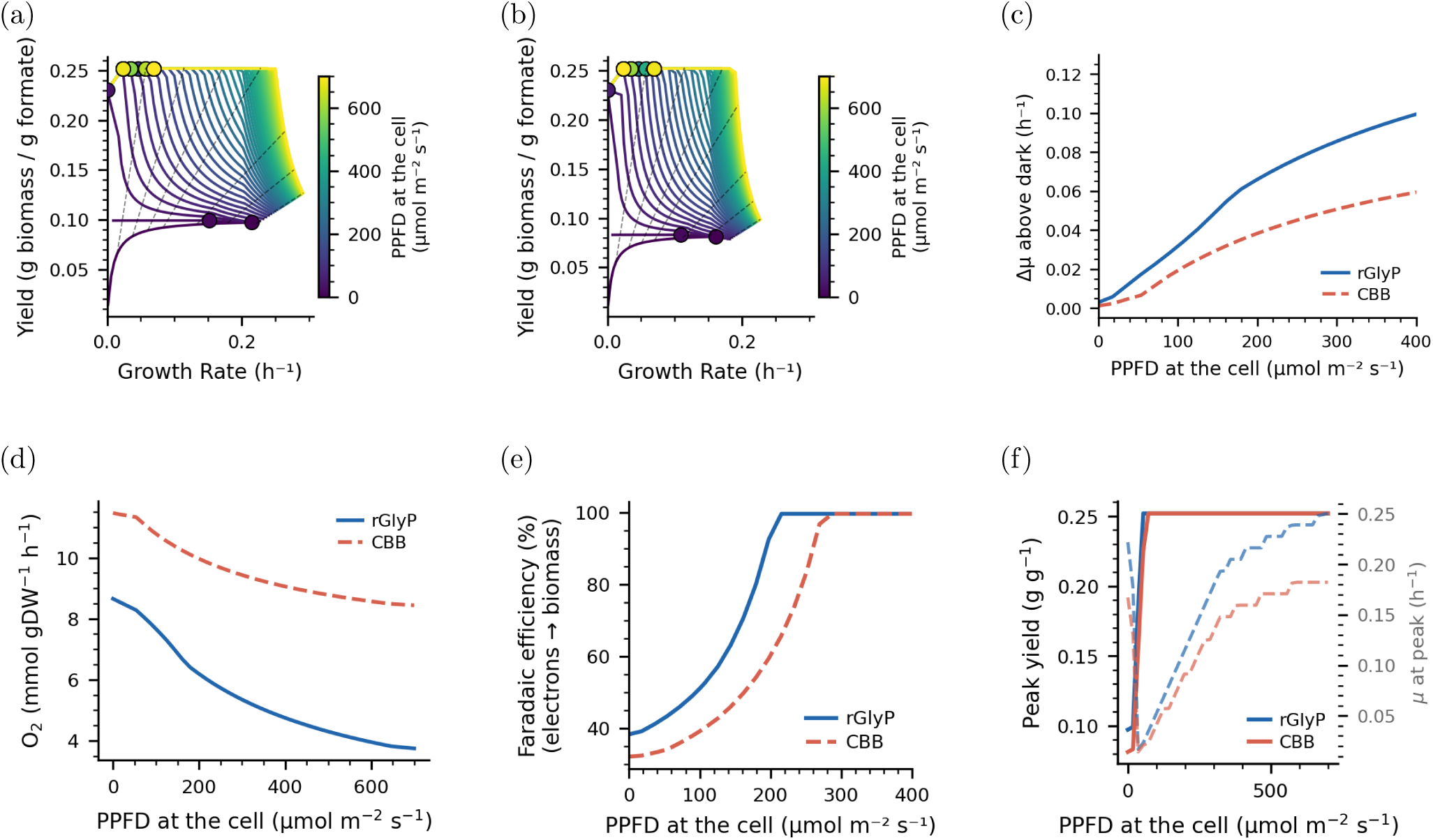
Fixed-growth formate saving under rhodopsin illumination (*µ* = 0.126 h*^−^*^1^). **(a,b)** Growth–yield Pareto fans showing illumination expanding the feasible frontier. **(c,d)** Growth and O_2_ response at fixed dark substrate. **(e)** Faradaic efficiency. **(f)** Highest growth rate within 1% of peak yield at formate *≥*2 mmol gDW*^−^*^1^ h*^−^*^1^. Curves use the full nominal membrane budget as a mechanistic envelope; the text reports the 45%-occupancy result at exactly 100 PPFD.

### Shadow prices and proteome allocation explain the fixed-growth rGlyP advantage

Light changes the energy source, not the carbon pathway: the yield and realised-pump curves trace both pathways towards the same carbon-only formate floor at high PPFD (Fig. 4a,b). Rhodopsin pumping feeds the same ATP-synthase step as dark respiration (Fig. 2f), so light replaces respiratory formate oxidation rather than opening a new pathway (Fig. 4c). rGlyP starts with lower formate demand and needs less light to remove respiration: about 215 PPFD, compared with 287 PPFD for CBB. The respiratory trace in Fig. 4d includes all native proton-exporting reactions, expressed at the model’s 4 H^+^ per ATP coupling. Steps in the ATP shadow price mark changes in the active constraints, not new biochemical interactions.

**Figure 4:**
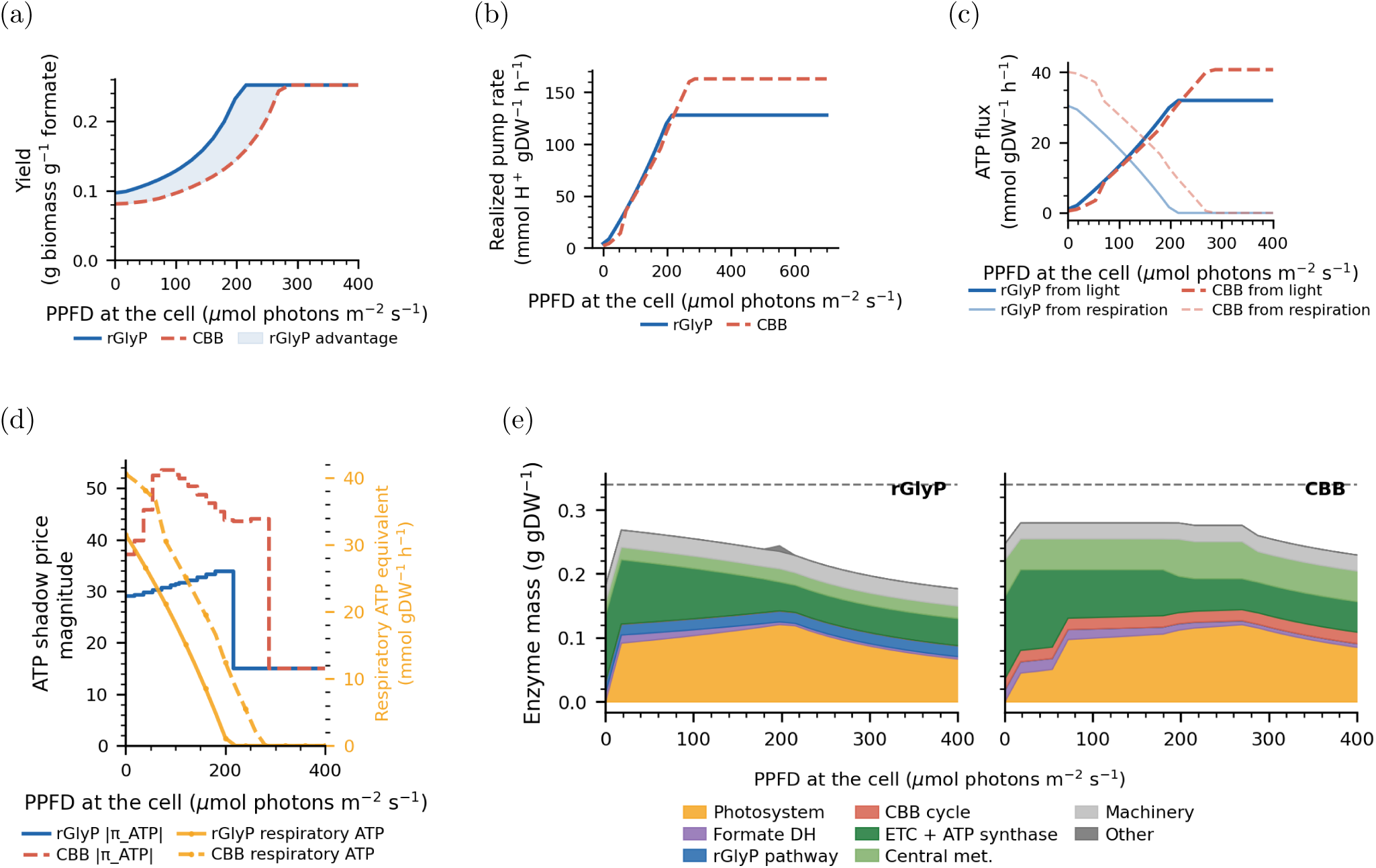
Constraint structure at fixed growth (*µ* = 0.126 h*^−^*^1^), full-membrane ceiling (Section S15). **(a)** Fixed-growth yield against PPFD. **(b)** Realised rhodopsin H^+^ pump flux. **(c)** ATP partitioning between rhodopsin and respiration. **(d)** ATP shadow price (left axis) and respiratory ATP-equivalent supply from all native proton-exporting reactions (right axis). Reducing-power shadow price vs. ATP shadow price is shown in Figure S12. **(e)** Proteome allocation, including both formate-dehydrogenase routes (rGlyP left, CBB right); “Other” contains unclassified enzyme draws; dashed line, pool capacity.

Eliminating respiratory flux does not eliminate the respiratory proteome: when light drives respiratory ATP production to zero, that enzyme sector falls by only *∼*4–5% because respiration and rhodopsin pumping share ATP synthase, while light adds a large photosystem sector (Fig. 4e). Its main benefit is therefore saved formate, not freed protein capacity.

Because membrane area becomes limiting before total protein mass, the full-membrane calculation in Fig. 4 is a mechanistic ceiling rather than the process assumption. Headline process efficiencies impose a 45% rhodopsin occupancy cap and report lower caps separately (Discussion; Section S15).

### Spectral tuning improves molecular pumping but not the process optimum

Figure 5a summarises the sequence-to-pump calculation. The molecular model predicts a modest blue preference: shorter-wavelength photons provide more thermodynamic headroom (Fig. 5b), and the posterior median quantum yield declines from 0.64 at 520 nm to 0.60 at wild-type 568 nm and 0.58 at 600 nm (Fig. 5c). In the model, added energy at the conical intersection increases branching toward the photoproduct state; together with photocycle kinetics, this raises protons pumped per photon toward shorter wavelengths (Fig. 5d, 5e). The wavelength ordering propagates to fixed-growth yield, but its effect is smaller than the rGlyP–CBB difference and disappears once either pathway reaches its carbon-only formate floor (Fig. 5f).

**Figure 5:**
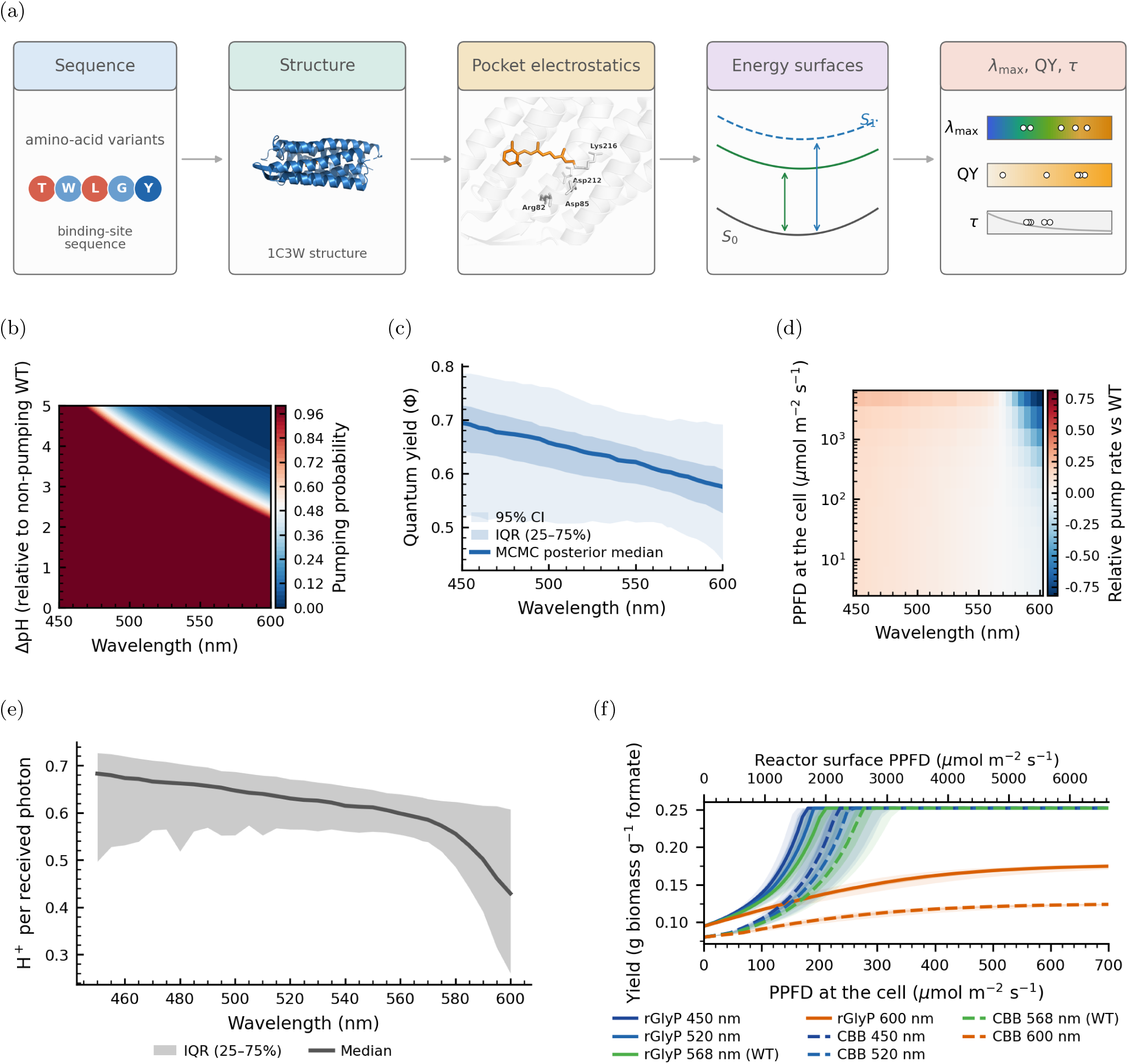
Wavelength dependence of quantum yield, proton output and fixed-growth yield. **(a)** Conceptual chain from binding-pocket electrostatics to *λ*_max_, QY and photocycle time; structure from PDB 1C3W and illustrative energy surfaces. **(b)** Thermodynamic feasibility at ΔΨ = 150 mV. **(c)** QY posterior against wave-length. **(d)** Relative pump rate against wavelength and PPFD, normalised to WT 568 nm. **(e)** H^+^ per photon at 100 PPFD. **(f)** Fixed-*µ* yield at four wavelengths; lower axis, per-cell flux, and upper axis, equivalent surface flux at *f*_del_ = 0.1055.

The only independent spectral check is Xie’s 110 K dataset, which was not used for calibration and also declines from blue to red^18^. The model overpredicts both the quantum yield and the size of this decline, while the experiment measures a cryogenic forward photoreaction rather than room-temperature pumping. It supports the direction, but not the size, of the predicted trend. Testing the magnitude requires a room-temperature action spectrum measured at several wavelengths (Section S9, Figure S14).

Higher molecular quantum yield does not create a narrow blue process optimum: after photon-energy weighting and reactor propagation, cell-scale efficiency stays within 0.05 percentage points of its maximum from 526 to 569 nm, and closed-boundary reactor efficiency does so from 509 to 568 nm. Because both ranges include wild-type bacteriorhodopsin, retinal peak tuning has little effect by itself; varying it with a carotenoid band tests whether extra pumping repays the photovoltaic output displaced by the wider band. This sensitivity represents transferred photons, without modelling a separate pigment in the two-flux calculation or charging antenna assembly.

### Reactor geometry changes light delivery and the pathway advantage

Reactor self-shading compresses spectral differences rather than selectively removing blue light. In the fixed-geometry sweep, delivery is almost wavelength-neutral because the culture is optically thick across the full range; changing 450 to 600 nm therefore changes delivered fraction far less than the molecular pump calculation suggests. When density, depth and PPFD are allowed to compensate during optimisation, the spectral response flattens further, so the molecular blue preference does not survive as a narrow reactor optimum.

Geometry matters mainly by redistributing irradiance: across the four configurations shown in Fig. 6a, external cylinders and annuli focus incoming light as the illuminated cross-section contracts, whereas the flat panel and internally illuminated bag do not (Fig. 6b). The density, surface-power and depth sweeps show the same trade-off (Fig. 6c,d,e), but do not reverse the metabolic conclusion: rGlyP remains more efficient than CBB across all four geometries (Fig. 6f). At high density the geometries converge because the electricity required to supply formate, rather than local light delivery, dominates the energy balance.

**Figure 6:**
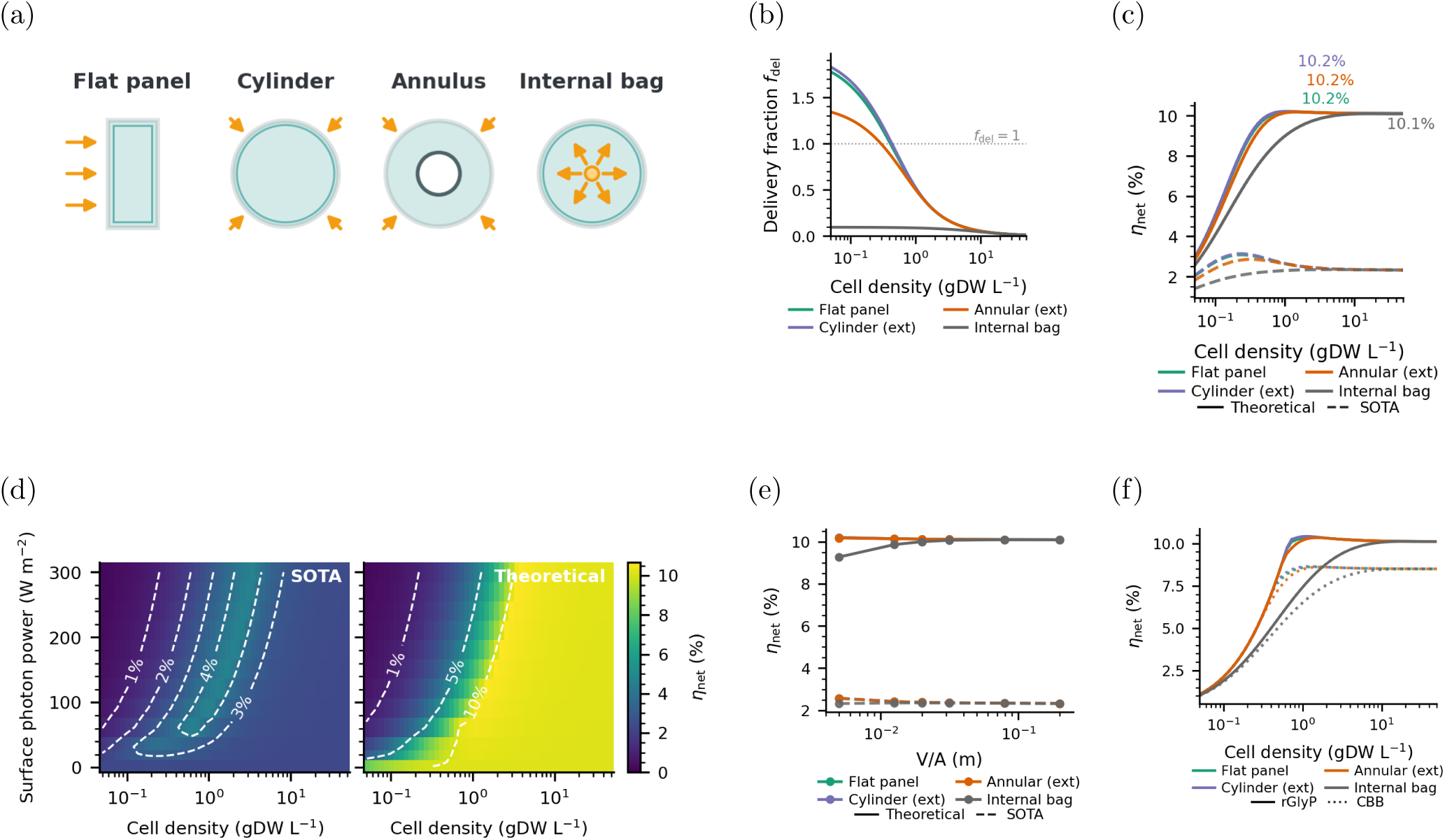
Diagnostic reactor-geometry sweep (*V/A* = 0.02 m). Solid, thermodynamic limit; dashed, state of the art. Values use the fixed-geometry diagnostic boundary (Table S10); panels show rGlyP unless stated. **(a)** Four geometries at common *V/A*. **(b)** Delivery fraction at 568 nm and 21 W m*^−^*^2^. **(c)** Efficiency against cell density. **(d)** Cylinder across density and surface power at 568 nm; white dashed, *η* contours. **(e)** *V/A* swept at 5 gDW L*^−^*^1^ and 21 W m*^−^*^2^. **(f)** rGlyP against CBB at 568 nm and 50 W m*^−^*^2^.

Figure 6 is a diagnostic comparison at fixed *V/A*, not the process optimisation. Its denominator counts in-band photons delivered to the culture plus the photovoltaic input for formate and CO_2_ capture, but not unused sunlight incident on a whole collector aperture. It therefore must not be compared numerically with the closed-boundary values in Fig. 7; Table S10 gives both definitions. Only its light-delivery trends, not its absolute efficiencies, carry into the closed-boundary optimisation.

**Figure 7:**
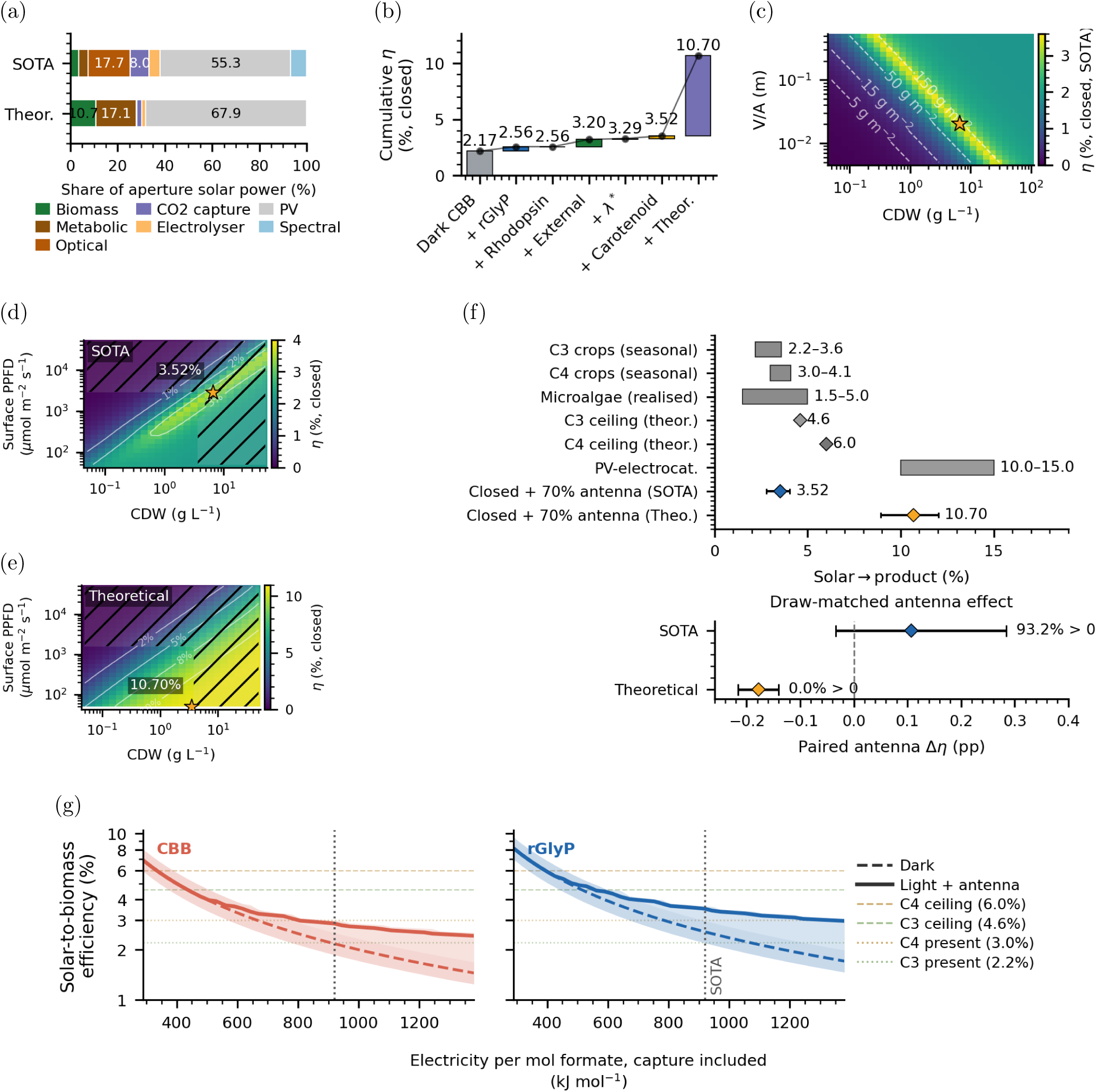
Optimised solar-to-biomass efficiency under membrane and reactor constraints. Values use *µ* = 0.126 h*^−^*^1^, the 45% membrane cap and the closed boundary. Panels use direct-air capture except for the sweep in panel g; except for labelled intermediates in panel b, they use the model-fit 70% carotenoid–retinal case. **(a)** Energy cascade. **(b)** Ordered lever decomposition ending with the re-optimised thermodynamic-limit antenna case; increments depend on order. Dark rungs use standard PV without a light aperture. **(c)** Closed-boundary *η* over density and depth; dashed lines, constant areal loading. **(d,e)** Current and thermodynamic-limit operating maps; hatching, constraint violations; stars, optima. **(f)** Literature benchmarks and the 70% Kin4B8 case ^1,21,22^; whiskers show propagated ranges and paired antenna effects; assembly burden is excluded. **(g)** Dark and light-enabled optima for CBB (left) and rGlyP (right) across formate electricity demand on a logarithmic efficiency axis; the vertical line marks SOTA, shading shows propagated 95% ranges, and coloured horizontal lines give seasonal C3/C4 conversion and theoretical ceilings ^21,22^. Light-enabled optima include the dark branch.

### Rhodopsin has process value when formate is costly

The best design uses rGlyP, external illumination and green absorption. CO_2_ capture accounts for about 46% of the electrical demand per mole of formate (Fig. 7a). At the 45% membrane cap, retinal alone gives 3.29% closed-boundary efficiency with current electrocatalysis. A Kin4B8-like band raises this to 3.38% at the measured *∼*50% transfer and 3.52% at the model-derived 70% estimate ^19,20^. Across matched uncertainty samples, the antenna adds a median 0.107 percentage points (95% range *−*0.034 to +0.284) and helps in 93.2% of samples, although the calculation excludes antenna assembly cost (Fig. 7f). Figure 7b shows the ordered steps from CBB to rGlyP, rhodopsin, external illumination, retinal tuning, the antenna and the re-optimised thermodynamic-limit process; the individual increments depend on their order.

The four re-optimised curves in Fig. 7g show that the value of light falls as formate becomes cheaper. They compare CBB and rGlyP with and without light while holding the other current-technology assumptions fixed; the light-enabled search can choose the dark design, so it never performs worse. Replacing the deployed-DAC assumption with CO_2_ separation from a 14 wt% source lowers electricity use from 919.9 to 529.3 kJ per mole formate, a 69% reduction in separation duty on the same minimum-work basis. In this point-source case, the 70% carotenoid–retinal design raises efficiency from 4.45% in the dark to 4.82%, an 8% gain; under deployed DAC, it rises from 2.56% to 3.52%, a 38% gain (Table S19) ^2^. At the thermodynamic limit, the standard-PV dark design gives 11.15%; imposing retinal or the antenna lowers this to 10.89% or 10.70%, so the light-enabled optimum retains the dark design. The operating maps show the feasible regions and bounded optima for both technology cases (Fig. 7d,e).

Every illuminated optimum reaches the imposed rhodopsin-occupancy limit, although the 45% cap has not been achieved in *C. necator* ; lower caps are reported in Table S27. Retinal-only efficiency spans 3.0–3.3% across the 30–45% caps. Relaxing any one transport limit tenfold changes efficiency by at most 0.05 percentage points because density and depth compensate along a broad ridge (Fig. 7c). A 6.7% biomass burden would erase the central antenna gain, and the required expression level, steady illumination and omitted plant duties remain untested.

## Discussion

Rhodopsin supplies ATP and reduces the formate burned for respiration, saving almost the same amount of formate per pump-derived ATP in rGlyP and CBB because both use the same respiratory chain. rGlyP shows the larger percentage yield gain because its dark demand is lower: at 100 PPFD and the 45% membrane cap, the model predicts gains of 17% for rGlyP and 14% for CBB.

Retinal tuning has little process effect because the wavelength optimum is broad and includes wild type. A carotenoid antenna helps under current electrocatalysis: its median paired gain is 0.107 percentage points and it helps in 93.2% of uncertainty samples. At the thermodynamic limit it lowers the imposed-light result in every sample because photovoltaics make better use of the same photons; a light-enabled plant would select the dark branch. The antenna also adds less than 0.01 points when CO_2_ comes from a 14 wt% point source. Both results have the same cause: light is most useful when formate is expensive to make. Our earlier study estimated capture equipment costs of £0.9–1.2 million for this flue-gas case and £1.7–2.6 million for DAC at 1 kt CO_2_ yr*^−^*^1^ ^2^. These are capture costs, not total plant costs, and point-source use depends on source lifetime, impurities and transport. Unless the source is biogenic or the carbon is stored, it recycles an emission rather than removing CO_2_ from the air.

The metabolic comparison is better supported than the spectral prediction: the predicted 17.9% rGlyP yield advantage is close to the measured 16.5%, and the ordering survives the tested enzyme-cost and allocation assumptions (Sections S14, S11, S13). The spectral model instead fits sixteen parameters to three wild-type measurements^23–25^. Withheld 110 K data match the direction but not the size of the wavelength trend, and the model misses the mutant ordering^18^. Room-temperature measurements of quantum yield, photocycle lifetime and proton flux across variants are needed.

Expression is the main construction risk because the 45% membrane cap has not been reached in *C. necator* and requires about 8% of cell protein for rhodopsin. Retinal-supplemented *E. coli* can assemble about 187,000 proteorhodopsin copies per cell^26^, but formate-grown *C. necator* has a smaller membrane sector ^27^. The model omits folding stress, incomplete retinal loading, inactive protein, collector spacing, weather, downtime, nutrients, recovery, most plant energy and antenna assembly. The reported optima lie on broad density–depth ridges and are not unique hardware designs.

Because the combined system has not been tested, a first experiment could compare rGlyP and CBB chemostats at *D* = 0.05 h*^−^*^1^ under dark and 100-PPFD, 568-nm conditions, with a non-pumping control. At the 45% cap, predicted formate demand falls from 12.50 to 8.27 mmol gDW*^−^*^1^ h*^−^*^1^ for rGlyP and from 14.55 to 10.32 for CBB. Measuring pump abundance, retinal loading and proton flux would distinguish poor expression from weak metabolic coupling. Reactor design and the cost of formate then require a separate engineering study ^1^.

## Resource Availability

Materials, data, and code requests should be directed to and will be fulfilled by the lead contact, Haris Saeed. This study generated no new biological materials. The data and complete analysis pipeline used in this study are publicly available at https://github.com/hs280/Formate_assimilation_rGlyPvsCBB_Cupriavidus_necator; specific file paths, dependent scripts, and software versions are listed in the Resource Availability and Key Resources Table subsections of STAR⋆ Methods below.

## Acknowledgements

H.S. acknowledges funding from the UK Research and Innovation Engineering and Physical Sciences Research Council Doctoral Training Partnership (grant EP/W524311/1). W.E.H. acknowledges support from the EPSRC EEBio programme grant (EP/Y014073/1) and the BBSRC UK–Japan programme for Engineering Biology (ISPF-234, APP38606, UKRI251).

## Author Contributions

H.S. conceived the study, developed the modelling framework, performed the simulations and analyses, prepared the figures, and wrote the initial manuscript. W.E.H. and A.Y. supervised the work and contributed to interpretation, framing and manuscript revision. All authors reviewed the manuscript.

## Declaration of Interests

The authors declare no competing interests.

## STAR⋆ Methods

### Key Resources Table

**Table 1:**
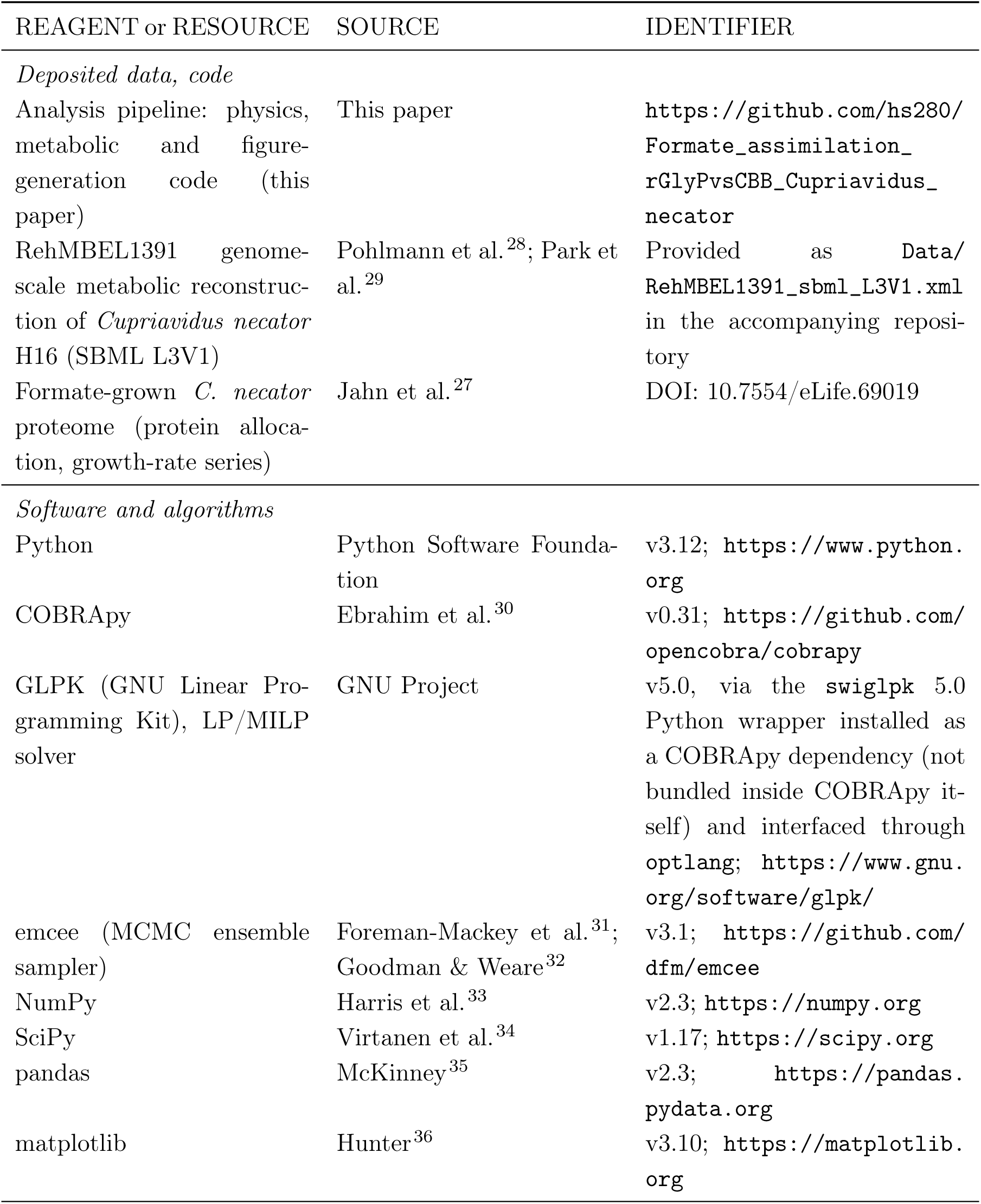
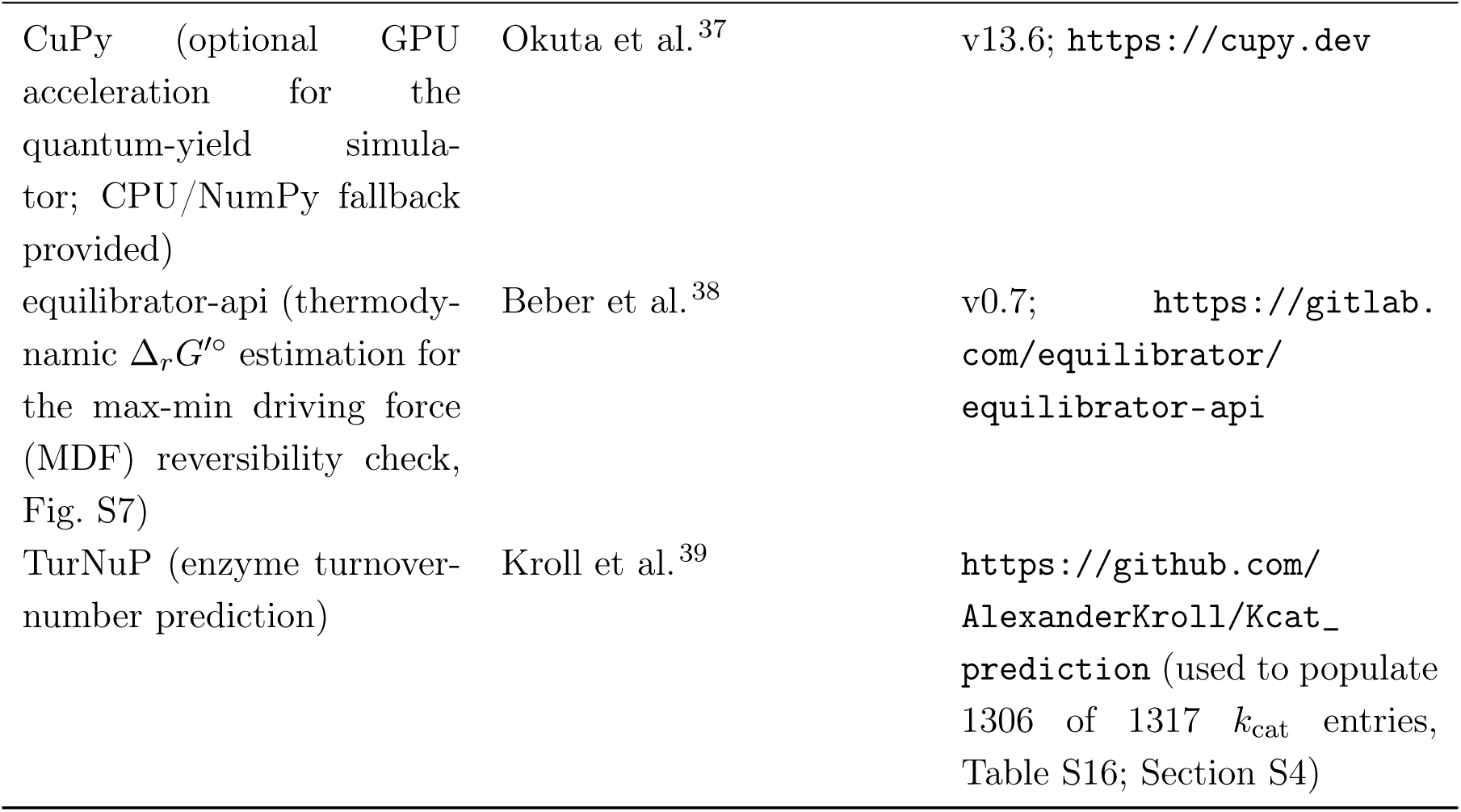
Key resources. Software, reference datasets and the deposited analysis pipeline used in this study.

| REAGENT or RESOURCE | SOURCE | IDENTIFIER |
| --- | --- | --- |
| *Deposited data, code* |  |  |
| Analysis pipeline: physics, metabolic and figure-generation code (this paper) | This paper | https://github.com/hs280/Formate_assimilation_rGlyPvsCBB_Cupriavidus_necator |
| RehMBEL1391 genome-scale metabolic reconstruction of *Cupriavidus necator* H16 (SBML L3V1) | Pohlmann et al.<sup>28</sup>; Park et al.<sup>29</sup> | Provided as Data/RehMBEL1391_sbml_L3V1.xml in the accompanying repository |
| Formate-grown *C. necator* proteome (protein allocation, growth-rate series) | Jahn et al.<sup>27</sup> | DOI: 10.7554/eLife.69019 |
| *Software and algorithms* |  |  |
| Python | Python Software Foundation | v3.12; https://www.python.org |
| COBRApy | Ebrahim et al.<sup>30</sup> | v0.31; https://github.com/opencobra/cobrapy |
| GLPK (GNU Linear Programming Kit), LP/MILP solver | GNU Project | v5.0, via the swiglpk 5.0 Python wrapper installed as a COBRApy dependency (not bundled inside COBRApy itself) and interfaced through optlang; https://www.gnu.org/software/glpk/ |
| emcee (MCMC ensemble sampler) | Foreman-Mackey et al.<sup>31</sup>; Goodman & Weare<sup>32</sup> | v3.1; https://github.com/dfm/emcee |
| NumPy | Harris et al.<sup>33</sup> | v2.3; https://numpy.org |
| SciPy | Virtanen et al.<sup>34</sup> | v1.17; https://scipy.org |
| pandas | McKinney<sup>35</sup> | v2.3; https://pandas.pydata.org |
| matplotlib | Hunter<sup>36</sup> | v3.10; https://matplotlib.org |
| CuPy (optional GPU acceleration for the quantum-yield simulator; CPU/NumPy fallback provided) | Okuta et al.<sup>37</sup> | v13.6; https://cupy.dev |
| equilibrator-api (thermodynamic $\Delta_r G'^o$ estimation for the max-min driving force (MDF) reversibility check, Fig. S7) | Beber et al.<sup>38</sup> | v0.7; https://gitlab.com/equilibrator/equilibrator-api |
| TurNuP (enzyme turnover-number prediction) | Kroll et al.<sup>39</sup> | https://github.com/AlexanderKroll/Kcat_prediction (used to populate 1306 of 1317 $k_{cat}$ entries, Table S16; Section S4) |

### Resource Availability

#### Lead Contact

Further information and requests should be directed to and will be fulfilled by the lead contact for day-to-day materials, data and code requests, Haris Saeed, one of the two co-corresponding authors listed on the title page.

#### Materials Availability

This study is purely computational and generated no new biological materials, reagents, or organisms.

#### Data and Code Availability

The public repository at https://github.com/hs280/Formate_assimilation_rGlyPvsCBB_Cupriavidus_necator contains the SBML metabolic model and experimental inputs in `Data/`, analysis code in `Bin/` and `PaperScripts/`, and the numerical outputs underlying the headline process results in `Results/reactor_geometry_wavelength/`. The main reproducibility entry point is `PaperScripts/reproduce_all.py`; figure panels are regenerated by this pipeline and are embedded in the supplied manuscript PDFs. The deposited headline outputs include the closed-boundary optima, retinal-only and joint-carotenoid uncertainty summaries and their per-draw tables, carotenoid-band sensitivity, and the matched carbon-supply comparison. In particular, `closed_boundary_uncertainty_draws_cap45_dac_global_hhv_t.csv` and `carotenoid_uncertainty_draws_joint_antenna_cap45_dac.csv` support the paired antenna analysis. This URL provides public access; a versioned archival record can be cited separately when available. Additional information is available from the lead contact on request.

### Experimental Model and Study Participant Details

Not applicable. This study is purely computational: it models the physiology of *Cupriavidus necator* H16 using a genome-scale reconstruction and literature-derived parameters, and involves no new wet-laboratory experiments, live organisms, cell lines, or study participants of our own. This section is therefore omitted per Cell Press guidance for computational studies.

### Method Details

#### Integrated computational model

We coupled photophysics and quantum-yield prediction, photocycle kinetics, and enzyme-constrained metabolic optimisation under proteome and membrane constraints. Each condition specifies the assimilation pathway (rGlyP or CBB), illumination and environmental constraints. Derivations, identifiability and parameterisation are described in the Supplemental Methods (Sections S1–S8). Thermodynamic checks cover the proton-motive-force (PMF) subnetwork containing formate dehydrogenase, terminal oxidases, transhydrogenase and ATP synthase. These checks constrain the direction of the reactions through which rhodopsin displaces respiratory ATP production; they do not establish concentration-resolved feasibility of every reaction in the genome-scale model.

#### Headline assumptions for the process-efficiency calculation

All *η* claims depend on the parameter set in Table S22; the two electrolyser scenarios bracket present (“SOTA”) and near-thermodynamic (“Theoretical”) CO_2_-to-formate windows.

The model splits incident sunlight between a rhodopsin absorption band and a photovoltaic branch. The dichroic transmits a selectable fraction of the absorption band to the culture with an efficiency of 0.90; the remaining light supplies photovoltaics for formate electrosynthesis, CO_2_ capture and any LED top-up. Culture attenuation is evaluated after light reaches the reactor surface: at the reported optimum, mean cellular irradiance is approximately 10% of surface irradiance. The headline efficiency charges all sunlight incident on the collector, including conversion and optical losses, without credit for unused light or exported energy (Table S11). Concentration can require a collector larger than the reactor. The resulting metric is biomass energy per incident collector solar energy at the design irradiance; it does not include the additional land required for plant spacing or infrastructure. The calculation is implemented in `PaperScripts/reactor_geometry/data/optimize_hybrid_dichroic_led.py`.

#### Physics layer I: activation and quantum yield

Photon arrivals at a single molecule follow Poisson statistics with rate *k*_abs_ = PPFD *× N*_A_ *×* 10*^−^*^6^ *× σ*_eff_. The physical single-chromophore cross-section is *σ*_0_ = *ε* ln 10 *×* 10*^−^*^1^*/N*_A_ *≈* 2.41 *×* 10*^−^*^20^ m^2^, from the bacteriorhodopsin extinction coefficient *ε* = 63,000 M*^−^*^1^ cm*^−^*^1^ ^40^. The pump conversion uses an inherited lumped scale, *σ*_eff_ = 4.5*σ*_0_. Its original independent calibration datum could not be traced; at the 100-PPFD wild-type reference it changes the calculated rate from 0.87 to 3.85 H^+^ s*^−^*^1^ per molecule. We therefore treat it as an uncertain model scale, not a measured or spectroscopic cross-section (Section S6.8). Self-shading uses *σ*_0_. Each photon triggers productive isomerisation with probability Φ_QY_, giving a Poisson mean *k*_abs_Φ_QY_*τ* per photocycle and

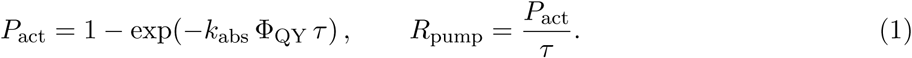

Proton stoichiometry is 1 H^+^ per completed photocycle^24^, and the action-spectrum quantum efficiency for pumping in intact cells matches Φ_QY_ ^41^, so no additional prefactor is used beyond the fixed *σ*_eff_ */σ*_0_ = 4.5 scale. In the low-activation limit, *k*_abs_Φ_QY_*τ ≪* 1, pump rate is approximately proportional to *σ*_eff_. A shared multiplier then cancels approximately from ratios of per-molecule pump rates evaluated within that limit. It does not generally cancel from the nonlinear pump equation, pathway growth or yield ratios, or optimised process efficiencies. Sensitivity of the headline efficiency to this calibration is evaluated separately (Section S6.8).

Φ_QY_ (all-*trans →* 13-*cis* isomerisation per absorbed photon) was predicted from a one-dimensional torsional potential-energy-surface model. Excited-state (S_1_) dynamics were propagated as coherent ballistic trajectories, reflecting the sub-200 fs ballistic wavepacket motion observed for retinal on S_1_ ^42,43^. Langevin friction and thermal noise (300 K) apply only after transfer to S_0_. Mass and friction were perturbed per trajectory to represent environmental heterogeneity. Hopping follows a Landau–Zener surface-hopping scheme without propagated electronic amplitudes, so it is neither fewest-switches surface hopping nor the fuller Zhu–Nakamura treatment^44,45^ (Section S1). The velocity is rescaled to conserve energy on a successful hop, and forbidden hops are rejected as frustrated, *P*_hop_ = exp(*−πV*^2^*/*2ħ*|v|α*), with *V* the diabatic coupling at the crossing, *v* the nuclear velocity at the seam and *α* the diabatic slope difference. The reduction is a deliberate trade against full multireference QM/MM dynamics (Section S1). Four parameters were inferred by MCMC: reduced mass *m*, ground-state friction *γ*, diabatic coupling scale *c_s_* and conical-intersection gap Δ*E*_CI_. A fifth, the CI-shift fraction *β ∈* [0, 1], has zero likelihood contribution at the wild-type wavelength and was therefore sampled from its Uniform(0, 1) prior during wavelength predictions. Sampling used emcee ^31,32^, calibrated against the wild-type quantum yield (Φ_QY_ = 0.64, *σ*_QY_ = 0.04) only, with no multi-mutant spectral data in the likelihood; sampler settings and convergence diagnostics are in Section S2. Posterior-predictive QY at 568 nm is 0.60 *±* 0.05.

#### Physics layer II: photocycle kinetics and feasibility

Photocycle lifetime *τ* (*λ*) comes from a seven-state master equation with Marcus barriers and Grote–Hynes transmission coefficients, calibrated to *τ_M_ ≈* 5 ms and *τ*_total_ *≈* 15 ms for wild-type bacteriorhodopsin; the equations and 12 parameters are in Section S3. At 520–568 nm and 100 PPFD the Poisson saturation parameter *x* = *σ*_eff_ QY *τ I ≈* 0.031–0.051, so *R*_pump_ *≈ σ*_eff_ QY *I* to within 2.5% and *τ* enters only as a weak second-order modulator (*|*d ln *R/*d ln *τ | <* 2.6%); the dominant wavelength dependence is through QY. At the 600 nm endpoint the longer *τ* pushes *x ≈* 0.6, and partial saturation reduces the linear-regime estimate by *∼*25%. Thermodynamic feasibility compares usable photon energy against proton electrochemical demand, with *E*_usable_ = max(0*, γ hc/λ −* 12 *k_B_T*), where *γ ≈* 0.284 is the K-state electrostatic storage fraction (from the 0.62 eV K-intermediate energy at 568 nm ^46^) and 12 *k_B_T ≈* 0.31 eV the minimum dissipation for unidirectional cycling. Against Δ*µ_H_*_+_ = *e* ΔΨ + 2.303 *k_B_T* ΔpH, pumping is feasible (*P*_pump_ = 1) when *E*_usable_ *≥* Δ*µ_H_*_+_ and is otherwise penalised by *P*_pump_ = exp[*−*(Δ*µ_H_*_+_ *− E*_usable_)*/k_B_T* ].

#### Metabolic layer: enzyme-constrained flux balance analysis (EC-FBA) with resource balance allocation (RBA) and coupled photosynthesis

Following standard constraint-based practice^47–49^, the metabolic model is a linear programme on the *C. necator* H16 reconstruction RehMBEL1391^28,29^ (1538 reactions, 1348 metabolites, 1350 genes):

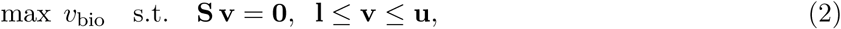

with **v** in mmol gDW*^−^*^1^ h*^−^*^1^. The biomass reaction embeds a growth-associated maintenance cost GAM = 135.0 mmol ATP gDW*^−^*^1^ of biomass formed; non-growth-associated maintenance is fixed at NGAM = 3.0 mmol ATP gDW*^−^*^1^ h*^−^*^1^ regardless of growth rate, within reviewed bacterial ranges^50^. Neither term is fitted here; both are the values reported for formate-grown *C. necator* by Jahn et al. ^27^ and are the same pair Dronsella et al. adopted for this same RehMBEL1391 reconstruction ^14^, so the maintenance costs are held fixed across every scenario rather than tuned against any yield reported here. The base model was extended with pathway-specific formate assimilation (rGlyP or CBB, one active per simulation^9^), a rhodopsin proton pump 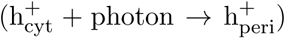, a passive proton leak (1.0 mmol gDW*^−^*^1^ h*^−^*^1^), and optional ectoine and trehalose routes retained but not exercised here.

Enzyme-capacity constraints follow GECKO (GEnome-scale model with Constraints based on Kinetic and Omics data) ^15^: *v_j_ ≤ σ_j_ k*_cat*,j*_ *E_j_*, where *σ_j_ ∈* (0, 1] is a saturation factor capturing the ratio of *in vivo* to *in vitro* turnover ^51^. Turnover numbers and molecular weights form a database of 1317 enzyme/reaction entries (Table S16), of which 1306 are TurNuP predictions^39^ keyed to H16 genes through the model gene–protein–reaction associations; reactions not covered are left unconstrained by the proteome pool. The three heterologous rGlyP enzymes absent from the host have no TurNuP prediction and were curated individually from measured homologues (Section S4). Reaction-level complex-cost overrides are applied before gene-level mapping for FDH, FDH2, Complex I, cytochrome *bo*_3_ and F_1_F_0_ ATP synthase, preventing multi-subunit respiratory enzymes from being left free or costed as single subunits. Reversible reactions were split into half-reactions with independent draws, reverse *k*_cat_ set to 0.1 *× k*_cat,fwd_ as a uniform Haldane approximation^52^, except where directionality enforces thermodynamic feasibility (Section S8). A further 54 reactions were constrained to zero or given directional bounds for formate-only aerobic growth (Table S15). Among them, the soluble NAD(P) transhydrogenase is held at zero flux so that rhodopsin-derived proton-motive force cannot be converted implicitly into NADPH; proton-coupled transhydrogenase is retained but treated as PMF-coupled rather than as a free NADPH source, and reopening the soluble route leaves the dark pathway advantage and the rGlyP formate demand unchanged (Section S14). Draws aggregate into one proteome pool, Σ*_j_* (MW*_j_/σ_j_k*_cat*,j*_) *v_j_≤ P*_total_ *f*_enzyme,eff_ (*µ*), in which the available enzyme fraction is reduced by a growth-rate-dependent ribosome sector (Eq. S29), with *P*_total_ = 0.68 g gDW*^−^*^1^ and *f*_enzyme_ = 0.50. These were calibrated against the formate-grown wild-type proteome of Jahn et al. ^27^: measured total protein set *P*_total_, mapping model-associated plus ribosomal proteins supported *f*_enzyme_ = 0.50, and ordinary least squares across five formate growth rates gave *f*_ribo_(*µ*) = 0.068 + 0.157 *µ* (capped at *f*_ribo,max_ = 0.40). These measurements calibrate the aggregate native partition and were not imposed as reaction-level abundances for the heterologous rGlyP or rhodopsin sectors, whose allocations remain optimisation variables. In fixed-growth runs the imposed *µ* sets the pool bound directly; in free-growth runs growth and pool are mutually dependent, so LP optimisation alternates with an update of *f*_enzyme,eff_ (*µ*) until convergence. Membrane-embedded enzymes additionally compete for finite inner-membrane area,

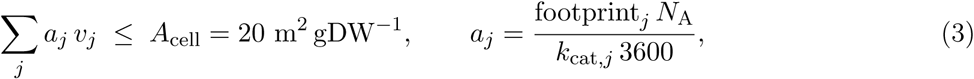

the area draw built analogously to the proteome draw so that a per-molecule cross-section becomes an area cost per unit flux. Both sectors that occupy the inner membrane are priced. The light-harvesting modules take *∼*7 nm^2^ per rhodopsin monomer and *∼*1000 nm^2^ per chlorophyll photosystem complex. The integral-membrane respiratory complexes that convert the pump’s proton-motive force take cross-sections estimated from published structures: *∼*22 nm^2^ for the F_o_ domain of F_1_F_o_ ATP synthase (a *c*-ring of *≈*5 nm diameter plus the *a*-subunit), *∼*70 nm^2^ for the elongated membrane arm of Complex I (*≈*18 *×* 4 nm), *∼*25 and *∼*30 nm^2^ for the *bd* and *bo*_3_ terminal oxidases, *∼*50 nm^2^ for quinone-linked formate dehydrogenase, and *∼*25 nm^2^ for the Complex II membrane anchor ^53–58^. The reconstruction’s FDH2 reaction has no gene association, so it is charged as a 170-kDa FdnGHI monomer rather than left free; lacking a host-specific turnover measurement, it uses a 200 s*^−^*^1^ proxy equal to the curated NAD-dependent FDH value. Membrane-associated FDH activity in *C. necator* supports retaining the route, but neither its physiological ubiquinone turnover nor the predicted high-growth switch has been measured ^16,17^. Cytochrome *bo*_3_ is charged as the complete 144-kDa four-subunit monomer at 200 s*^−^*^1^, the lower bound implied by quinol exchange at least once per 5 ms ^59^. These proxy costs prevent missing or incomplete gene mappings from favouring a respiratory route solely because its enzyme is absent from the reconstruction. Both protein and membrane occupancy must be priced for the constraint to be meaningful: the modules that *use* the proton-motive force have to compete for the same area as the module that makes it, so the ceiling arbitrates between pump and respiration rather than merely bounding pump abundance. Soluble enzymes and the transporters are left unpriced, the latter because the reconstruction does not resolve their aggregate occupancy, which makes the budget permissive rather than strict. The respiratory footprints are order-of-magnitude estimates, so their level is varied over a decade in Section S15. The budget is taken from the membrane proteome inventory of Jahn et al. ^27^ and is consistent with the membrane-economy budget of Zhuang et al. ^60^. Saturation factors are assigned by EC class, approximating typical *in vivo* levels ^51^ (values in Section S4). Four saturation policies were evaluated: unconstrained; uniformly saturated (*σ_j_* = 1); uniform 50% occupancy (*σ_j_* = 0.5); and the EC-class-specific multipliers used throughout, which are primary because they reproduce the measured proteome partition most closely but are not individually calibrated per reaction. The pathway ranking is invariant across all four (Section S4.8; Figure S13a).

#### Respiratory proton stoichiometry

Because the light module supplies protons, what a light-derived ATP is worth depends entirely on how many protons respiration would otherwise have pumped to make it: at fixed growth the formate displaced per proton is exactly 1*/*(H^+^ per formate oxidised). The RehM-BEL1391 reconstruction contains only the quinol-oxidase branches, cytochrome *bo*_3_ (4 H^+^ per quinol) and cytochrome *bd* (2 H^+^), and no cytochrome *c* carrier, *bc*_1_ reductase or *cbb*_3_ oxidase, so its most efficient route is FDH *→* Complex I *→ bo*_3_ at 7 H^+^ per formate. This stoichiometry is used throughout and required one correction to the implementation: the ATP-synthase proton budget must count every reaction that delivers H^+^ to the periplasm, whereas the earlier formulation credited only *bo*_3_ and formate dehydrogenase and left Complex I, the second NADH dehydrogenase and cytochrome *bd* uncounted although all three pump into the same periplasmic pool. That omission drove Complex I to zero flux and capped respiration at 6 H^+^ per formate; with the budget corrected the chain runs at the 7 H^+^ the reconstruction allows.

The reconstruction’s ceiling is not the organism’s. *C. necator* carries nine respiratory complexes, and its cytochrome *c* branch translocates 6 H^+^ per quinol against 4 and 2 for the quinol oxidases, with the high-O_2_-affinity *cbb*_3_ complex showing its strongest fitness requirement specifically during formatotrophic growth ^17^; a chain carrying that branch would reach 9 H^+^ per formate. We keep the published stoichiometry rather than adding reactions the reconstruction does not contain, and record which way that choice cuts: since the formate displaced per proton is 1*/*(H^+^ per formate), a 9 H^+^ chain would scale the light-driven formate saving by roughly 7*/*9, so the saving reported here is an upper bound with respect to respiratory efficiency. Because stoichiometry and GAM are not separately identifiable from a yield measurement, GAM is held at its literature value rather than refit to absorb this ceiling; the residual disagreement with measured yields is reported in Section S12.

Rhodopsin is represented as a transmembrane proton pump bounded by *R*_pump_ (Eq. 1); native ATP synthase converts the resulting PMF into ATP, with external proton-uptake shortcuts closed. The GECKO entry has MW = 27 kDa and a 7 nm^2^ footprint, so it draws from both proteome and membrane budgets. Its *k*_cat_ equals the calculated per-molecule pump rate. At the 100-PPFD wild-type reference the model gives *≈*3.85 H^+^ s*^−^*^1^ per molecule, subject to the unresolved 4.5*×* scale above. Shadow prices were taken from the primary GLPK solution; they report marginal growth value in free-growth runs and marginal substrate-saving value in fixed-growth runs. Solver tolerances and fixed-point settings are in Table S14.

#### Reactor-level optical model

Light propagation uses the two-flux (Schuster) approximation ^10^, tracking forward and backward irradiance and decomposing suspension optics into mass-specific absorption *a*(*λ*) = *κ*_abs,bg_ + *κ*_abs,rho_ and scattering *s*(*λ*) = *κ*_scat_. Scattering follows a literature-based power law^11,61^; in the absence of an H16 visible-absorption spectrum, the production calculation uses a flat background absorption of 0.02 m^2^ gDW*^−^*^1^ (Eqs. S33, S34). Rhodopsin self-shading uses a Gaussian centred at the variant’s own *λ*_max_ (FWHM 100 nm) with abundance taken from the EC-FBA allocation rather than imposed. The actinic flux at depth *z* under collimated front illumination, *G*(*z, λ*) = *I*^+^ + *I^−^*, is the Cornet–Schuster two-flux solution derived in Section S5, with attenuation set by 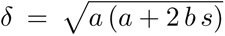, small-backscatter coefficient *b ≈* 0.01^11^ and biomass concentration *X*. Only the backscattered fraction is lost from the forward beam, so *δ ≪ a* + *s* (Figure S9a,b). Each depth slice drives the Poisson pump model to a local rate, converted to mmol H^+^ gDW*^−^*^1^ h*^−^*^1^ and mapped to local fixed-growth yield by interpolation from the metabolic sweep. Volume averages use trapezoidal integration over 200 slices, while the perfect-mixing limit instead evaluates the metabolic model at the volume-averaged irradiance. Radial geometries reduce to the same ODE under a power transformation, leaving a 1*/r* factor on recovery, so external illumination focuses irradiance with depth and internal illumination spreads it; the four cylindrical and annular configurations and their radii are in Section S5. A broad optical-input audit preserves the pathway and geometry conclusions but shows that wavelength-dependent host absorption can erase the small 520-nm advantage; spectral claims are therefore made for the broad green optimum rather than a resolved blue target (Section S5).

#### Parameter calibration across model layers

The model layers were parameterised separately, with no joint fitting objective. The four quantum-yield parameters were inferred by MCMC from wild-type yield (Φ_QY_ = 0.64 at 568 nm), and the twelve photocycle parameters from wild-type timing (*τ_M_ ≈* 5 ms, *τ*_total_ *≈* 15 ms). Their identifiability limits are described in Section S2. In the metabolic layer, GAM and NGAM were adopted unchanged from the literature, EC-class saturation factors were assigned from biochemical estimates, and total protein and the growth-dependent ribosome fraction were calibrated to Jahn et al.^27^. Proteome mapping supported the retained enzyme-accessible fraction of 0.50. Grunwald et al. ^62^ supplied the growth-rate reference used to choose operating conditions; their formate yield was compared with model predictions without refitting the maintenance or saturation parameters in this analysis. These yield comparisons are conditional on the literature-derived maintenance terms, native-proteome calibration and reconstruction assumptions (Section S12). Quantum-yield posteriors give wavelength-dependent credible bands; metabolic parameters are fixed point estimates with sensitivity checks across saturation policies (Section S4.8).

#### Computational operating modes and outputs

Three modes differ in which variable is fixed. The *operating-envelope sweep* grids formate uptake against photon flux (ranges in Table S14), fixing uptake and maximising *v*_bio_ (Eq. 2) to map the growth–yield envelope. *Fixed-growth formate minimisation* fixes *v*_bio_ at 0.126 h*^−^*^1^ (70% of the observed 0.18 h*^−^*^1^ formate-growth reference) and minimises formate uptake while sweeping per-cell PPFD; this is the primary mode and defines the yield, formate-saving, constraint-topology, wavelength and reactor results. *Fixed-dark-formate growth response* fixes formate at the dark reference from a separate dark calculation with photosynthesis disabled and maximises *v*_bio_ across a pump sweep; because each pathway’s formate level is set independently, these are within-pathway light responses. Outputs include growth rate, yield, ATP fluxes from rhodopsin and respiration, O_2_ consumption and shadow prices. Metabolic outputs are deterministic LP optima over the 450–600 nm analysis range. The quantum-yield posterior is over physical parameters rather than over wavelength, so it predicts across the whole band; its credible interval is narrowest at the 568 nm calibration anchor and widens toward both ends, quantified together with the full 450–700 nm sweep in Section S7 and Figure S4. The diagnostic efficiency reported in Figures S5a and S5b is

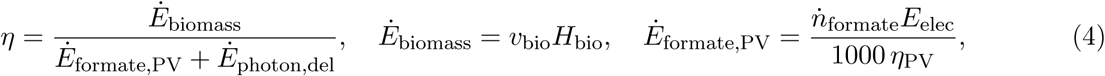

This equation defines the diagnostic *η* in Table S10. It includes the solar-equivalent electrical demand for formate and the photon input delivered to the culture, but does not charge unused collector sunlight. The headline whole-collector efficiency is defined separately below. Stage efficiencies (*η*_PV_, *η*_dichroic_, *η*_electrolysis_) and the formate-saving metric *η*_save_ describe individual contributions. Here *H*_bio_ = 22.4 kJ g*^−^*^1^ and *E*_elec_ is the total electrical demand per mole of formate delivered to the culture. It is carried as two named terms rather than as a single electrolysis constant: the *electrosynthesis energy E*_synth_ and the *CO*_2_ *capture energy E*_cap_, with *E*_elec_ = *E*_synth_ + *E*_cap_ = 500.0 + 419.9 and 270.0 + 19.16 kJ mol*^−^*^1^ in the two scenarios (see below). The two electrical contributions are shown separately in Fig. 7a. Headline process and fixed-geometry diagnostic efficiencies include both; the cell-scale *η*_cell_ omits capture (Table S10, Section S6.12). Finally *Ė*_photon, del_ = PPFD *A*_gDW_(*hcN*_A_*/λ*). A formate-saving efficiency *η*_save_ = Δ*ṅ*_formate_*E*_elec_*/*(1000 *η*_PV_ *Ė*_photon,del_) isolates the photosynthetic contribution, with Δ*ṅ*_formate_ referenced to the dark fixed-growth demand for the same pathway and wavelength.

#### System boundary

Headline process efficiencies use the *closed energy-accounting boundary*: all solar power incident on the collector aperture enters the denominator, including any unused energy, with no export credit. The reactor exchanges matter and heat with its surroundings; “closed” refers only to this accounting convention. On a unit-reactor-area basis,

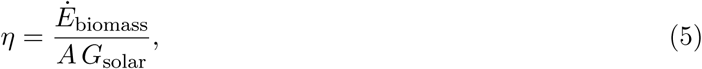

where *G*_solar_ = 1000 W m*^−^*^2^ (AM1.5G), *A* is the collector-to-reactor area ratio, and biomass chemical power is expressed per unit reactor area. The cell-scale and fixed-geometry diagnostics in Table S10 instead include specified photon and photovoltaic inputs, as defined above. Supplementary route-level calculations use that partial-input accounting and are labelled separately. Crop values are shown only beside the whole-collector efficiency and only as context: the calculation is a steady design point, not a prediction of seasonal land productivity.

The aperture is sized so that photovoltaic output covers the whole electrical load: all energy entering the system originates as sunlight on that aperture, and no unpriced electrical or thermal utility is admitted. Inside the boundary are the dichroic split and its optical losses; the direct photon flux delivered to the culture; photovoltaic conversion of the complementary spectrum; electrosynthesis of formate; any light-emitting-diode top-up, charged at its wall-plug efficiency; and CO_2_ supply. That last item is the *CO*_2_ *capture energy E*_cap_, drawn as electrical work from the same photovoltaic branch as electrosynthesis and reported as its own term throughout rather than folded into the electrolysis constant *E*_synth_, so that what the boundary spends on carbon can be read off directly (Fig. 7a, Table S10). Capture is charged so that carbon supply is not treated as a free input. This also keeps the boundary explicit when seasonal crop values, which include atmospheric carbon acquisition internally, are shown for context. The *capture basis*, meaning the separation duty priced as *E*_cap_, is paired to scenario maturity: deployed all-electric sorbent direct air capture at *E*_cap_ = 419.9 kJ mol*^−^*^1^ for the state of the art (2000 kWh thermal plus 650 kWh electric per tonne CO_2_, the regeneration heat supplied resistively since the boundary admits no free thermal source), and the minimum second-law separation work *W*_min_ = *RT* ln(1*/y*_CO2_), *E*_cap_ = 19.16 kJ mol*^−^*^1^ from 440 ppm air, at the thermodynamic limit. Both constants, and the derivation of the deployed figure, are in `PaperScripts/reactor_geometry/data/optimize_flexible_aperture.py`; the maturity-matched pairing is applied to the headline closed-boundary optimisation in the same directory’s `optimize_hybrid_dichroic_led.py`. The maturity-matched direct-air-capture rows remain the primary basis; alternative carbon supplies are reported as sensitivities. At *η*_II_ = 0.20, a 14 wt% flue-gas stream requires 29.3 kJ mol*^−^*^1^ and air requires 95.8 kJ mol*^−^*^1^, a like-for-like reduction of 69%. The larger numerical change from deployed all-electric DAC (419.9 kJ mol*^−^*^1^) to the point-source estimate is a cross-scenario sensitivity rather than a comparison of deployed capture plants. These bases are tabulated in Table S19. The point-source basis follows the previous stochastic techno-economic assessment of this system^2^. Flue-gas supply answers the different question of co-location with an industrial point source. It is treated as a separate sensitivity because its carbon supply differs from direct air capture.

Outside the boundary are the biological and process terms the model does not resolve: retinal supply, construction failure and the burden of heterologous expression; nutrients other than the carbon and oxygen balances solved explicitly; downstream recovery, separation and drying; plant capital, financing, collector tracking and inter-row spacing; weather, downtime and duty cycle, since *η* is evaluated at a steady design irradiance whereas crop figures are integrated over a growing season. Mixing and circulation are likewise outside the headline metric. Two illustrative duties are charged in separate sensitivities; the *∼*0.35 W L*^−^*^1^ case in Table S11 lowers *η* by *≈*0.03 percentage points at the state of the art. Converting *η* into an areal productivity requires the omitted terms and is not attempted. The headline value is therefore a core conversion efficiency, not a full life-cycle or techno-economic result.

#### Bounded design-space optimisation

The constrained-optimal designs of Fig. 7 maximise *η* over six continuous variables: absorption maximum *λ ∈* [450, 600] nm, culture dry weight CDW *∈* [0.05, 10] g L*^−^*^1^, surface PPFD *∈* [50, 50,000] *µ*mol m*^−^*^2^ s*^−^*^1^, depth *V/A ∈* [0.005, 0.1] m, membrane fraction allocated to rhodopsin *f*_mem_ *∈* [0, 1.0], and dichroic transmission *t ∈* [0, 1], for every discrete geometry, pathway and electrolyser combination. Three reactor limits bound the search at the operating point: volumetric O_2_ uptake below an oxygen transfer rate of 100 mmol L*^−^*^1^ h*^−^*^1^, volumetric formate uptake below 100 mmol L*^−^*^1^ h*^−^*^1^, and surface PPFD below 3*×* the unconcentrated AM1.5G rhodopsin-band irradiance after dichroic efficiency. Full membrane allocation corresponds to *∼*94.8% of the original uncapped pump inventory, whose footprint would be 21.1 m^2^ gDW*^−^*^1^. The realised per-gDW proton flux, not only the per-molecule rate, queries the fixed-growth lookup, so allocation changes both self-shading and ATP supply. The volumetric limits break the CDW–*V/A* degeneracy, because gas-transfer and feed loads depend on density directly (Section S6). CDW, PPFD and *V/A* were searched on a log_10_ scale and wavelength, *f*_mem_ and *t* linearly, each evaluation calling the same physics–metabolism–optics pipeline used throughout. Global search used differential evolution (population 32, up to 250 iterations, bounded mutation [0.4, 1.0], recombination 0.7, deferred updating, internal polishing disabled) followed by an L-BFGS-B polish accepted only where it improved the objective. Feasibility is enforced by hard rejection rather than a soft penalty: a design breaching the concentration limit, or whose worst-case volumetric utilisation exceeds unity, is assigned a large positive objective that rises with overshoot, which keeps the search out of the infeasible region without deforming the objective surface inside it. Reported design variables are rounded conservatively, CDW floored, so that a rounded design cannot breach the volumetric limits. All six variables are searched together, so the design-lever stack (Fig. 7b) is a post-hoc attribution: continuous variables are held at their optimised values and the discrete choices switched on in sequence, with *η* re-evaluated at each step. Because levers interact, bar heights depend on ordering whereas endpoints do not (Section S6). One transport limit is active in each scenario: the 3*×* concentration cap at the state of the art and the formate feed at the thermodynamic limit. Yet none of the three bounds *η*: re-optimising with each relaxed tenfold moves it by at most 0.05 points, and relaxing either transport limit moves it by under 0.002 (Section S6). The membrane allocation reaches its imposed upper bound. Headline efficiencies cap rhodopsin occupancy at 45%, an overproduction target inferred from heterologous expression rather than demonstrated in this host; the unrestricted optimum is reported separately. Efficiency saturates with optical depth and barely changes along the density–depth trade at fixed areal loading, so similar modelled efficiencies occur across a range of density–depth combinations whose practical feasibility remains untested. Heat removal is outside the accounting boundary; a balance on metabolic and absorbed solar heat confirms it is not a hidden fourth binding constraint (Section S6). Per-scenario parameters are tabulated in Table S17.

### Quantification and Statistical Analysis

#### Monte Carlo uncertainty propagation

Metabolic outputs are deterministic LP optima. Process-efficiency intervals are the 2.5th–97.5th percentiles from re-optimising designs under Latin-hypercube parameter draws; no classical hypothesis tests were performed. The headline whole-collector calculation used 512 feasible draws per scenario, implemented in `PaperScripts/reactor_geometry/data/propagate_ closed_boundary.py`. Each draw varied PV efficiency, formate-electrosynthesis energy, biomass heating value, effective pump scale, aggregate fixed-growth formate demand, solar-band delivery, the concentration limit, and O_2_-transfer and formate-feed limits. CO_2_ capture was charged in every draw: deployed all-electric sorbent direct air capture was sampled from a triangular distribution over 300–550 kJ mol*^−^*^1^ with mode 419.9 in the state-of-the-art scenario; the thermodynamic-limit scenario used minimum separation work of 19.2 kJ mol*^−^*^1^.

For each draw, density, surface PPFD, depth and dichroic transmission were searched by differential evolution on the optimum manifold identified by the full six-variable point-estimate calculation: external cylinder, membrane allocation at the 45% cap, and wavelength 550 nm (state of the art) or 600 nm (thermodynamic limit). The resulting medians and intervals were 3.25% (2.74–3.89%) for the state of the art and 10.53% (9.09–12.24%) at the thermodynamic limit. These intervals describe the sampled model assumptions. They do not include structural model uncertainty, seasonal operation, construction failure or economic uncertainty (Tables S23, S20 and S26).

The joint carotenoid–retinal calculation used the same 512 process draws per scenario and additionally sampled Kin4B8 transfer uniformly from 0.50 to 0.70, optical delivery from 0.75 to 1.0, and the borrowed antenna bandwidths from 0.8 to 1.2 times their reference values. Each draw was re-optimised at 45% occupancy, giving 3.35% (2.79–4.04%) for the state of the art and 10.35% (8.93–12.02%) at the thermodynamic limit. Pairing every antenna draw with its retinal-only result isolates the incremental antenna effect: median +0.107 percentage points (*−*0.034 to +0.284; 93.2% positive) for the state of the art, and *−*0.178 points (*−*0.216 to *−*0.140; none positive) at the thermodynamic limit. No probability distribution was assigned to antenna assembly burden because none is measured; the reported 6.7% break-even burden is a sensitivity bound.

#### Optimisation audit and crop context

The manifold restriction was audited on the headline whole-collector objective, not the route-boundary diagnostic. Twelve draws spanning the calculated efficiency distribution in each scenario were re-optimised by differential evolution over all six continuous variables and every reactor geometry. The largest absolute difference from the propagated-search result was 4.8 *×* 10*^−^*^5^ percentage points; the median differences were *−*1.2 *×* 10*^−^*^5^ and *−*0.3 *×* 10*^−^*^5^ points for the state-of-the-art and thermodynamic-limit scenarios, respectively (Table S24). Thus the earlier random-restart convergence problem is removed at the precision reported here. This numerical audit does not test the model equations or omitted deployment terms.

Crop values use incident solar radiation integrated over a growing season^21^; the model uses the whole incident collector input at a steady design irradiance. Table S18 reports these accounting bases with the values shown in Fig. 7f.

## Supplemental Information

Supplemental Information includes 15 figures and 29 tables and can be found with this article online.

## Notes

### Competing Interest Statement

The authors have declared no competing interest.

