## Supplementary Information for "A conditional upper-bound design map for rhodopsin-augmented electro-microbial CO_2_-to-biomass conversion"

### Supplemental Methods

Symbols and model terms used throughout are defined in Table S13.

#### S1 Quantum-yield model: one-dimensional surface hopping

The photoisomerisation quantum yield $\Phi_{\rm QY}$ enters the Poisson activation model via $P_{\rm act} = 1 - \exp(-k_{\rm abs}\Phi_{\rm QY}\tau)$ with no additional prefactor. The main Methods summarises the model and its MCMC calibration; this section gives the full equation-level derivation. Implementation is in `Bin/PhysBin/QuantumYield/`; calibration of the four free parameters is in Section S2.

##### S1.1 Reduced-coordinate torsional potential-energy surface

The primary photochemical event is treated as motion along one collective coordinate, the $C_{13}{=}C_{14}$ torsion $\phi$. Bond-length-alternation and hydrogen-out-of-plane modes are not propagated; their effect is folded into the effective torsional inertia and diabatic coupling. Two adiabatic surfaces are built on a uniform grid in $\phi$. The ground state is the lower envelope of two diabatic parabolas joined at $\phi_{\rm CI}$:

$$E_{\rm S_0}(\phi) = \min\big[\, q_{\rm trans}(\phi),\ q_{\rm cis}(\phi) \,\big], \tag{S1}$$

where each branch $q_w(\phi) = a_w\phi^2 + b_w\phi + c_w$ is the unique quadratic satisfying $q_w(\phi_w) = E_w$, $q'_w(\phi_w) = 0$ and $q_w(\phi_{\rm CI}) = E_{\rm CI}$, giving a double well with a cusp. The convention is $\phi_{\rm trans} = -90°$, $\phi_{\rm cis} = +110°$, $\phi_{\rm CI} = 0°$, with *cis* slightly less stable than *trans*. The excited state is a cubic,

$$E_{\rm S_1}(\phi) = c_3\phi^3 + c_2\phi^2 + c_1\phi + c_0, \tag{S2}$$

whose coefficients are fixed by four anchors: the vertical Franck–Condon energies at the two minima,

$$\begin{aligned} E_{\rm S_1}(\phi_{\rm trans}) &= E_{\rm trans} + \Delta_{\rm abs}^{\rm trans}, \\ E_{\rm S_1}(\phi_{\rm cis}) &= E_{\rm cis} + \Delta_{\rm abs}^{\rm cis}, \end{aligned} \tag{S3}$$

the excited-state energy at the crossing,

$$E_{S_1}(\phi_{CI}) = E_{CI} + \Delta E_{CI}, \tag{S4}$$

and a stationarity condition locating the $S_1$ minimum on the product side,

$$\phi_{min} = \phi_{CI} + \tfrac{1}{4}\big(\phi_{cis} - \phi_{CI}\big). \tag{S5}$$

This placement guarantees a wavepacket launched from the *trans* Franck–Condon point descends *through* the crossing on a downslope in one ballistic pass. The momentum delivered to the seam is then set by the Franck–Condon-to-crossing energy difference, making the predicted yield sensitive to $\lambda_{max}$ without imposing a trend by hand.

##### S1.2 Franck–Condon excitation and the crossing region

A spectral shift displaces the absorption anchors by the photon-energy difference,

$$\Delta E(\lambda) = \frac{hc}{\lambda} - \frac{hc}{\lambda_{ref}}, \qquad \lambda_{ref} = 568 \text{ nm}, \tag{S6}$$

so both terms in Eq. S3 carry the full shift while the crossing carries only a fraction $\beta$,

$$E_{CI} \to E_{CI} + \beta\,\Delta E(\lambda), \qquad \beta \in [0, 1]. \tag{S7}$$

$\beta$ controls how much of a spectral perturbation is absorbed by the crossing topology rather than the Franck–Condon gradient: at $\beta = 1$ the yield is nearly flat in wavelength; at $\beta = 0$ a blue shift raises the Franck–Condon point, increases kinetic energy at the seam and raises the yield. Because $\Delta E(\lambda) = 0$ at the calibration wavelength (Eq. S6 at $\lambda = 568$ nm), $\beta$ receives *zero* likelihood constraint and is treated as an explicit source of predictive uncertainty (Section S2).

Near the crossing the surfaces approach within $\Delta E_{CI}$ (Eq. S4); a positive gap opens an avoided crossing that suppresses hopping. A touching in one coordinate is a crossing point, not a conical intersection (a genuine conical intersection requires a branching space of at least two coordinates, not represented here). The coupling $V$ is estimated as half the local adiabatic gap scaled by $c_s$, and $\alpha = |F_2 - F_1|$ is read from the surfaces at the seam.

##### S1.3 Landau–Zener non-adiabatic transition probability

The probability that a trajectory traversing the crossing remains diabatic is:

$$P_{hop} = \exp\!\left(-\frac{\pi V^2}{2\hbar\,|v|\,\alpha}\right). \tag{S8}$$

Symbols are in Table S1. The prefactor differs from the textbook convention $\exp(-2\pi V^2/\hbar|v|\alpha)$ by a constant absorbed into the posterior for $c_s$. This is not the Zhu–Nakamura expression[1,2], which adds tunnelling and turning-point corrections not carried here. The $v \to \infty$ limit gives $P_{hop} \to 1$ (sudden, diabatic crossing); $v \to 0$ gives adiabatic abortive return.

The scheme is Landau–Zener surface hopping as described in the main Methods: one classical trajectory on one adiabatic surface at a time, no electronic amplitudes or coherences propagated. On a successful hop the velocity is rescaled to conserve energy; frustrated hops (insufficient kinetic energy) are rejected. Each crossing-region passage allows at most one hop.

$S_1$ motion is propagated as a *coherent ballistic* trajectory (no friction), because the wavepacket reaches the crossing within ~200 fs[3,4]. Only after a successful hop are Langevin friction $\gamma$ and thermal force ($T = 300$ K) switched on. Integration uses a semi-implicit Euler scheme at $\Delta t = 0.1$ fs,

$$v_{n+1} = (1 - \gamma\,\Delta t)\,v_n + \frac{F(\phi_n)}{m}\,\Delta t,$$
$$\phi_{n+1} = \phi_n + v_{n+1}\,\Delta t, \tag{S9}$$

with $F = -\mathrm{d}E/\mathrm{d}\phi$ on the active surface and $\gamma \equiv 0$ on $S_1$. Reflective boundaries at $\phi = \pm 200°$ keep trajectories in the chemically relevant range without absorbing population.

##### S1.4 Trajectory ensemble and the QY estimator

For each parameter set, 1024 trajectories are launched from the *trans* Franck–Condon point with a thermal velocity distribution. To represent chromophore-environment heterogeneity, $m$ and $\gamma$ per trajectory are multiplied by a truncated-normal factor (mean 1.0, s.d. 0.1, clipped to $[0.1, 5.0]$). The yield is:

$$\Phi_{\mathrm{QY}} = \frac{N_{\mathrm{cis}}}{N_{\mathrm{traj}}}, \qquad N_{\mathrm{traj}} = 1024, \tag{S10}$$

where trajectories that fail to hop, or relax back to *trans*, count as abortive. Wavelength predictions draw parameter sets from the posterior, sample $\beta \sim \mathrm{Uniform}(0,1)$ per draw, regenerate shifted surfaces (Eqs. S3–S7) and re-run the ensemble; the spread is reported as a posterior-predictive band (Section S2).

##### S1.5 Parameters, units, and prior ranges

The friction prior ($2 \times 10^{-3}$–$0.02$ fs$^{-1}$) spans intermediate damping where protein-bound retinal settles on the 50–500 fs scale; outside this window the yield collapses to either a random-partition regime ($\Phi_{\mathrm{QY}} \approx 0.5$) or complete *cis* trapping. The crossing threshold is 0.50 eV, exceeding the upper prior bound on $\Delta E_{\mathrm{CI}}$ so the inferred gap never artificially eliminates hopping. A soft half-normal prior ($\sigma = 0.05$) penalises $\Phi_{\mathrm{QY}} > 0.90$, encoding that measured yields are typically 0.3–0.7[5,6].

##### S1.6 The effective absorption cross-section, and what its calibration absorbs

The pump conversion uses $\sigma_{\mathrm{eff}} = 4.5\,\sigma_0$, where $\sigma_0 \approx 2.41 \times 10^{-20}$ m$^2$ follows from the measured molar extinction coefficient. The 4.5× value is inherited from the model calibration, but its independent pump-flux datum is not documented. A factor above unity cannot be explained by orientation averaging, oligomeric crowding or trap-state losses, all of which *reduce* an effective cross-section. It also cannot be assigned to the per-molecule cross-section, the ≈1 $H^+$ photocycle stoichiometry, or additional rhodopsin abundance: a 4.5× abundance increase would require 94.8 m$^2$ gDW$^{-1}$ against a 20-m$^2$ budget (Section S4).

Table S1: Free and structural parameters of the one-dimensional surface-hopping quantum-yield model. The four inferred parameters are sampled from uniform priors over the ranges shown (Section S2); the CI-shift fraction $\beta$ is not inferred but sampled from its prior during wavelength prediction because it has zero likelihood weight at the wild-type calibration wavelength.

| **Symbol** | **Name / description** | **Prior range** | **Units** | **Physical role** |
|---|---|---|---|---|
| $m$ | Effective torsional inertia | [0.01, 5.0] | $\mathrm{eV{\cdot}fs^2/deg^2}$ | Sets nuclear velocity at the crossing seam |
| $\gamma$ | Ground-state Langevin friction | $[2 \times 10^{-3}, 0.02]$ | $\mathrm{fs^{-1}}$ | Dissipation governing $S_0$ relaxation after the hop |
| $c_s$ | Diabatic coupling scale | [0.1, 10.0] | 1 | Multiplies the adiabatic-gap estimate of $V$ |
| $\Delta E_{\mathrm{CI}}$ | $S_1$–$S_0$ gap at the crossing | [0.0, 0.30] | eV | Opens an avoided crossing; suppresses hopping |
| $\beta$ | CI-shift fraction | [0, 1] | 1 | Fraction of spectral shift absorbed by the crossing (not inferred) |
| $V$ | Diabatic coupling at crossing | , | eV | Numerator of the LZ (Landau–Zener) exponent (derived from $c_s$) |
| $v$ | Nuclear velocity at seam | , | $\mathrm{deg\,fs^{-1}}$ | Per-trajectory; controls $P_{\mathrm{hop}}$ |
| $\alpha$ | Diabatic slope difference | , | $\mathrm{eV\,deg^{-1}}$ | Width of the nonadiabatic region |

The scale is retained as a lumped model parameter that may absorb error in the activation model, the molecule-to-biomass conversion or the flux index used by the metabolic lookup; available evidence does not identify which. Its effect on absolute pump rates and formate savings is large: at 45% occupancy, changing the forward pump-rate scale from 0.5× to 2× moves the capture-charged whole-collector SOTA optimum from 2.69% to 4.63% (Section S6.8). The optimiser cannot fully compensate by increasing PPFD because the nominal optimum already reaches the solar-concentration limit. A common multiplier cancels approximately from pump-rate ratios only when both rates are in the low-activation regime. It does not generally cancel from the nonlinear Poisson response or from downstream metabolic optimisation; pathway and spectral robustness must therefore be assessed by the reported sensitivity analyses.

#### S1.7 Why a one-dimensional model, and its limitations

A full multireference QM/MM treatment needs days per variant[4,7–10], prohibitive for a wavelength sweep coupled to a metabolic optimiser. The reduced model retains the three features controlling wavelength dependence: Franck–Condon-to-crossing gradient, velocity-dependent hop probability and post-hop branching. It runs at $\sim 10^3$ trajectories per second on one GPU, enabling full posterior propagation.

The model omits bond-length-alternation and hydrogen-out-of-plane coordinates, so residue-specific $S_1$ electronic effects are missing; it reproduces the magnitude and the declining wild-type-to-red trend but not the modest upward mutant trend in some point estimates (Section S9). Nuclei are classical, without zero-point energy or tunnelling; diabatic coupling is estimated from the adiabatic gap rather than computed *ab initio*; and the protein enters only through parametrised friction, mass and CI-shift fraction.

Landau–Zener is also derived for a two-state avoided crossing with locally linear diabats and a well-defined crossing velocity, whereas the $S_1/S_0$ funnel of retinal is a genuine conical intersection whose branching topology one coordinate cannot carry. The model is a calibrated functional form for the wavelength trend, not a first-principles yield prediction. The projected-out, residue-specific coordinates probably explain the mutant-ordering disagreement (Section S9).

These limitations affect the predicted pump input to the metabolic and reactor calculations. They are addressed at the process level by propagating the QY posterior and by the forward pump-scale sensitivity (Section S6.8); neither calculation turns the reduced surface-hopping model into a first-principles electronic-structure treatment.

### S2 MCMC calibration and posterior uncertainty

The photophysical models comprise two independently calibrated blocks: the four-parameter surface-hopping quantum-yield model (Section S1) and the twelve-parameter photocycle model (Section S3), both sampled with affine-invariant ensemble MCMC. Because the quantum-yield block is constrained by a single observable, the emphasis is on identifiability structure and on propagating limited information into wavelength-dependent predictive uncertainty rather than on a tightly identified fit.

#### S2.1 Likelihood and the single calibration observable

Calibration uses exactly one observable, the wild-type photoisomerisation quantum yield $\Phi_{\text{iso}} = 0.64$ with uncertainty $\sigma_{\text{QY}} = 0.04$[5,11]. For $\boldsymbol{\theta} = (m, \gamma, c_s, \Delta E_{\text{CI}})$ the model returns a simulated yield from the 1024-trajectory ensemble (Eq. S10), and

$$\ln \mathcal{L}(\boldsymbol{\theta}) = -\frac{1}{2}\left(\frac{\Phi_{\text{QY}}(\boldsymbol{\theta}) - 0.64}{\sigma_{\text{QY}}}\right)^2 - \frac{1}{2}\left[\frac{\max\big(0,\, \Phi_{\text{QY}}(\boldsymbol{\theta}) - 0.90\big)}{0.05}\right]^2, \tag{S11}$$

the second term being a one-sided half-normal prior penalising yields above 0.90, since measured rhodopsin yields lie in 0.3–0.7[5,6]. Photocycle timing is excluded: at 300 K thermal fluctuations exceed the settling threshold for all admissible parameter combinations, so that observable carries no discriminating information. The calibration is therefore a one-observable, four-parameter inverse problem and is structurally under-determined, as reflected in the posterior.

#### S2.2 Priors and the unit-hypercube transform

Each parameter has an independent uniform prior over its physical range (Table S1): $m \in [0.01, 5.0]$ eV·fs$^2$/deg$^2$, $\gamma \in [2 \times 10^{-3}, 0.02]$ fs$^{-1}$, $c_s \in [0.1, 10.0]$ and $\Delta E_{\text{CI}} \in [0.0, 0.30]$ eV. Sampling uses a unit-hypercube coordinate mapped affinely to those bounds, so the prior is flat and proposals are scale-matched; proposals leaving the hypercube receive $\ln \mathcal{L} = -\infty$. The friction range brackets the physical 50–500 fs settling regime and avoids assigning substantial prior mass to qualitatively different damping regimes. A separate 1024-point Sobol likelihood integration widened only this prior to $2 \times 10^{-4}$–0.1 $\mathrm{fs}^{-1}$ and re-ran 512 trajectories per point. The median wavelength trend was retained: the paper-range and expanded-prior diagnostics gave QY medians of 0.675 versus 0.682 at 520 nm and 0.622 versus 0.596 at 600 nm. The expanded prior widened the 95% ranges from 0.517–0.761 to 0.562–0.897 at 520 nm and from 0.519–0.718 to 0.450–0.717 at 600 nm. Thus the friction bound affects predictive width more than the median red-shift trend. This likelihood-weighted audit (effective sample sizes 51 and 35) is a prior-sensitivity diagnostic, not a replacement for the production MCMC chain (`PaperScripts/Supplementary/data/audit_qy_friction_prior.py`; `Results/Supplementary/qy_friction_prior_audit_summary.csv`).

##### S2.3 Sampler, move mixture, and convergence

Sampling uses emcee[12,13] with 32 walkers for 1100 steps (300 burn-in, 800 retained; 25,600 draws at 1024 trajectories per evaluation). A mixture of 80% `DEMove` and 20% `DESnookerMove` replaces the default stretch move, because $c_s$ and $\Delta E_{\mathrm{CI}}$ are strongly anti-correlated (larger coupling and smaller gap give the same hop probability, Eq. S8), forming a diagonal ridge along which the stretch move mixes poorly.

Convergence was assessed three ways. Integrated autocorrelation time ranged from 61 to 69 steps ($\sim$12 autocorrelation times per chain, effective sample sizes 371–423). Split-$\hat{R}$ was 1.02, 1.03, 1.00 and 1.00 for $m$, $\gamma$, $c_s$ and $\Delta E_{\mathrm{CI}}$ (all below 1.05), and mean acceptance fraction was 0.22. Posterior medians ($\pm 1\sigma$, half-width of the 16th–84th-percentile interval) are

$$m = 1.42 \pm 1.60, \quad \gamma = 0.010 \pm 0.007\ \mathrm{fs}^{-1}, \quad c_s = 2.65 \pm 2.78, \quad \Delta E_{\mathrm{CI}} = 0.08 \pm 0.08\ \mathrm{eV}. \tag{S12}$$

##### S2.4 Identifiability and the role of the $\beta$ prior

The half-widths in Eq. S12 are comparable to or larger than the medians: one scalar observable cannot localise four parameters, so the posterior is broad and prior-dominated along sloppy directions, as expected for an under-determined inverse problem.

Wavelength dependence is governed by the CI-shift fraction $\beta$, which determines how a spectral shift partitions between Franck–Condon gradient and crossing energy (Eqs. S6–S7). Because the calibration point carries zero spectral shift, $\beta$ has *zero* likelihood contribution and is entirely unconstrained: during prediction it is sampled from $\mathrm{Uniform}(0, 1)$ per draw, and the predicted band encodes maximal prior uncertainty about the spectral-tuning mechanism. The predicted decline with red shift and its broad credible intervals are therefore insensitive to residual ridge uncertainty in the four inferred parameters.

##### S2.5 Posterior-predictive quantum yield

Predictions marginalise over the joint posterior and the $\beta$ prior,

$$p(\Phi_{\mathrm{QY}} \mid \lambda) = \int p(\Phi_{\mathrm{QY}} \mid \lambda, \boldsymbol{\theta}, \beta)\ p(\boldsymbol{\theta} \mid \mathrm{data})\, p(\beta)\, \mathrm{d}\boldsymbol{\theta}\, \mathrm{d}\beta, \tag{S13}$$

estimated by re-running the ensemble for each draw on surfaces regenerated at the target wavelength. At the calibration wavelength the posterior-predictive median is $0.60 \pm 0.05$, undershooting the target by ∼6% through Bayesian shrinkage but within $1\sigma$ and consistent with $0.58 \pm 0.05$[14]. The 95% credible interval (2.5th–97.5th percentiles of 200 draws per wavelength; `Bin/PhysBin/QuantumYield/wavelength_raw_qy_samples.csv`) spans 0.51–0.74 at 520 nm and 0.44–0.69 at 600 nm. It widens in both directions from the anchor as the unconstrained $\beta$ prior compounds with the broader posterior, so the band is narrowest at the anchor rather than over a sub-range. These bands propagate into all downstream physics-layer quantities; metabolic-layer outputs are deterministic LP optima at each sampled operating point.

### S3 Photocycle master equation (full model)

The photocycle lifetime $\tau$ enters the Poisson pump model (Section S1) and sets the time available for a productive isomerisation within one turnover. Implementation is in `Bin/PhysBin/Photocycle_time/`; the twelve free parameters are calibrated with the sampler of Section S2. At the wild-type wavelength the model returns $\tau_M = 5.2$ ms against a 5 ms target and $\tau_{\text{total}} = 13.0$ ms against 15 ms (`PaperScripts/physics_data/physics_sweep.csv`), the total 14% low but well inside a posterior width of ±18.8 ms.

#### S3.1 Seven-state master equation and steady state

Dynamics follow a reversible seven-state cycle. Light drives a net direction through bR, K, L, $M_1$, $M_2$, N and O[6], with populations evolving under

$$\frac{d\mathbf{p}}{dt} = \mathbf{K}\,\mathbf{p}, \qquad \mathbf{K}\,\mathbf{p}^* = \mathbf{0}, \qquad \sum_i p_i = 1. \tag{S14}$$

Rate constants fill off-diagonal entries of $\mathbf{K}$; diagonal elements follow by column conservation. Under continuous illumination the system reaches a non-equilibrium steady state (solved by least squares on the singular system). The closed directed cycle forces a single common circulating current through every edge,

$$J = k_i^+\, p_i^* - k_i^-\, p_{i+1}^* \qquad \text{(equal at all edges } i\text{)}, \tag{S15}$$

which is the steady-state turnover flux of the pump.

#### S3.2 Marcus barriers and Langevin dipole-saturation coupling

Forward rates follow Marcus-type activated kinetics[15]. For the rate-limiting $M_1 \to M_2$ transition,

$$W_M = \frac{(\Delta G_0 + \lambda_0)^2}{4\,\lambda_0} + \Delta E_{\text{eff}}(\Delta\lambda), \tag{S16}$$

with $\lambda_0$ the reorganisation energy and $\Delta G_0$ the driving force. The first term is the standard Marcus activation free energy[15]; the early (K, L) and late (N, O) transitions use fixed base barriers with the same Arrhenius form. The spectral perturbation couples photon-energy change to an electrostatic barrier modulation through a Langevin dipole-saturation function,

$$L(x) = \coth(x) - \frac{1}{x}, \qquad x = \frac{\Delta E_{\text{opt}}}{E_{\text{pole}}}, \tag{S17}$$

with $\Delta E_{\text{opt}}$ the optical shift and $E_{\text{pole}}$ a characteristic electrostatic scale. $L(x)$ is linear for small shifts and saturates at $\pm 1$ for large shifts, bounding the response without extra fitting parameters; a 1% linear leakage keeps it differentiable. The coupling enters as $\Delta E_{\text{eff}} = \alpha_L \, E_{\text{pole}} \, L(x)$, with conformational disorder of amplitude $\sigma_{\text{conf}}$ broadening the barrier distribution inhomogeneously.

Late-cycle transitions see attenuated coupling, modelled as dipole-field shielding ($U \propto r^{-2}$, hence $r^{-3}$ decay),

$$\chi_i = \left(\frac{R_{\text{core}}}{R_i}\right)^3, \qquad R_{\text{core}} = 2.0 \text{ Å}, \;\; R_N = 11.0 \text{ Å}, \;\; R_O = 20.0 \text{ Å}, \tag{S18}$$

so the N and O transitions see the core perturbation reduced by $\chi_N$ and $\chi_O$, reflecting decreasing electrostatic sensitivity with distance from the Schiff base.

##### S3.3 Arrhenius rates and detailed-balance reverse rates

Each forward rate combines the barrier with a quantum transition-state attempt frequency and a Grote–Hynes friction correction,

$$k_i^+ = \frac{k_B T}{h} \, \kappa_{\text{fric}} \, \exp\left(-\frac{W_i}{k_B T}\right), \tag{S19}$$

with $\kappa_{\text{fric}} \leq 1$ a Grote–Hynes friction correction for barrier recrossing (Table S2), set to unity for $M_2 \rightarrow N$, treated as a conformational reprotonation switch rather than a diffusive crossing; this is a modelling assumption, not an independently measured property of that transition. Reverse rates follow detailed balance,

$$k_i^- = k_i^+ \, \exp\left(\frac{G_{i+1} - G_i}{k_B T}\right), \tag{S20}$$

with state free energies from literature photocycle energetics[6] and the sampled driving forces for $M_1 \rightarrow M_2$, $M_2 \rightarrow N$ and $N \rightarrow O$. This guarantees zero net current without light-driven forcing.

##### S3.4 Lifetimes from occupancy–flux relations and calibration

For a cyclic Markov chain in steady state the residence times and cycle period follow from the occupancies and the common current of Eq. S15[6,16]:

$$\tau_i = \frac{p_i^*}{J}, \qquad \tau_M = \frac{p_{M1}^* + p_{M2}^*}{J}, \qquad \tau_{\text{total}} = \sum_i \tau_i = \frac{1}{J}. \tag{S21}$$

The twelve free parameters (Table S2) are calibrated against wild-type targets $\tau_M^{\text{obs}} \approx 5$ ms and $\tau_{\text{total}}^{\text{obs}} \approx 15$ ms at 568 nm, pH 7, 20°C[6,16], using the sampler of Section S2. The posterior broadens substantially below 540 nm and to the red of 568 nm, where the lengthening photocycle amplifies parameter uncertainty, so the model is used for kinetic trends rather than precise mutant lifetimes away from the anchor.

##### S3.5 Poisson saturation regime and the role of $\tau$

The per-molecule pump rate depends on lifetime through the saturation parameter

$$x = \sigma_{\text{eff}} \cdot \text{QY} \cdot \tau \cdot I, \qquad R_{\text{pump}} = \frac{1 - e^{-x}}{\tau}, \tag{S22}$$

At the main operating conditions (520–568 nm, 100 PPFD) $x \approx 0.031$–$0.051$ (`PaperScripts/physics_data/physics_sweep.csv`), so $R_{\mathrm{pump}} \approx \sigma_{\mathrm{eff}} \cdot \mathrm{QY} \cdot I$ to within 2.5% and the logarithmic sensitivity $|\mathrm{d}\ln R_{\mathrm{pump}}/\mathrm{d}\ln\tau| < 2.6\%$; lifetime cancels to leading order and quantum yield carries the dominant wavelength dependence (see Section S7). At 600 nm the longer $\tau$ pushes $x \approx 0.61$ into partial saturation, where $(1-e^{-x})/x \approx 0.75$, reducing the linear-regime estimate by ∼25%; the broadened lifetime posterior away from the 568 nm anchor has its largest effect in this partially saturated regime.

### S4 Enzyme-constrained metabolic model (EC-FBA/RBA)

This is the expanded form of the metabolic layer summarised in the main Methods. Symbols are collected in Table S3; numerical parameters and curated turnover numbers are in Tables S14 and S16.

#### S4.1 Base linear programme and maintenance demands

The model is an LP on the *C. necator* H16 reconstruction RehMBEL1391[17,18] (1538 reactions, 1348 metabolites, 1350 genes; SBML L3V1). Following standard practice[19–21],

$$\max\ v_{\mathrm{bio}} \quad \text{s.t.} \quad \mathbf{S}\,\mathbf{v} = \mathbf{0},\ \ \mathbf{l} \le \mathbf{v} \le \mathbf{u}, \tag{S23}$$

with $\mathbf{S} \in \mathbb{R}^{m\times n}$ the stoichiometric matrix and $\mathbf{v}$ in mmol gDW$^{-1}$ h$^{-1}$ (biomass in h$^{-1}$). Two maintenance demands are embedded: GAM = 135.0 mmol ATP per gDW formed (drain $\mathrm{GAM}\cdot v_{\mathrm{bio}}$) and NGAM = 3.0 mmol ATP gDW$^{-1}$ h$^{-1}$ independent of $v_{\mathrm{bio}}$[22]. Both are literature values from Jahn et al.[23], adopted unchanged by Dronsella et al.[24]. Neither was refitted to the yield measurements in this analysis; the benchmark in Section S12 is conditional on these maintenance values and the proteome calibration.

##### Pirt decomposition and the basis of the specific fluxes

Because GAM enters per gram of biomass *formed* while NGAM enters per gram *present* per hour, specific formate uptake is a function of growth rate:

$$q = \frac{\mu}{Y_{\max}} + m_s, \tag{S24}$$

with $q$ the specific formate uptake, $Y_{\max}$ the maximum growth-associated yield and $m_s$ the maintenance coefficient. Regressing the dark rGlyP operating-envelope sweep ($\mu$ free, formate imposed) over the 46 points below the basis change at $\mu \approx 0.19$ h$^{-1}$ on Eq. S24 returns $r^2 = 1.000000$ with

$$m_s = 1.714 \text{ mmol gDW}^{-1}\text{ h}^{-1}, \qquad Y_{\max} = 4.637 \text{ gDW mol}^{-1} \text{ formate}. \tag{S25}$$

The fit is exact because the LP is piecewise linear in the imposed substrate, so the decomposition recovers the maintenance terms rather than approximating them.

The intercept is the zero-growth maintenance flux: at $\mu = 0$ the model consumes 2.0 mmol formate gDW$^{-1}$ h$^{-1}$. NGAM plus the proton leak (1.0 mmol $H^+$ gDW$^{-1}$ h$^{-1}$, ≈0.25 mmol ATP-equivalent at 4 $H^+$/ATP) require 3.25 mmol ATP gDW$^{-1}$ h$^{-1}$, supplied by 2.0 mmol formate at an effective P/O of ≈1.6. Recovering $m_s$ therefore confirms the implementation rather than testing it:

3 mmol ATP $gDW^{-1}$ $h^{-1}$ of NGAM divided by 7/4 ATP per formate gives 1.714, the fitted value to three decimals. Under the earlier proton budget (since corrected; Complex I uncounted, chain capped at 6 $H^+$ per formate) the same identity gave $3/(6/4) = 2.0$; the intercept tracks the stoichiometry exactly.

$Y_{\max}$ is growth-associated and comparable with measurement: 4.637 against $4.52 \pm 0.17$ gDW $mol^{-1}$ in chemostat[24], within reported uncertainty. This carries the same information as the overall-yield benchmark of Section S12 rather than adding independent evidence, since both derive from the same maintenance terms; it is reported because the decomposition separates contributions the single yield figure combines. Reproduce with `PaperScripts/metabolic_audits/pirt_decomposition.py`.

The reconstruction extensions (pathway-specific assimilation[25], rhodopsin pump, proton leak, solver settings) are summarised in the main Methods. The proton leak is 1.0 mmol $gDW^{-1}$ $h^{-1}$ (∼20–40% of NGAM dissipated as PMF leakage). LPs are solved with COBRApy[26] over optlang[27] and GLPK (feasibility $10^{-6}$, optimality $10^{-7}$). Because the LPs are typically degenerate, a lexicographic tie-break pins the primary objective within a small window and a secondary minimisation (proteome draw, then respiratory $O_2$ uptake) selects a unique distribution; reported shadow prices are retained from the *primary* LP.

##### S4.2 GECKO enzyme-capacity constraint

Capacity limits follow GECKO[28] (summarised in the main Methods). For each enzyme-catalysed reaction $j$,

$$v_j \;\le\; \sigma_j \, k_{\mathrm{cat},j} \, E_j, \tag{S26}$$

with $k_{\mathrm{cat},j}$ in $s^{-1}$, $E_j$ the molar enzyme allocated ($\mathrm{mmol\,gDW^{-1}}$), and $\sigma_j \in (0,1]$ an apparent saturation factor[29]. Writing the allocated protein mass as $m_j = \mathrm{MW}_j \, E_j$ and inverting gives the minimum mass sustaining a given flux,

$$m_j \;\ge\; \frac{\mathrm{MW}_j}{\sigma_j \, k_{\mathrm{cat},j} \, 3600} \, v_j \;=\; c_j \, v_j, \qquad c_j \;=\; \frac{\mathrm{MW}_j}{\sigma_j \, k_{\mathrm{cat},j} \, 3600}, \tag{S27}$$

where $c_j$ is the proteome *draw coefficient.* Draws aggregate against one shared pool,

$$\sum_j \frac{\mathrm{MW}_j}{\sigma_j \, k_{\mathrm{cat},j} \, 3600} \, v_j \;\le\; P_{\mathrm{total}} \, f_{\mathrm{enzyme,eff}}(\mu), \tag{S28}$$

implemented as a pseudo-metabolite `prot_pool` consumed in proportion $c_j$ and supplied by a bounded draw reaction. Reactions outside the kinetic mapping draw nothing and are unconstrained by Eq. (S28). A numerical floor excludes effective turnovers below $10^{-3}$ $s^{-1}$ to prevent ill-conditioned draw coefficients. No mapped reaction in either pathway hits this floor: rebuilding and re-solving both fixed- and free-growth models at floors of $10^{-4}$, $10^{-3}$ and $10^{-2}$ $s^{-1}$ leaves formate demand and maximum growth unchanged to numerical precision (`PaperScripts/Supplementary/data/audit_kcat_floor.py`; `Results/Supplementary/kcat_floor_audit.csv`).

##### S4.3 Reversible reactions and the Haldane factor

Reversible reactions are split into irreversible halves with independent draws, as summarised in the main Methods ($\kappa = 0.1$[30]). Reactions whose SBML "reverse" direction is physiological during formatotrophy are exempted ($k_{\mathrm{cat}}^{-} = k_{\mathrm{cat}}^{+}$): the gluconeogenic/CBB block in the carbon-fixing direction, and the rGlyP $C_1$/THF block (formate activation, methenyl-cyclohydrolase, methylene-THF dehydrogenase, glycine cleavage, serine hydroxymethyltransferase) in the biosynthetic direction. Directionality is pinned where thermodynamic consistency requires it (e.g. ALCD19 restricted to its favourable reduction direction). Both $\kappa$ and the exemption membership are modelling choices, and their effect on the pathway comparison is bounded by the sensitivity analyses in Sections S8 and S14.

##### S4.4 Turnover database and curated heterologous enzymes

The database of 1317 entries (Table S16; 1306 TurNuP predictions[31], three heterologous rGlyP enzymes and five reaction-level complex entries) is summarised in the main Methods.

The ATP synthase entry takes $k_{\mathrm{cat}} = 270\ \mathrm{s}^{-1}$, the maximal turnover for the *E. coli* $F_oF_1$ enzyme in the ATP-*synthesis* direction[32]; the faster literature figures are hydrolysis-direction rates and are not used. Because this turnover sets how much ATP a unit of proteome can make from the proton-motive force, the choice of direction is reported rather than left to the database.

The three heterologous enzymes absent from the host genome have no TurNuP prediction: methenyl-THF cyclohydrolase (Fch, 120 $\mathrm{s}^{-1}$; $V_{\mathrm{max}}$-derived from *M. extorquens* FchA, UniProt Q49135[25,33]), formate-activating GAR transformylase (GART, 37.6 $\mathrm{s}^{-1}$, from *E. coli* PurT[34]), and formate–THF ligase (Ftl, 116 $\mathrm{s}^{-1}$, from *M. extorquens* AM1, specific activity 117 U $\mathrm{mg}^{-1}$, subunit mass 59.4 kDa[35]). Live-reaction screening was disabled: capacity limits can reroute flux through reactions dormant in the unconstrained optimum, so constraining only the active set would understate proteome cost. A further 54 reactions were constrained to zero or given directional bounds (Section S4.9; Table S15).

##### S4.5 Ribosome-coupled resource balance and the free-growth fixed point

The resource-balance closure of Goelzer and Fromion[36] is imposed at sector level (calibration described in the main Methods):

$$f_{\mathrm{enzyme,eff}}(\mu) = \max[0,\ f_{\mathrm{enzyme}} - \min(f_{\mathrm{ribo,max}},\ f_{\mathrm{ribo,0}} + k_{\mathrm{ribo}}\,\mu)]\,, \tag{S29}$$

with $f_{\mathrm{enzyme}} = 0.50$, $f_{\mathrm{ribo,max}} = 0.40$, $f_{\mathrm{ribo}}(\mu) = 0.068 + 0.157\,\mu$[23,37,38], and $P_{\mathrm{total}} = 0.68\ \mathrm{g\,gDW}^{-1}$[23]. At $\mu = 0.126\ \mathrm{h}^{-1}$ these give $f_{\mathrm{ribo}} = 0.0878$, $f_{\mathrm{enzyme,eff}} = 0.4122$ and a pool bound of $\approx$0.280 $\mathrm{g\,gDW}^{-1}$. These calibrate the aggregate native partition only; heterologous rGlyP and rhodopsin allocations remain free variables within the shared pool.

In fixed-growth runs the imposed $\mu$ sets the pool bound directly. In free-growth runs $\mu$ and the pool are mutually dependent, so the coupled problem is solved by damped fixed-point iteration:

$$\mu^{(n+1)} = (1-d)\,\mu^{(n)} + d\,\mu^{*}\big(f_{\mathrm{enzyme,eff}}(\mu^{(n)})\big), \tag{S30}$$

with initial damping $d = 0.5$, terminating at $|\mu^{(n+1)} - \mu^{(n)}| < 10^{-6}$ or 100 iterations. If successive changes increase (oscillation), damping is reduced ($d \leftarrow 0.8\,d$, floored at 0.1). The map is contractive near the solution because $\mu^*$ is non-decreasing in the pool bound while $f_{\mathrm{enzyme,eff}}$ is non-increasing in $\mu$.

##### S4.6 Membrane-area budget

Membrane-embedded enzymes compete for finite inner-membrane area (footprints and structural sources listed in the main Methods):

$$\sum_j a_j\, v_j \;\leq\; A_{\text{cell}} = 2.0 \times 10^{13}\ \mu\text{m}^2\,\text{gDW}^{-1} = 20\ \text{m}^2\,\text{gDW}^{-1}, \tag{S31}$$

with $a_j = \text{footprint}_j\, N_\text{A}/(k_{\text{cat},j}\, 3600)$. The budget derives from the membrane-proteome inventory of Jahn et al.[23], consistent with Zhuang et al.[39], and is implemented through a pseudo-metabolite `membrane_space`. Footprints are $\sim$7 nm$^2$ for rhodopsin monomers, $\sim$1000 nm$^2$ for chlorophyll photosystem complexes, and $\sim$22–70 nm$^2$ for the respiratory complexes ($F_o$ ATP synthase $\sim$22, quinone-linked FDH $\sim$50, Complex I $\sim$70, terminal oxidases $\sim$25–30, Complex II $\sim$25 nm$^2$ [40–45]). Soluble enzymes and transporters remain unpriced, the latter because their aggregate occupancy is not resolved by the reconstruction. The respiratory footprints are order-of-magnitude estimates, so their level is treated as a sensitivity (Section S15).

##### S4.7 Rhodopsin as a physics-coupled GECKO enzyme

Rhodopsin enters the metabolic layer as summarised in the main Methods: a transmembrane pump upper-bounded by $R_\text{pump}$ (Section S1), MW = 27 kDa, 7 nm$^2$ footprint, drawing on both Eq. (S28) and Eq. (S31), with $k_{\text{cat,rho}} = R_\text{pump}$ ($\approx$3.85 $H^+$ s$^{-1}$ at 100 PPFD wild type). This ties expression cost to the same photophysics that sets output: a slower pump must be expressed at proportionally higher abundance, competing more strongly for both budgets. No independent turnover parameter is introduced, so the photosystem incurs no free fitting parameter in the metabolic layer.

##### S4.8 Enzyme-capacity saturation policies

Because intracellular concentrations cannot be estimated from a genome-scale reconstruction, the $\sigma_j$ factors in Eqs. (S26)–(S28) are assigned by policy. Four were evaluated (see also main Methods). *Unconstrained*: no capacity limits (ordinary FBA). *Uniformly saturated*: $\sigma_j = 1$. *Uniform 50%*: $\sigma_j = 0.5$. *EC-class-specific* (ECC, primary): EC 1 (oxidoreductases) 0.7 [NAD(P)H-dependent 0.75, quinone-dependent 0.5, $O_2$-dependent 0.35], EC 2 0.6, EC 3 0.7, EC 4 0.5, EC 5 0.6, EC 6 0.4 and unmapped 0.5, approximating typical *in vivo* patterns[29]. Per-reaction overrides include RuBisCO at $\sigma = 0.8$ because formate oxidation floods the cell with $CO_2$, raising carboxylase substrate well above $K_m$; the specific value is a qualitative judgement rather than a back-calculated one, since intracellular $CO_2$ was not independently estimated here.

The pathway ranking is invariant across all four policies: rGlyP requires less formate than CBB through low-to-moderate PPFD, the fixed-growth advantage is preserved, and only the absolute pump-rate thresholds shift (Section S10). Since the factors are not individually calibrated, this cross-policy invariance supports the ranking and shadow-price structure.

##### S4.9 Constrained and disabled reactions

Reactions were assigned zero-flux bounds or directional caps reflecting formate-only aerobic growth without exogenous amino acids or nucleosides (Table S15). Five categories: absent-medium transport, physiologically unidirectional reactions, futile/thermodynamically infeasible cycles, reactions inactive in H16 under these conditions, and exchange fluxes for metabolites neither supplied nor secreted. Every constraint was verified by screening FBA: removing any one did not change the optimal growth rate, confirming the disabled set encodes the medium without restricting the attainable flux space.

**Transhydrogenase and cofactor policy.** The soluble NAD(P) transhydrogenase (NADTRHD) is fixed at zero flux to prevent rhodopsin-derived PMF from being converted implicitly into NADPH (see also main Methods). The proton-coupled THD2 is retained but treated as PMF-coupled in the thermodynamic validation and flux audit. A stress test reopening NADTRHD to 100 mmol $gDW^{-1}$ $h^{-1}$ gives the same fixed-growth formate demand, with zero NADTRHD flux across dark, falsification-point and high-light cases (Table S28). A validated *in vivo* transhydrogenase route would be a different physiology, but it is not a hidden optimum in the current solutions.

#### S5 Reactor optics: two-flux Schuster model and radial extension

The problem is to convert per-molecule photophysics into a volume-averaged metabolic response for a dense, self-shading culture. Radial geometries reuse the flat-panel solution through an exact power transformation. Implementation: `Bin/PhysBin/ReactorOptics/optical_properties.py`, `reactor_model.py`, `cylindrical_model.py`.

##### S5.1 Decomposition of the suspension optical properties

The suspension is characterised by two mass-specific coefficients in $m^2\,gDW^{-1}$, absorption $a(\lambda)$ and scattering $s(\lambda)$; multiplying either by the biomass concentration $X$ ($gDW\,m^{-3}$) gives attenuation per unit depth. Absorption is the sum of a non-pigment background and rhodopsin self-shading,

$$a(\lambda) = \kappa_{\text{abs,bg}}(\lambda) + \kappa_{\text{abs,rho}}(\lambda), \qquad s(\lambda) = \kappa_{\text{scat}}(\lambda). \tag{S32}$$

**Mie scattering.** Scattering by ∼1–2 $\mu$m gram-negative cells is a power law anchored at $\lambda_{\text{ref}} = 550$ nm,

$$\kappa_{\text{scat}}(\lambda) = \kappa_{\text{scat,ref}} \left(\frac{\lambda}{\lambda_{\text{ref}}}\right)^{-p}, \qquad \kappa_{\text{scat,ref}} = 0.8\ m^2\,gDW^{-1}, \quad p = 1, \tag{S33}$$

adapted from radiative-transfer practice for bacterial suspensions[46,47]. This is total cuvette-measured scattering; the two-flux closure below separates forward from back-scattered light.

**Background biomass absorption.** *C. necator* H16 is unpigmented and no organism-specific visible absorption spectrum was available. The production calculation therefore uses a small wavelength-independent background,

$$\kappa_{\text{abs,bg}}(\lambda) = 0.02\ \text{m}^2\,\text{gDW}^{-1}, \tag{S34}$$

within the bacterial radiative-transfer range[46,47]. A 512-draw optical audit varied the scattering coefficient (0.3–2.0 $\text{m}^2\,\text{gDW}^{-1}$), scattering exponent (0.5–1.5), backscatter fraction (0.008–0.015), background magnitude, rhodopsin bandwidth and abundance. With a flat background, the 520:568 nm low-activation pump proxy remained 1.045–1.058 at the reported SOTA areal depth. Replacing it by a deliberately broad exponential background range widened that ratio to 0.676–1.024; wavelength-dependent host absorption can therefore erase the small blue-shift ordering. This does not affect the pathway comparison or geometry ranking, which use the same optical field, and it reinforces the main result that the efficiency optimum is a broad green region rather than a resolved blue-shift target (`PaperScripts/Supplementary/data/audit_optical_inputs.py`; `Results/Supplementary/optical_input_audit_summary.csv`).

**Rhodopsin self-shading.** The expressed rhodopsin has a Gaussian band centred on its own $\lambda_{\max}$,

$$\varepsilon(\lambda) = \varepsilon_{\max} \exp\left[-\tfrac{1}{2}\left(\frac{\lambda-\lambda_{\max}}{\sigma_\lambda}\right)^2\right], \qquad \sigma_\lambda = \frac{\text{FWHM}}{2\sqrt{2\ln 2}}, \quad \text{FWHM} = 100\ \text{nm}, \tag{S35}$$

with peak molar extinction $\varepsilon_{\max} = 63{,}000\ \text{M}^{-1}\text{cm}^{-1}$ at 568 nm[48], converted to a mass-specific natural-log coefficient through the Beer–Lambert unit chain,

$$\kappa_{\text{abs,rho}}(\lambda) = \varepsilon(\lambda)\, n_{\text{rho}}\, \underbrace{\ln(10)\,\frac{1\ \text{cm}}{100\ \text{cm}\,\text{m}^{-1}}\,\frac{1\ \text{m}^3}{1000\ \text{L}}}_{=\ln(10)/10\,=\,0.2303}, \tag{S36}$$

where $n_{\text{rho}}$ is the abundance in $\text{mol}\,\text{gDW}^{-1}$ and 0.2303 converts base-10 to base-$e$ and $\text{M}^{-1}\text{cm}^{-1}$ to $\text{m}^2\text{gDW}^{-1}$. A reference $n_{\text{rho}} = 5.0 \times 10^{-6}\ \text{mol}\,\text{gDW}^{-1}$ is used in standalone ablations; the integrated efficiency figures substitute the LP-derived allocation. Rhodopsin dominates absorption (∼94% at $\lambda_{\max}$). Self-shading uses the physical extinction cross-section; the empirical $\sigma_{\text{eff}}/\sigma_0 = 4.5$ scale enters only the per-molecule pump rate, not the bulk optics.

##### S5.2 Two-flux (Schuster) radiative transfer in a flat panel

The two-flux approximation[47,49] avoids the over-prediction of a single Beer–Lambert exponential (∼99% of scattered light is forward directed) by tracking forward and backward irradiances exchanging only through the back-scattered fraction. For collimated front illumination of a panel of depth $L$,

$$\begin{aligned}\frac{\mathrm{d}I^+}{\mathrm{d}z} &= -(a+b\,s)\,X\,I^+ + b\,s\,X\,I^-,\\ \frac{\mathrm{d}I^-}{\mathrm{d}z} &= +(a+b\,s)\,X\,I^- - b\,s\,X\,I^+,\end{aligned} \tag{S37}$$

with $I^+(0) = I_0$ and $I^-(L) = 0$. Here $b \approx 0.01$ is the backscatter fraction[47]. Defining

$$\alpha = a + b\,s \quad \text{(forward attenuation)}, \qquad \beta_{\text{s}} = b\,s \quad \text{(backscatter coupling)}, \tag{S38}$$

the actinic flux is $G(z) = I^+(z) + I^-(z)$ (the chromophore absorbs regardless of propagation direction). Seeking solutions $\propto e^{\pm\delta X z}$ gives

$$\delta = \sqrt{\alpha^2 - \beta_{\text{s}}^2} = \sqrt{a\,(a + 2\,b\,s)}, \tag{S39}$$

and applying the boundary conditions yields the Cornet–Schuster solution[47,49]

$$I^+(z) = I_0\,\frac{\delta\cosh[\delta X(L-z)] + \alpha\sinh[\delta X(L-z)]}{\delta\cosh(\delta XL) + \alpha\sinh(\delta XL)},$$
$$I^-(z) = I_0\,\frac{\beta_\mathrm{s}\sinh[\delta X(L-z)]}{\delta\cosh(\delta XL) + \alpha\sinh(\delta XL)}, \tag{S40}$$

whose sum gives the compact form

$$G(z,\lambda) = I_0\,\frac{2\,\delta}{(\delta+\alpha)\,e^{\delta Xz} + (\delta-\alpha)\,e^{-\delta Xz}}. \tag{S41}$$

Because $\delta \ll \kappa_\mathrm{total} = a + s$, only true absorption and back-scattered loss attenuate the beam; $\delta$ varies by only $\sim$4% across 520–600 nm, so wavelength affects penetration far less than it affects the per-molecule pump rate.

**Numerical conditioning and refinements.** For $\delta XL > 500$ the hyperbolic functions overflow and the implementation switches to the absorption-dominated asymptote $G(z) \approx I_0\,e^{-\delta Xz}$, with the backward stream scaled by $\beta_\mathrm{s}/(\delta+\alpha)$. An optional specular ($R = 0.95$) or Lambertian ($R = 0.85$) back-wall reflector is added for flat panels as $I_\mathrm{refl}(z) = R\,I^+(L)\,e^{-\alpha(L-z)}$, exact when $\beta_\mathrm{s} \ll \alpha$ and not applied to radial geometries. A volumetric pure-water absorption term (Pope and Fry values, linearly interpolated) is negligible (<1.3% over a 5 cm path) in 520–600 nm but rises $\sim$6× across the band and steeply beyond 600 nm, giving a systematic red-side penalty in dilute reactors.

Discrete-ordinates ($S_N$ with $\delta$-Eddington truncation) and vectorised Monte Carlo solvers are implemented as cross-checks; the two-flux closed form is used for all production figures.

#### S5.3 Radial geometries via the power transformation

In an axially uniform cylinder the $1/r$ divergence terms are removed by transforming to radial *power* fluxes

$$P^\pm(r) = 2\pi r\,I^\pm(r), \tag{S42}$$

the photon power crossing the shell at $r$ per unit axial length. Substituting cancels the $1/r$ terms:

$$\frac{\mathrm{d}P^+}{\mathrm{d}r} = -\alpha\,X\,P^+ + \beta\,X\,P^-,$$
$$\frac{\mathrm{d}P^-}{\mathrm{d}r} = +\alpha\,X\,P^- - \beta\,X\,P^+, \tag{S43}$$

which is formally identical to Eq. (S37), so the solution (S40) applies directly to $P(r)$ along the radial depth measured inward from the illuminated surface (with $P^+(0) = 2\pi r_\mathrm{illum} I_0$ and $P^- = 0$ at the far boundary). Inverting the transform,

$$G(r) = \frac{P^+(r) + P^-(r)}{2\pi r}, \tag{S44}$$

reintroduces the $1/r$ geometric concentration factor. Under external illumination light converges toward smaller $r$, *focusing* it toward the axis and partially compensating optical attenuation; under internal illumination light diverges, *spreading* and compounding attenuation.

Four configurations share the same optics, pump kinetics and metabolic interpolators: solid cylinder with external illumination ($R = 2.5$ cm); cylinder with internal lamp ($r_1 = 0.5$ cm, $R = 2.5$ cm); and annular reactors with external and with internal illumination ($r_1 = 1.0$ cm, $r_2 = 3.5$ cm). The externally illuminated cylinder delivers the largest volume-averaged flux ($\sim 2\times$ the flat panel at matched conditions) and anchors the efficiency closure (Figure S9).

##### S5.4 Volume averaging and coupling to the pump–metabolism model

The field is sampled on 200 depth or radial slices (<0.1% quadrature error) and each slice mapped to a local response. At local irradiance $G$ the per-rhodopsin absorption rate is $k_{\mathrm{abs}} = G\,N_{\mathrm{A}}\,10^{-6}\,\sigma_{\mathrm{eff}}$ with $\sigma_{\mathrm{eff}} = 4.5\,\sigma_0$, and

$$R_{\mathrm{pump}}(G) = \frac{1 - \exp[-k_{\mathrm{abs}}(G)\,\Phi_{\mathrm{QY}}\,\tau]}{\tau}, \tag{S45}$$

converted to a biomass-specific flux by $r_{\mathrm{pump}}^{\mathrm{gDW}} = R_{\mathrm{pump}}\,n_{\mathrm{rho}}\,(3600 \times 1000)$ in $\mathrm{mmol\,H^+\,gDW^{-1}\,h^{-1}}$ (the Avogadro factor cancels since $R_{\mathrm{pump}}$ is per molecule). The local flux indexes the fixed-growth metabolic sweep via a monotone PCHIP interpolator.

Two mixing limits bracket hydrodynamics:

$$\begin{aligned} \text{stratified:} \quad \langle Y\rangle &= \frac{\int Y\big(G(z)\big)\,w(z)\,\mathrm{d}z}{\int w(z)\,\mathrm{d}z}, \\ \text{perfectly mixed:} \quad \langle Y\rangle &= Y\big(\langle G\rangle\big), \qquad \langle G\rangle = \frac{\int G(z)\,w(z)\,\mathrm{d}z}{\int w(z)\,\mathrm{d}z}, \end{aligned} \tag{S46}$$

by trapezoidal integration, with geometric weight $w = 1$ for the flat panel and $w(r) = r$ for the radial element. Because the pump-rate-to-yield map is concave, Jensen's inequality makes the perfectly mixed average an upper bound on the stratified one. The fixed-growth mode used throughout corresponds to the perfectly mixed limit; geometry ranking depends only on the light-field integral and is independent of the mixing assumption.

##### S5.5 Regimes and modelling limitations

The optical penalty is pathway-independent because rGlyP and CBB share the same local-flux-to-pump map. Spectral advantage peaks at low density ($\sim$0.1–0.2 $\mathrm{gDW\,L^{-1}}$) and collapses above $\sim$1 $\mathrm{gDW\,L^{-1}}$. The externally illuminated cylinder under perfect mixing widens the useful window most and anchors the efficiency analysis.

The broad optical-input audit above leaves the pathway comparison and geometry ranking intact but makes the fine wavelength ordering conditional on the host absorption spectrum. The back-wall reflector is a closed-form flat-panel correction; it does not enter the externally illuminated cylinder used for the headline efficiency.

##### S5.6 Volume-averaged light delivery

The delivery fraction

$$f_{\text{del}} = \frac{\langle \text{PPFD} \rangle}{\text{PPFD}_0} \tag{S47}$$

is the volume-averaged actinic flux normalised to the incident surface flux. It depends only on geometry, biomass concentration and wavelength through the two-flux solution, not on the electrocatalysis scenario or efficiency accounting.

For externally illuminated cylindrical and annular vessels, geometric focusing can raise $f_{\text{del}}$ above unity at low optical depth (∼1.9 for the cylinder, ∼1.4 for the annulus), because the illuminated cross-section contracts inward while photon number is conserved. The flat panel has no focusing term and $f_{\text{del}} \leq 1$ throughout.

At ∼2 gDW $L^{-1}$, $f_{\text{del}}$ is 0.90 for the external cylinder, 0.79 for the annulus and 0.65 for the flat panel, whereas the internally illuminated bag has fallen to 0.09. This ordering motivates carrying the external curved geometries into the process optimisation.

Note that $\alpha$ is used elsewhere in this section for the two-flux attenuation coefficient $\alpha = a + b\,s$ (Eq. S38), a distinct quantity from the delivery fraction defined here.

#### S6 Solar-to-biomass efficiency and bounded design optimisation

Headline parameters: Table S22; constrained-optimal designs: Table S17; Pareto points: Table S29. Diagnostic implementation: `Bin/PhysBin/ReactorOptics/process_efficiency.py` and `PaperScripts/reactor_geometry/data/optimize_design.py`; whole-collector implementation: `PaperScripts/reactor_geometry/data/optimize_hybrid_dichroic_led.py`.

##### S6.1 Energy-flow definition of $\eta$

The diagnostic route efficiency divides biomass chemical energy by the solar input assigned to two branches: carbon-and-reductant ($CO_2$ electrolysed to formate via photovoltaic electricity) and photon (rhodopsin-band sunlight delivered to the reactor). At the fixed setpoint,

$$\begin{aligned} \eta &= \frac{\dot{E}_{\text{biomass}}}{\dot{E}_{\text{formate,PV}} + \dot{E}_{\text{photon,del}}}, \\ \dot{E}_{\text{biomass}} = v_{\text{bio}}\, H_{\text{bio}}, &\qquad \dot{E}_{\text{formate,PV}} = \frac{\dot{n}_{\text{formate}}\, E_{\text{elec}}}{1000\, \eta_{\text{PV}}}, \end{aligned} \tag{S48}$$

with $v_{\text{bio}} = 0.126$ $\text{g}\,\text{gDW}^{-1}\,\text{h}^{-1}$, biomass higher heating value $H_{\text{bio}} = 22.4$ $\text{kJ}\,\text{g}^{-1}$, $\dot{n}_{\text{formate}}$ the LP-minimised formate uptake, $E_{\text{elec}} = E_{\text{synth}} + E_{\text{cap}}$ the combined electrical demand for formate synthesis and $CO_2$ capture per mole formate, and $\eta_{\text{PV}}$ the photovoltaic efficiency. Maintenance is already embedded in the LP through GAM and NGAM. Equation S48 defines the diagnostic *route* efficiency (Table S10). The headline *whole-collector* efficiency, also called the closed boundary, instead counts all AM1.5G sunlight incident on the collection aperture required to meet the modelled electrical and photon loads, with no credit for exported electricity. The aperture is sized within the optimisation below. This is a conversion efficiency at steady design irradiance; it excludes seasonal operation, collector spacing and other site-area requirements, so it is not a prediction of annual land productivity. For the bounded optimisation, biomass output per illuminated area is expressed via density CDW ($\mathrm{g\,L^{-1}}$) and depth $V/A$ (m):

$$\dot{m}_{\mathrm{bio}} = \mu\,\mathrm{CDW} \times 1000 \times (V/A) \quad [\mathrm{g\,m^{-2}\,h^{-1}}], \qquad \dot{E}_{\mathrm{biomass}} = \dot{m}_{\mathrm{bio}}\,H_{\mathrm{bio}}. \tag{S49}$$

The formate branch charges $\dot{m}_{\mathrm{bio}}/Y$ with yield $Y$ (g CDW per mol formate); Faradaic and overpotential losses are folded into $E_{\mathrm{elec}}$ ($\eta_{\mathrm{electrolysis}}$ is unity to avoid double-counting). The photon branch is $\dot{E}_{\mathrm{photon,del}} = \mathrm{PPFD}\,(hcN_{\mathrm{A}}/\lambda)$ per unit area, using monochromatic photon energy rather than a broadband PAR factor (up to ∼19% error at the blue end).

In the main scenario these are *dichroic solar* photons: a wavelength-selective optic (0.90 efficiency, 100 nm FWHM) redirects band-matched sunlight at native photon energy, so no PV or LED loss applies. Two LED-based alternatives (full solar→PV→LED chain) are retained as sensitivity branches; neither enters headline numbers. The non-rhodopsin spectral fraction is available to the PV/formate branch or allocated outside this denominator.

#### S6.2 Formate-saving efficiency and electrolyser scenarios

A complementary efficiency isolates the photosynthetic contribution by crediting only the electrolyser energy displaced by light relative to the dark reference,

$$\eta_{\mathrm{save}} = \frac{\Delta\dot{n}_{\mathrm{formate}}\,E_{\mathrm{elec}}/(1000\,\eta_{\mathrm{PV}})}{\dot{E}_{\mathrm{photon,del}}}, \qquad \Delta\dot{n}_{\mathrm{formate}} = \dot{n}_{\mathrm{formate,dark}} - \dot{n}_{\mathrm{formate,light}}, \tag{S50}$$

measuring the marginal value of light forcing. Two electrolyser scenarios bracket present and near-thermodynamic technology. *SOTA*: $E_{\mathrm{synth}} = 500\ \mathrm{kJ\,mol^{-1}}$ (≈2.6 V, ≈51% electrical-to-chemical efficiency on the formate HHV basis); Faradaic efficiencies exceed 90% at lab scale and ∼80% in scaled cells[50], with the assumed electrosynthesis demand including conversion losses. Paired with $\eta_{\mathrm{PV}} = 0.241$[51]. *Theoretical*: $E_{\mathrm{synth}} = 270\ \mathrm{kJ\,mol^{-1}}$ (≈254.6 $\mathrm{kJ\,mol^{-1}}$ formate HHV plus modest overpotential) with Shockley–Queisser $\eta_{\mathrm{PV}} = 0.330$. For capture-charged calculations, $E_{\mathrm{cap}} = 419.9$ and 19.16 $\mathrm{kJ\,mol^{-1}}$ are added to these electrosynthesis demands, respectively, giving $E_{\mathrm{elec}} = 919.9$ and 289.16 $\mathrm{kJ\,mol^{-1}}$. Capture-free diagnostics omit that additional term and are labelled accordingly.

#### S6.3 Three physical reactor limits replace an areal-biomass cap

An earlier, arbitrary areal-biomass cap is replaced by three physically interpretable limits. *Gas–liquid $O_2$ transfer*: $\mathrm{OUR} = q_{O_2}\,\mathrm{CDW} \le \mathrm{OTR_{max}} = 100\ \mathrm{mmol\,L^{-1}\,h^{-1}}$, a conservative mid-range for a light-transmitting vessel (∼50–300 in well-aerated reactors[52]); rhodopsin relieves this bottleneck by displacing respiration. *Substrate feed*: volumetric formate uptake ≤ 100 $\mathrm{mmol\,L^{-1}\,h^{-1}}$, bounding the light-saturated optimum. *Solar concentration*: surface PPFD ≤ 3× unconcentrated AM1.5G rhodopsin-band PPFD at the design wavelength after 0.90 dichroic efficiency (100 nm window).

The first two are volumetric (density-dependent); the third bounds surface optical input. At the external-reactor optima, solar concentration binds, with $O_2$ uptake ∼0.04 and formate uptake ∼14 $\mathrm{mmol\,L^{-1}\,h^{-1}}$ at the SOTA rGlyP cylinder optimum.

##### S6.4 Order-of-magnitude cost of the dichroic and concentrating optics

The 3× concentration limit is active at every SOTA optimum (Table S27; `concentration_factor`$\approx$ 3.0), so the design relies on two uncosted optical elements: the dichroic (Methods) and a low-concentration non-tracking collector. No published cost exists for this combination; what follows is an order-of-magnitude bracket.

Non-tracking 2–5× collectors cost \$50–300 $m^{-2}$ [53]. The dichroic filter is the recurring cost obstacle[54]; we bound it at \$200–2,000 $m^{-2}$ (mass-manufactured to small-batch deposition), dominating the uncertainty.

Summed: \$250–2,300 $m^{-2}$ of aperture. At the SOTA design's ≈3.0 $m^2$ aperture per $m^2$ reactor (45% occupancy), this is ≈\$750–6,900 $m^{-2}$ of reactor footprint, spanning below to several times the ∼\$1,600 $m^{-2}$ installed capital of a $CO_2$-to-formate electrolyzer[55] (electrode-area basis, not like-for-like).

##### S6.5 Six-variable bounded optimisation

For every discrete geometry, pathway and electrolyser combination, $\eta$ is maximised over $\lambda \in [450, 600]$ nm, CDW $\in [0.05, 10]$ $\mathrm{g\,L^{-1}}$, surface PPFD $\in [50, 50{,}000]$ $\mu\mathrm{mol\,m^{-2}\,s^{-1}}$, $V/A \in [0.005, 0.1]$ m, $f_{\mathrm{mem}} \in [0, 1.0]$, and dichroic transmission $t \in [0, 1]$. CDW, PPFD and $V/A$ are searched on $\log_{10}$ scale; each evaluation runs the full physics–metabolism–optics pipeline. Global search uses differential evolution (population 32, up to 250 iterations, mutation $[0.4, 1.0]$, recombination 0.7, deferred updating, polishing disabled), followed by L-BFGS-B polish accepted only if it improves the objective.

Infeasible designs receive a large positive objective ($\gtrsim 10^3$, rising with overshoot). Reported variables are rounded conservatively: CDW is floored to four decimal places ($\mathrm{g\,L^{-1}}$) and $V/A$ standard-rounded to six decimal places (m), so a rounded design cannot exceed the volumetric limits (`_round_feasible_loading` in `PaperScripts/reactor_geometry/data/optimize_design.py`); PPFD, $f_{\mathrm{mem}}$ and dichroic $t$ are reported at the optimiser's native output precision. The headline point estimates charge $CO_2$ capture and optimise the dichroic split (see "Why the LED branch is never used" below).

On the closed boundary at 45% occupancy, SOTA $\eta \approx 3.29\%$ and Theoretical $\approx 10.89\%$ for rGlyP; at full-budget ceiling 3.74% and 10.90%, against 3.06% and 9.22% for CBB. The route-level counterpart, which does not count the whole collection aperture, is tabulated in Table S27(a).

For the formate-cost sweep in Figure 7g, total electrical demand is varied from 289.16 kJ $mol^{-1}$ (the synthesis-plus-capture thermodynamic minimum) to $1.5 \times 919.9$ kJ $mol^{-1}$. PV efficiency remains at the current-technology value, so only the energetic cost of formate changes. Each pathway and optical case is re-optimised along the sweep. The light-enabled result is $\max(\eta_{\mathrm{dark}}, \eta_{\mathrm{illuminated}})$, representing a plant free to omit the light-harvesting system. Shading propagates the same PV, biomass-energy, pump, metabolic, solar-delivery and transport uncertainties used for the headline calculation through the selected design at each cost. The generator supports either the model-fit 70% Kin4B8 antenna used in the main panel or retinal alone.

**Carotenoid-antenna sensitivity.** A separate calculation adds a carotenoid antenna to the 45%-occupancy rGlyP external cylinder while leaving the direct retinal band unchanged. Two measured three-peak envelopes are used: 457, 487 and 521 nm for the salinixanthin bands of natural xanthorhodopsin (∼40% carotenoid-to-retinal transfer)[56], and 432, 457 and 487 nm for lutein bound to the freshwater xanthorhodopsin Kin4B8, whose peaks are blue-shifted from salinixanthin's. The Kin4B8 envelope is used for two distinct transfer cases: a directly measured ∼50% value for 3-hydroxylated xanthophylls[57], and a model-derived ∼70% zeaxanthin/lutein estimate fitted to measured transfer kinetics[58]. Zeaxanthin and lutein are structural isomers with equivalent binding and transfer efficiency in Kin4B8[57], so the measured lutein envelope stands in for zeaxanthin as well; zeaxanthin's own bound peak positions were not located in the accessible literature. Peak widths and relative amplitudes for the Kin4B8 envelope are carried over from the salinixanthin fit (shape only; not independently measured for lutein-Kin4B8). Antenna absorption overlapping the 100-nm retinal collection window is removed before integration so that no solar photon is counted twice. Off-band absorbed photons are removed from the photovoltaic complement, multiplied by the stated transfer efficiency and by a delivery factor that brackets coupling and additional short-wavelength loss, and supplied to the existing pump model as retinal-equivalent excitations. This is an equivalent-photon sensitivity, not a second two-flux pigment model.

A further scenario treats the antenna's spectral position as a free variable: all three peak centres are shifted together by an additional optimised offset (bounded so every peak stays inside $[380, 560]$ nm), at the model-fit 70% transfer estimate and ideal delivery. This asks where a freely tunable antenna would want to sit, as an upper bound distinct from any single measured pigment's actual position. Figure S11 maps the complementary interaction between retinal wavelength and transfer efficiency around the fixed-position Kin4B8 optimum.

Wavelength, density, surface PPFD, depth, dichroic transmission and (for the free-wavelength row) the envelope shift were re-optimised for each row with four independent differential-evolution runs (population 32, up to 250 iterations) and L-BFGS-B polishing. The zero-antenna controls recover the headline calculation to numerical precision. The antenna envelopes use primary-study peak locations, but relative peak heights are illustrative. No carotenoid biosynthetic, protein-expression or assembly cost is charged; burden thresholds are therefore calculated below for the favourable cases.

Table S4: **Re-optimised whole-collector equivalent-photon sensitivity to a xanthorhodopsin-like carotenoid band.** $CO_2$ capture and the 45% occupancy cap are retained. $f_{ET}$ is carotenoid-to-retinal transfer; $f_{del}$ is relative antenna-band delivery after dichroic loss; changes are relative to retinal only. Rows 3–4 use published ∼40% salinixanthin transfer, row 6 directly measured ∼50% Kin4B8 transfer, and rows 7–8 a model-fit ∼70% estimate. Rows 1–5 use the salinixanthin envelope; rows 6–8 the measured blue-shifted Kin4B8 envelope, with row 8 additionally shifted. The band is not an explicit two-flux pigment, and assembly costs are excluded.

| Case | $f_{ET}$ | $f_{del}$ | SOTA $\eta$ (%) | $\Delta\eta$ (pp) | Limit $\eta$ (%) | $\Delta\eta$ (pp) |
|---|---|---|---|---|---|---|
| Retinal only | 0 | 0 | 3.288 | 0 | 10.888 | 0 |
| Conservative antenna | 0.20 | 0.50 | 3.218 | −0.071 | 10.671 | −0.217 |
| Measured transfer, partial delivery | 0.40 | 0.75 | 3.260 | −0.028 | 10.679 | −0.209 |
| Salinixanthin transfer, ideal delivery [56] | 0.40 | 1.00 | 3.305 | +0.016 | 10.682 | −0.206 |
| High transfer, ideal delivery (hypothetical) | 0.60 | 1.00 | 3.396 | +0.108 | 10.687 | −0.201 |
| 3-hydroxyxanthophyll transfer, ideal delivery [57] | 0.50 | 1.00 | 3.381 | +0.093 | 10.694 | −0.194 |
| Kin4B8 zeaxanthin/lutein model-fit transfer, ideal delivery [58] | 0.70 | 1.00 | 3.524 | +0.235 | 10.698 | −0.190 |
| Free-wavelength, 70% transfer, ideal delivery | 0.70 | 1.00 | 3.524 | +0.236 | 10.756 | −0.132 |

Rows 6 and 7 use distinct evidence levels on the measured, blue-shifted Kin4B8 envelope rather than a hypothetical ceiling on the salinixanthin envelope: 3-hydroxylated xanthophylls (zeaxanthin, lutein, nostoxanthin) bound to the freshwater xanthorhodopsin Kin4B8 reach ∼50% transfer by transient-absorption/CD spectroscopy [57], whereas a QM/MM plus excitonic model of the same zeaxanthin/lutein–Kin4B8 pairing, fitted to measured transfer times, gives the separate ∼70% estimate [58]. Both exceed the ∼40% salinixanthin figure rows 1–5 are anchored to [56]. At the state of the art the model-fit 70% case gives the largest gain in the table, +0.235 percentage points (3.288 → 3.524%); the directly measured 50% case adds 0.093 points. For the 70% Kin4B8 case, 6.67% is the optical-gain break-even assembly burden, not a validated allowable biological burden; the corresponding break-even values are 2.74% for the 50% Kin4B8 case and 0.49% for measured salinixanthin transfer with ideal delivery.

Row 8 lets the antenna's spectral position optimise freely at the model-fit 70% transfer efficiency, rather than pinning it to the measured Kin4B8 peaks. At the state of the art the optimiser moves the envelope only −2.9 nm from the measured lutein-Kin4B8 position (peaks at 429.1, 454.1 and 484.1 nm), with no change at the reported precision (3.524%). At the thermodynamic limit it moves the envelope +73.0 nm redder (peaks at 505.0, 530.0 and 560.0 nm), raising the imposed-light result from 10.698% to 10.756% but still leaving it below the 11.15% standard-PV dark design. Natural xanthorhodopsin shows that carotenoid transfer is possible, but adding free carotenoid to the present bacteriorhodopsin is not equivalent: salinixanthin failed to form an antenna complex with bacteriorhodopsin or proteorhodopsin in the primary study [56], and the high-efficiency pairings reported since are specific to xanthorhodopsins (Kin4B8) rather than bacteriorhodopsin. Realising them would require a compatible binding pocket as well as a compatible pigment.

**Active-bound audit.** Every output was checked against its imposed bounds, and membrane allocation reaches its upper bound in every case because the objective increases with allocation. The reported efficiency therefore depends on the assumed allocation limit, quantified for both accounting boundaries in Table S27: panel (a) route and panel (b) closed.

At low membrane caps, PPFD falls to its lower bound and $\lambda$ drifts to a window edge, collapsing to the dark baseline. At the thermodynamic limit the PPFD lower bound (50 $\mu$mol m$^{-2}$ s$^{-1}$) binds once the dichroic closes; relaxing it fifty-fold to 1 $\mu$mol m$^{-2}$ s$^{-1}$ moves $\eta$ by <0.15% relative (10.8899 → 10.9032%), confirming a flat-surface pattern.

The closed-boundary thermodynamic-limit optimum lands on the $V/A$ upper bound of 0.1 m ($V/A^{\star}$ = 0.09999 m at 45% occupancy). A depth sweep with CDW and dichroic split re-optimised at each $V/A$ (capture charged) gives $\eta$ = 10.91, 10.92, 10.98, 10.94, 11.00 and 10.90% at $V/A = 0.0999$, 0.12, 0.15, 0.20, 0.30 and 0.50 m, a spread of 0.10 percentage points over a fivefold range (the density–depth degeneracy below). Areal loading saturates near 376–389 g m$^{-2}$; all other headline optima are interior in $V/A$ (78–93% of the bound).

This flatness sets a resolution limit: five random seeds reproduce $\eta$ to reported precision, but position within this nearly flat region can shift $\eta$ by of order a tenth of a percentage point. The three external geometries span only 0.0009 percentage points at SOTA (3.364, 3.365, 3.365) and 0.0002 at the thermodynamic limit (10.890, 10.890, 10.890). These differences do not resolve a preference among external shapes; the larger effect is external versus internal illumination (1.6 percentage points). Pump cross-section calibration has a separate, larger effect, quantified in Section S6.8.

**Why the LED branch is never used.** The closed boundary allows photovoltaic electricity to drive LEDs, yet the optimiser routes <0.1% of photons that way at every membrane cap and both scenarios, for opposite reasons.

At SOTA, $t^{\star} = 1$ (dichroic fully open) because direct transmission is cheaper than the PV–LED round trip: 0.0067 m$^2$ of aperture per delivered photon-watt directly versus 0.0095 through PV and LEDs (factor 1.4). Replacing the concentration cap with an aperture-only limit leaves $\eta$ unchanged (5.004% vs 5.004%; 5.010% at 10× aperture, $CO_2$ supplied).

At the thermodynamic limit, $t^{\star} \approx 0.029$ (dichroic nearly closed): electrical demand on the aperture (≈803 W m$^{-2}$) dominates photon demand (≈10 W m$^{-2}$) by two orders of magnitude, and the residual direct-transmission fraction supplies photon demand exactly (9.97 vs 9.97 W m$^{-2}$). The LED branch is thus uneconomic at SOTA and unnecessary at the thermodynamic limit.

#### S6.6 Iso-efficiency degeneracy and how the limits break it

In the perfectly mixed limit the optical response depends on density and depth only through their product (areal optical depth), because two-flux attenuation enters as $\delta XL$ with $X \propto$ CDW and $L = V/A$ (Eq. S41), while areal productivity (Eq. S49) and areal formate demand scale identically. $\eta$ is therefore *degenerate* along constant areal optical depth: a thin dense culture and a thick dilute one are energetically equivalent. The reported optimum is one point on an iso-efficiency ridge.

The volumetric limits break this degeneracy: $O_2$-transfer and feed loads depend on density *directly*, not only through areal optical depth, so two designs equal in $\eta$ can differ in feasibility. Dense self-shaded cultures incur the highest aeration load and are excluded first.

##### S6.7 Sensitivity of the optimum

Unless stated otherwise, the mass-transfer and optical-scale diagnostics in this subsection use the route boundary without $CO_2$-capture charge (Table S7); they should not be compared directly with the closed, capture-charged main result. The growth-setpoint audit below is the exception and re-optimises the headline closed boundary. Because light delivery, not aeration or feed, bounds the route diagnostic, its optimum changes by <0.1 percentage points as the $O_2$ limit varies over 20–200 and the feed limit over 50–300 $\mathrm{mmol\,L^{-1}\,h^{-1}}$.

A separate audit uses the headline whole-collector, capture-charged boundary. Re-optimising with each reactor limit relaxed tenfold: concentration binds at SOTA (utilisation 1.00), formate feed at the thermodynamic limit (1.00, concentration 0.02 once the dichroic closes). Relaxing transport limits leaves $\eta$ unmoved (0.000 points for $O_2$, ≤0.001 for feed); only the concentration cap yields anything (≤0.05 points). Aeration runs at 4–30% of the transfer rate (`PaperScripts/reactor_geometry/data/audit_transport_limit_relaxation.py`; `Results/reactor_geometry_wavelength/transport_limit_relaxation.csv`).

Mirrored rescaling of $\sigma_{\mathrm{eff}}$ in both the forward pump map and inverse lookup is an algebraic consistency check: SOTA changes by only 0.14 percentage points (7.06, 7.10, 7.20% at 0.5×–2×). The physical sensitivity instead rescales only the forward map and gives the wider interval quantified in Section S6.8. A ±10% change in GAM shifts formate demand by ∓6.5% and $\eta$ by $\sim \pm 7\%$.

**Sensitivity to the growth setpoint.** All Results use $\mu = 0.126$ $\mathrm{h^{-1}}$, chosen as 70% of the measured 0.18 $\mathrm{h^{-1}}$ maximum formate-growth rate. The current engineered rGlyP strain reaches about 0.063 $\mathrm{h^{-1}}$ (Section S12); lower-rate cases are supplementary robustness tests and are not used for the figures or headline efficiencies.

Dark fixed-growth yield falls from 4.36 to 4.12 g CDW $\mathrm{mol^{-1}}$ between $\mu = 0.126$ and 0.063 $\mathrm{h^{-1}}$. Holding the reference antenna–retinal design fixed would reduce $\eta$ to 2.19% at the lower rate; re-optimisation recovers 3.47% by moving from 6.57 g $\mathrm{L^{-1}}$ and $V/A = 0.0207$ m to 2.74 g $\mathrm{L^{-1}}$ and 0.0970 m. The selected retinal maximum shifts only from 552 to 554 nm, and the concentration limit remains active.

The thermodynamic-limit carotenoid branch gives 10.70%, 10.22% and 10.10% at the same three rates. Every point remains on the 600-nm and 50-PPFD bounds, so these coordinates do not identify an interior spectral optimum. Reproduce the calculation with `PaperScripts/reactor_geometry/data/eta_net_vs_growth_setpoint.py`; output is in `Results/reactor_geometry_wavelength/eta_net_vs_growth_setpoint.csv`.

##### S6.8 Sensitivity to the pump cross-section calibration ($\sigma_{\mathrm{eff}}$)

`SIGMA_PUMP_SCALE` (= 4.5, `Bin/PhysBin/ReactorOptics/reactor_model.py`) is an inherited lumped scale whose independent calibration datum is undocumented (Methods), giving $\sigma_{\mathrm{eff}}$ =

$\sigma_0 \times$ SIGMA_PUMP_SCALE. It enters the *forward* map (illumination to $H^+$ flux) and the *inverse* map (Phase-II LP table index). Rescaling both largely compensates by construction (`Results/reactor_geometry_wavelength/sigma_mirrored_test.csv`); the physical sensitivity test rescales only the forward map (`PaperScripts/reactor_geometry/data/audit_sigma_eff_scale.py`).

For pump-rate ratios, a common cross-section multiplier cancels to leading order only when both rates are in the low-activation regime, where $R_{\text{pump}} \approx k_{\text{abs}}\Phi_{\text{QY}}$. It does not generally cancel inside $1 - \exp(-k_{\text{abs}}\Phi_{\text{QY}}\tau)$, or from the constrained metabolic response. Pathway and wavelength comparisons therefore require sensitivity analysis even when they share a calibration. The effect on whole-collector efficiency is quantified with the six-variable optimiser behind Table S11, at 45% occupancy, both scenarios, $CO_2$ capture charged (`PaperScripts/reactor_geometry/data/audit_sigma_eff_scale_headline.py`; output `Results/reactor_geometry_wavelength/sigma_eff_scale_headline_dac.csv`).

Table S5: **Point-estimate sensitivity of the closed-boundary, capture-charged optimum to the pump cross-section calibration.** (Related to Fig. 7.) Six-variable rGlyP, cyl. external, 45% occupancy and $CO_2$ capture charged, otherwise identical to Table S11's "Closed boundary" row. $\sigma_{\text{eff}}/\sigma_0 = 4.5$ is nominal; only the forward pump-rate map is rescaled.

| Scenario | 0.5× ($\sigma_{\text{eff}}/\sigma_0 = 2.25$) | 1× (nominal, 4.5) | 2× (9.0) | Swing (0.5× → 2×) |
|---|---|---|---|---|
| SOTA | 2.69% | 3.30% | 4.63% | −0.61 / +1.33 pp (−18.5% / +40.4% rel.) |
| Thermodynamic limit | 10.87% | 10.89% | 11.52% | −0.01 / +0.63 pp (−0.1% / +5.8% rel.) |

At SOTA $t = 1$ throughout, so $\eta$ responds near-linearly (−18.5% at 0.5×, +40.4% at 2×, comparable to the ∼1.9× formate-saving sensitivity). At the thermodynamic limit $t \approx 0.03$ (Table S27) at 0.5× and nominal scale, so $\eta$ is flat (10.87 → 10.89%); at 2× the optimum flips to $t = 1$, explaining the disproportionate jump.

Over the tested calibration range, the steady design-point sensitivity spans 2.69–4.63% at the state of the art and 10.87–11.52% at the thermodynamic limit.

A 96-draw sensitivity calculation per scenario (`PaperScripts/reactor_geometry/data/propagate_closed_boundary.py--sigma-eff-mult`) multiplies each draw's `pump_rate_scale` by the same factor (forward map only) and uses the same corrected global objective as the headline propagation. The nominal SOTA median is 3.27%, close to 3.25% in the 512-draw analysis.

Table S6: **Coarse ($n = 96$/scenario) Monte Carlo sensitivity of propagated closed-boundary interval to $\sigma_{\text{eff}}$.** (Related to Fig. 7.) Same accounting as Table S5; each draw's `pump_rate_scale` multiplied by the column factor.

| Scenario | Scale | Median $\eta$ (%) | 95% interval (%) |
|---|---|---|---|
| SOTA | 0.5× | 2.62 | 2.23–3.07 |
| SOTA | 1× (nominal) | 3.27 | 2.76–3.84 |
| SOTA | 2× | 4.68 | 3.92–5.39 |
| Thermodynamic limit | 0.5× | 10.47 | 9.01–12.29 |
| Thermodynamic limit | 1× (nominal) | 10.48 | 9.02–12.30 |
| Thermodynamic limit | 2× | 10.80 | 9.44–12.49 |

Pump calibration strongly affects the state-of-the-art distribution but has little effect at the thermodynamic limit until the design changes its dichroic routing. No crop-exceedance fractions are reported because these are steady-design calculations rather than seasonal field predictions.

##### S6.9 Propagated model-boundary uncertainty intervals

Quantified model-boundary uncertainty was propagated through the constrained rGlyP design for every Monte Carlo draw. The interval that results is a statement about the model boundary alone, not a process, TEA or land-use interval; two classes are separated: probabilised terms below, and implementation risks (strain construction, chromophore loading, deployment economics; Table S23).

Each draw sampled PV efficiency, formate electrical demand, biomass heating value, $O_2$-transfer and feed limits, solar-band delivery and concentration, an effective pump/absorption multiplier (quantum-yield posterior plus absorption-calibration uncertainty), and a formate-demand multiplier (GAM plus metabolic-structure term). Density, surface PPFD, depth and dichroic transmission were globally optimised for every draw on the optimum manifold identified by the full six-variable search: external cylinder, allocation at the 45% cap, and wavelength 550 nm (state of the art) or 600 nm (thermodynamic limit).

The earlier random-restart search was discarded because infeasible starts could remain on a flat rejection objective and bias maxima downward. The replacement uses differential evolution for each draw. For the headline whole-collector audit, twelve draws spanning each scenario's calculated distribution were re-optimised over all six variables and every geometry. The largest absolute difference was $4.8 \times 10^{-5}$ percentage points (Table S24). This establishes numerical agreement at the reported precision, but does not address uncertainty in model structure or omitted deployment terms.

At 45% occupancy on the declared route boundary with $CO_2$ supplied, the earlier intervals are 4.00–7.20% for SOTA (median 5.48%) and 9.93–13.40% at the thermodynamic limit (median 11.50%). At the full-budget ceiling they are 5.75–8.38% (median 6.86%) and 9.52–13.07% (median 11.07%), at an occupancy no phototroph reaches. These route-boundary diagnostics credit energy outside the rhodopsin band and should not be substituted for the headline whole-collector interval.

On the closed boundary with $CO_2$ supplied rather than captured, the earlier sensitivity intervals are 3.21–5.76% and 6.98–12.14%; they use a different carbon-supply basis and are not the headline result. Charging capture, the headline basis, gives retinal-only intervals of 2.74–3.89% (median 3.25%) and 9.09–12.24% (median 10.53%). All runs used 512 feasible draws per scenario with no failures. Crop values are not used as thresholds because their seasonal time and land bases differ. The capture-charged results are stored in `Results/reactor_geometry_wavelength/closed_boundary_uncertainty_cap45_dac_global_hhv_t.csv`, `closed_boundary_uncertainty_draws_cap45_dac_global_hhv_t.csv`, and `closed_boundary_global_audit_cap45_dac_global_hhv_t.csv`.

The joint antenna propagation adds uniform evidence ranges of 0.50–0.70 for Kin4B8 transfer, 0.75–1.0 for optical delivery and 0.8–1.2 for the borrowed antenna bandwidths. Re-optimising all 512 draws gives 2.79–4.04% (median 3.35%) for SOTA and 8.93–12.02% (median 10.35%) at the thermodynamic limit. Draw-matched subtraction of the retinal-only result gives a median antenna effect of +0.107 percentage points (95% range −0.034 to +0.284; 93.2% positive) for SOTA and −0.178 points (−0.216 to −0.140;

none positive) at the thermodynamic limit. These are propagated ranges under stated epistemic priors, not empirical confidence intervals. Peak centres remain measured Kin4B8 values. Assembly burden is retained as the separate 6.7% break-even bound because no probability distribution is supported by data. Generated outputs are `carotenoid_uncertainty_joint_antenna_cap45_dac.csv` and its per-draw companion.

##### S6.10 Outside-boundary transport and mixing stress tests

The intervals above exclude balance-of-plant loads. On the closed boundary such loads are met by photovoltaics on the same aperture, so the aperture grows and $\eta$ falls by that enlargement. For scale, a fixed-design auxiliary demand of ∼3.2 W $L^{-1}$ lowers the 45%-occupancy reference from 3.29% to 3.0%, and ∼13.5 W $L^{-1}$ lowers it to 2.2%; per unit reactor surface these are ∼76 and ∼323 W $m^{-2}$. These levels are engineering sensitivity markers, not crop-equivalence or land-productivity thresholds.

The reference cylinder (23.8 mm, 5.4 g $L^{-1}$, 0.68 g $L^{-1}$ $h^{-1}$) has biomass power ∼4.3 W $L^{-1}$ against counted solar input ∼126 W $L^{-1}$. The erasure threshold is only ∼2.5% of input yet costs 11% of $\eta$ because an electrical duty through 24%-efficient PV buys roughly four times its aperture.

Published mixing loads for low-head designs are well below this. Garcia-Galan et al. reported 0.088 W $L^{-1}$ ($\eta' \approx 3.293\%$) for an 8.5 $m^3$ tubular photobioreactor[59]; a twofold allowance (∼0.18 W $L^{-1}$, $\eta' \approx 3.286\%$) remains ∼18× below the erasure threshold.

Airlift power ($P/V \approx \rho g U_g$) implies ∼0.01–0.2 W $L^{-1}$[52]. Pumped tubular reactors cost ∼0.17–1.7 W $L^{-1}$; the upper end exceeds half the erasure threshold. This ∼0.01–0.2 W $L^{-1}$ bracket is below the 0.35 W $L^{-1}$ stress case of Table S11 ($\eta = 3.26\%$ against 3.29%).

These are thresholds, not claims that loads are absent; a TEA/LCA model should add $CO_2$/$NH_4^+$ supply, mixing, pumping and heat rejection. Table S20 confirms: low-head loads preserve the closed-boundary comparison against present field conversion; PV–LED photon routing drops it below.

##### S6.11 Heat removal is not a hidden fourth binding constraint

Heat removal sits outside the accounting boundary; a balance at the optimum shows it is not binding.

Metabolic heat follows the oxycaloric equivalent (≈0.46 kJ per mmol $O_2$). At the closed-boundary state-of-the-art rGlyP optimum (45% cap), volumetric $O_2$ uptake is ∼5.5 mmol $L^{-1}$ $h^{-1}$ (∼0.7 W $L^{-1}$); even at the 100 mmol $L^{-1}$ $h^{-1}$ aeration ceiling it stays below ∼13 W $L^{-1}$. At the 3× optical limit, ∼445 W $m^{-2}$ spread over the closed optimum's 90-mm depth corresponds to ∼5 W $L^{-1}$ of absorbed solar heat.

Both lie within conventional photobioreactor cooling capacity. This confirms the heat load is of a tractable scale, not that the thermal design is solved: a deployment study would still cost the cooling duty (Section S6.10).

##### S6.12 Cell-scale efficiency with absorbed-light accounting

To separate the reactor-optical contribution, we evaluate a diagnostic cell-scale efficiency for one cell at uniform flux $I$, with no spatial gradient or reactor geometry. The numerator is unchanged ($\mu\,\mathrm{HHV} = 0.126 \times 22.4$ kJ gDW$^{-1}$ h$^{-1}$); the denominator counts absorbed light and the solar energy required to supply formate, rather than all sunlight incident on a collection aperture:

$$\eta_{\mathrm{cell}} = \frac{\mu\,\mathrm{HHV}}{\dfrac{\kappa_\rho(\lambda)\,I\,E_{\mathrm{photon}}(\lambda)}{\eta_{\mathrm{dichroic}}} + \dfrac{(\mu/Y)\,\Delta G^{\mathrm{elec}}_{\mathrm{formate}}}{\eta_{\mathrm{PV}}}} \tag{S51}$$

where $\kappa_\rho(\lambda)$ is the mass-specific rhodopsin absorption coefficient (Eq. S36) and $Y$ the fixed-growth yield. Two consistency checks: in the dark limit $\eta_{\mathrm{cell}}$ reproduces the dark reactor value (4.71% rGlyP, 3.99% CBB at SOTA); absorbed flux at the reactor optimum (∼74% of delivered band) keeps the cell-scale denominator as a genuine lower bound.

Table S7: **Cell-scale and reactor-scale upper-bound efficiencies compared.** (Related to Fig. 7.) Cell-scale: maximise Eq. S51 over flux and absorption maximum; reactor-scale: bounded five-variable optima. All at $\mu = 0.126$ h$^{-1}$, full-membrane ceiling (not 45% reference). *Accounting:* route boundary *without* $CO_2$-capture charge, the boundary on which Eq. S51 is written; reactor columns re-optimised on that boundary (not Table S17, which charges capture). The main text's closed, capture-charged boundary scores several points lower (Table S27b); nothing here is comparable with a crop figure.

| Scenario | Pathway | Cell, dark | Cell, optimum | Reactor, ext. cylinder | Reactor, internal bag |
|---|---|---|---|---|---|
| SOTA | rGlyP | 4.71% | 7.97% ($\lambda = 556$ nm) | 7.09% | 5.18% |
| SOTA | CBB | 3.99% | 7.18% ($\lambda = 553$ nm) | 6.28% | 4.16% |
| Thermodynamic limit | rGlyP | 11.94% | 13.14% ($\lambda = 557$ nm) | 11.92% | 11.92% |
| Thermodynamic limit | CBB | 10.13% | 11.19% ($\lambda = 553$ nm) | 10.10% | 10.09% |

The pathway ranking is unchanged: rGlyP leads CBB by 0.79 points at cell scale and 0.81 at the best geometry. The best external geometry costs 0.9 points (7.97 → 7.09%, 11% relative), whereas the internal bag costs 2.8 (7.97 → 5.18%, 35%); the gap closes at the thermodynamic limit (11.92% rGlyP, 10.09–10.10% CBB). The spectral optimum is already set at cell scale (556.0 nm, from $E = hc/\lambda$), and the reactor moves $\lambda^\star$ by under half a nanometre.

At $\mu = 0.063$ h$^{-1}$ the structure holds (cell 560.9 nm, reactor 561.4 nm); the thermodynamic-limit reactor optimum remains on the 600-nm and minimum-PPFD bounds. The cell-scale objective is nearly degenerate across the green region; the supported statement is that photon energy moves the optimum off the blue edge. Optimum flux falls with the setpoint ($I \approx 204$ to $\approx 104$ $\mu$mol m$^{-2}$ s$^{-1}$). Reproduce with `PaperScripts/reactor_geometry/data/cell_scale_decomposition.py`.

Consistency check: the cell-scale optimum ($I \approx 204$ $\mu$mol m$^{-2}$ s$^{-1}$) matches reactor volume-averaged flux (214 $\mu$mol m$^{-2}$ s$^{-1}$) to ∼5%. The optimiser places the average cell at its single-cell optimum, at tenfold higher surface flux ($f_{\mathrm{del}} \approx 0.104$).

#### S7 Wavelength-range selection and how far the data constrain it

Figure 5 uses a 450–600 nm design sweep. The posterior is over physical parameters, so the simulator predicts at any $\lambda$; what varies is how tightly data constrain that prediction. The sweep contains one calibration wavelength (568 nm) and three design points (450, 520, 600 nm). The extended 450–700 nm sweep (Figure S4) shows how credible intervals behave outside that span.

##### S7.1 What the calibration contains, and what it does not

No mutant enters either photophysical likelihood. The quantum-yield block is trained on exactly one observable, the wild-type photoisomerisation yield ($\Phi_{\text{iso}} = 0.64 \pm 0.04$ at 568 nm)[11], with a prior penalising yields above 0.90 (Section S2); the photocycle block (Section S3) is calibrated against two wild-type timing targets ($\tau_M \approx 5$ ms, $\tau_{\text{total}} \approx 15$ ms)[6,16]. Sixteen free parameters are therefore anchored on three wild-type observables at a single wavelength.

##### S7.2 What independent variant data exist, and where they sit

All independent bacteriorhodopsin variant measurements lie at or to the red of 568 nm; none is blue-shifted. Logunov et al. report photoisomerisation yields for wild type (568 nm) and four variants: D212N (570 nm), R82Q (590 nm), deionised blue bR (605 nm) and D85N (610 nm)[14]. Marinetti et al. report proton-translocation yields placing D212N at 580 nm and D85N at 613 nm[60] (Table S12). The photocycle reference set (Section S3) places R82Q, D212N and D85N at +7, +17 and +32 nm from 568 nm. Its D85N value differs by more than 10 nm, and we quote each value from its own source.

None entered a likelihood (Section S9); all are out-of-sample checks, exclusively on wavelengths $\geq 568$ nm.

##### S7.3 How the credible intervals vary across the band

The quantum-yield 95% posterior-predictive CI (2.5th–97.5th percentile; `Bin/PhysBin/QuantumYield/wavelength_raw_qy_samples.csv`), narrowest at 568 nm, spans 0.51–0.74 at 520 nm and 0.44–0.69 at 600 nm. The width at 600 nm (0.25) nearly matches that at 450 nm (0.27): the unconstrained $\beta$ prior (Section S1) compounds with a four-parameter posterior that broadens continuously with distance from the anchor.

Within 520–600 nm $\Delta E_{\text{opt}}$ stays in the Langevin near-linear regime (Section S3), so photocycle predictions carry only MCMC parameter uncertainty.

##### S7.4 What widens the interval at 450 nm

At 450 nm (0.39 eV above the wild-type maximum) two independent factors widen the interval. For quantum yield, $\beta$ (the CI-shift fraction; Section S1) is sampled from $\text{Uniform}(0, 1)$ per draw with zero likelihood at the calibration wavelength, so the 450 nm prediction is prior-dominated. The direction (higher QY from faster non-adiabatic transfer) is physically motivated, but magnitude cannot be tightened without a multi-wavelength yield measurement.

For photocycle lifetime, Langevin saturation at $|x| \gg 1$ (Section S3) means large spectral shifts produce diminishing barrier changes; the electrostatic coupling parameter's 95% credible interval spans a factor of several (Table S2). This is largest for the rate-limiting $M_1 \rightarrow M_2$ transition, so the lifetime posterior broadens substantially below 540 nm and to the red of 568 nm.

##### S7.5 Low-PPFD regime: QY dominates over photocycle time

Despite that lifetime uncertainty, quantum yield dominates pump-rate variation. At 100–400 PPFD the Poisson saturation parameter (Eq. S22) stays in the near-linear regime ($x = 0.031$–$0.051$ over 520–568 nm at 100 PPFD; 0.12–0.20 at 400; `PaperScripts/physics_data/physics_sweep.csv`), where $\tau$ enters only as a second-order modulator with logarithmic sensitivity <2.6% (Section S3). A ±20% uncertainty on $\tau$ at 450 nm moves the pump rate by 0.3% at 100 PPFD and 1.3% at 400 PPFD; QY uncertainty dominates, already propagated into the photophysics results. At 600 nm the longer photocycle pushes $x \approx 0.6$, so saturation reduces the linear-regime estimate by ∼25% and lifetime becomes a non-negligible modulator, carried as a caveat in those comparisons.

##### S7.6 The extended 450–700 nm sweep

The extended sweep (Figure S4) reveals costs, not boundaries. Under the modelled conditions ($pH_{in}$ 7.5, $pH_{out}$ 7.0, $\Delta\Psi = 150$ mV) the minimum electrochemical work is $\Delta\mu_{H^+} = 0.180$ eV, while the usable energy per isomerisation is 0.193 eV at 700 nm and does not fall below $\Delta\mu_{H^+}$ until ∼719 nm (`Bin/PhysBin/Feasibility/`). The feasibility factor is unity across the sweep; the red-side penalty is photocyclic, not thermodynamic.

Prediction width does change: the coefficient of variation of the lifetime posterior rises from 1.16 at 568 nm to 18.4 at 450 nm and 9.5 at 700 nm (`PaperScripts/physics_data/supplementary_full_range/physics_sweep.csv`), and the quantum-yield interval broadens as the unconstrained $\beta$ prior displaces the likelihood. Spectral engineering thus trades predicted gain against a widening interval; how far to move is a design decision, not a model limit.

#### S8 Thermodynamic feasibility: max–min driving force (MDF) analysis and curated pyTFA

Thermodynamic self-consistency was assessed at three levels: a reverse-$k_{cat}$ (Haldane) approximation inside the enzyme-constrained LP for directionality correctness; pathway-level Max-min Driving Force (MDF) analysis for feasibility under stated concentration bounds; and a curated `pytfa.ThermoModel` over the active assimilation, respiratory, PMF and cofactor-balancing subnetworks for concentration-resolved direction coupling. All three are reproduced by `scripts/analyze_thermo_reversibility.py`, `scripts/audit_headline_thermodynamics.py` and `scripts/run_native_pytfa_mdf.py`; the MDF analysis and serine-concentration sensitivity are in Figure S7.

#### S8.1 Reverse-$k_{\mathrm{cat}}$ treatment and the C1/THF exemption

Each reversible reaction is split into halves competing for the shared proteome pool. Noor et al.[30] showed forward-to-reverse ratios broadly track the equilibrium constant through the Haldane relation $k_{\mathrm{cat,fwd}}/k_{\mathrm{cat,rev}} = K_{\mathrm{eq}}(K_m^{rev}/K_m^{fwd})$, with a typical ratio near 10. Lacking genome-scale $K_m$ data for *C. necator*, a uniform $\kappa = 0.1$ is applied to all reversible reactions, imposing a tenfold capacity penalty for reverse operation.

A small set of rGlyP $C_1$/tetrahydrofolate reactions are exempted: formate–THF ligase (Ftl), methenyl-THF cyclohydrolase (Fch), methylene-THF dehydrogenase, the glycine cleavage system and serine hydroxymethyltransferase. They carry assimilatory $C_1$ flux in the biosynthetic direction, which is thermodynamically downhill under the formate-excess, NADPH-rich, low-THF-product conditions modelled. Applying the penalty would impose an artificial tenfold disadvantage on rGlyP without biochemical justification. The exemption is treated as an explicit sensitivity assumption: Section S14 confirms the fixed-growth advantage remains positive at +10.4% even with the penalty applied (against +15.2% unpenalised), so it determines magnitude but not sign.

#### S8.2 MDF formulation and reference bounds

The MDF criterion[61] asks whether metabolite concentrations exist, within specified bounds, making every reaction thermodynamically downhill:

$$\mathrm{MDF} \;=\; \max_{\mathbf{x}} \min_{r\in\mathcal{R}} \left(-\Delta_r G'_r(\mathbf{x})\right), \tag{S52}$$

where $\mathbf{x} = \ln \mathbf{c}$ and $\Delta_r G'_r = \Delta_r G'^{\circ}_r + RT\,\mathbf{S}_r \cdot \mathbf{x}$. A positive MDF indicates feasibility. Standard transformed Gibbs energies came from eQuilibrator at pH 7, $I$ = 0.25 M and $p$Mg 3[62]. Free concentrations were bounded within [1 $\mu$M, 10 mM]; key cofactor ratios (ATP/ADP, NADPH/$NADP^+$, $NAD^+$/NADH) were fixed at eQuilibrator defaults, $CO_2$ and formate at their mass-transfer-implied upper concentrations, and $H_2O$ at unit activity. The optimisation is an exact LP in log-concentration variables.

#### S8.3 MDF results and the rGlyP–CBB gap

Under the reference bounds rGlyP has MDF = +6.8 and CBB +4.4 kJ mol$^{-1}$, so both routes are feasible. The +2.4 kJ mol$^{-1}$ rGlyP advantage arises because the reductive route has fewer reversible steps with small or positive $\Delta_r G'^{\circ}$ than the eleven-step CBB cycle, whose limiting reaction is typically RuBisCO carboxylation or phosphoglycerate kinase.

The gap does *not* persist when the concentration window is widened: raising the upper bound to 0.1 M gives +6.6 kJ mol$^{-1}$ for CBB and +6.8 to +7.5 for rGlyP, i.e. near parity (`Results/Thermodynamics/native_pytfa/mdf_sensitivity.csv`), because CBB's limiting reactions benefit disproportionately from relaxed product bounds. We therefore claim only joint feasibility, not a thermodynamic driving-force advantage for rGlyP.

##### S8.4 PMF sensitivity band on absolute formate demand

Simulations were repeated over $\Delta\Psi = 0.1$–$0.2$ V and $\Delta$pH = 0–1, giving $\Delta\mu_{H^+} = e\Delta\Psi + 2.303\, k_B T\, \Delta\text{pH}$ of roughly 100–250 meV, spanning reported values for actively growing bacteria. Over this range the 100 PPFD rGlyP formate demand spans 20.9–41.1 and CBB 27.0–76.2 mmol $\text{gDW}^{-1}$ $\text{h}^{-1}$ (`Results/Thermodynamics/curated_tfa_pmf_sensitivity.csv`). The rGlyP < CBB ranking holds throughout, but the roughly twofold (rGlyP) and threefold (CBB) spreads show that absolute formate-saving magnitudes carry substantial electrochemical-load uncertainty, unresolvable without organism-specific PMF measurements. The percentage saving varies less because PMF uncertainty shifts both pathways in the same direction.

##### S8.5 Curated pyTFA scope and limitations

A curated `pytfa.ThermoModel` applies log-concentration bounds, reaction-$\Delta_r G'$ constraints and direction coupling to the assimilation block (12 rGlyP and 16 CBB base reactions), the respiratory chain (NADH dehydrogenase, cytochrome $bc_1$, cytochrome oxidase), $F_1F_0$-ATP synthase with explicit PMF coupling, and the key cofactor-balancing reactions (transhydrogenase, thioredoxin reductase), with pyruvate dehydrogenase and aconitase as aggregate reactions. Compartment pH and membrane potential are treated explicitly so that $H^+$ translocation work enters the direction constraints correctly. The curated scope is complete within itself (12/12 and 16/16 base reactions); those fractions refer to that predefined scope, not to all active genome-scale reactions.

No genome-scale formation-energy database exists for *C. necator*; standard Gibbs energies were curated by hand for the reviewed reactions, so the remaining active reactions (TCA cycle, anaplerosis, biosynthesis, cell-wall precursors) are uncovered, and constraint-topology claims remain LP optima under the Haldane approximation. The $\kappa = 0.1$ factor is unrelated to reaction-specific $K_{\text{eq}}$ and $K_m$ ratios; the curated model provides directional consistency checks but does not validate that factor quantitatively (Section S14). The concentration bounds are standard physiological ranges, not *C. necator* formate metabolomics, which would tighten the MDF analysis and narrow the PMF band.

Table S2: Parameters of the twelve-parameter photocycle kinetics model (Marcus/Grote–Hynes framework). All parameters are sampled from uniform priors over the ranges shown; $\theta \in [0,1]$ denotes the unit-hypercube coordinate used in the MCMC prior transform (Section S2). Attempt-frequency prefactors ($k_0$, $k_N$, $k_O$) are fixed at the quantum transition-state-theory value $k_\mathrm{B}T/h \approx 6.25 \times 10^{12}$ s$^{-1}$ and are not free parameters. The three reorganisation energies and driving forces parameterise the Marcus barriers (Eq. S16), the base barriers set the early/late thermal steps (Eq. S19), and $\alpha_L$, $E_\mathrm{pole}$ and $\sigma_\mathrm{conf}$ govern the spectral coupling and its inhomogeneous broadening (Eqs. S17–S18).

| **Symbol** | **Name / description** | **Prior range** | **Units** | **Physical role** |
|---|---|---|---|---|
| $\lambda_0$ | $M_1 \to M_2$ reorganisation energy | [0.50, 2.00] | eV | Sets Marcus barrier height for M-state decay |
| $\Delta G_0$ | $M_1 \to M_2$ driving force | [−0.20, 0.20] | eV | Controls thermodynamic balance of M→N step |
| $\alpha_L$ | Langevin spectral coupling coefficient | [0.05, 0.45] | n/a | Scales how spectral shift modulates the Marcus barrier |
| $\sigma_\mathrm{conf}$ | Conformational disorder amplitude | [0.01, 0.06] | eV | Inhomogeneous broadening of the barrier distribution |
| $\kappa_\mathrm{fric}$ | Grote-Hynes transmission coefficient | [0.10, 0.60] | n/a | Friction correction to transition-state crossing rate |
| $\lambda_N$ | $M_2 \to$N reorganisation energy | [0.50, 1.50] | eV | Marcus barrier height for N-state formation |
| $\Delta G_N$ | $M_2 \to$N driving force | [−0.20, 0.20] | eV | Free-energy difference between M and N intermediates |
| $\Delta G_O$ | N→O driving force | [0.00, 0.40] | eV | Free-energy difference between N and O intermediates |
| $E_\mathrm{pole}$ | Electrostatic pole coupling energy | [0.02, 0.10] | eV | Characteristic energy scale for Langevin saturation of spectral coupling |
| $W_{K,\mathrm{base}}$ | K→L base barrier height | [0.30, 0.50] | eV | Intrinsic thermal barrier for K→L early photocycle step |
| $W_{L,\mathrm{base}}$ | L→$M_1$ base barrier height | [0.40, 0.60] | eV | Intrinsic thermal barrier for L→M proton-transfer step |
| $W_{O,\mathrm{base}}$ | O→bR base barrier height | [0.50, 0.70] | eV | Intrinsic thermal barrier for final ground-state recovery |

Table S3: Symbols used in the enzyme-constrained metabolic model. Values are those used throughout this work; the full principal-parameter inventory is given in Table S14.

| **Symbol** | **Meaning** | **Value / units** |
|---|---|---|
| $\mathbf{S}$ | Stoichiometric matrix | $1348 \times 1538$ |
| $\mathbf{v}$ | Flux vector | $\mathrm{mmol\,gDW^{-1}\,h^{-1}}$ |
| $v_{\mathrm{bio}}$ | Biomass-synthesis flux (growth rate) | $\mathrm{h^{-1}}$ |
| GAM | Growth-associated maintenance | 135.0 mmol ATP $\mathrm{gDW^{-1}}$ |
| NGAM | Non-growth-associated maintenance | 3.0 mmol ATP $\mathrm{gDW^{-1}}$ $\mathrm{h^{-1}}$ |
| $k_{\mathrm{cat},j}$ | Turnover number of reaction $j$ | $\mathrm{s^{-1}}$ (tabulated) |
| $\sigma_j$ | Apparent saturation factor | $(0, 1]$ (EC-class assigned) |
| $\mathrm{MW}_j$ | Enzyme molecular weight | kDa $\equiv$ $\mathrm{g\,mmol^{-1}}$ |
| $c_j$ | Proteome draw coefficient | g per $\mathrm{mmol\,gDW^{-1}\,h^{-1}}$ |
| $\kappa$ | Haldane reverse-turnover factor | 0.1 |
| $P_{\mathrm{total}}$ | Total protein content | 0.68 $\mathrm{g\,gDW^{-1}}$ |
| $f_{\mathrm{enzyme}}$ | Enzyme+ribosome protein fraction | 0.50 |
| $f_{\mathrm{ribo}}(\mu)$ | Ribosome protein fraction | $0.068 + 0.157\,\mu$ |
| $f_{\mathrm{ribo,max}}$ | Ribosome fraction ceiling | 0.40 |
| $f_{\mathrm{enzyme,eff}}$ | Effective metabolic-enzyme fraction | 0.4122 at $\mu = 0.126$ |
| $a_j$ | Membrane-area draw per unit flux | $\mu\mathrm{m}^2$ per $\mathrm{mmol\,gDW^{-1}\,h^{-1}}$ |
| $A_{\mathrm{cell}}$ | Inner-membrane area budget | 20 $\mathrm{m^2\,gDW^{-1}}$ |
| $R_{\mathrm{pump}}$ | Rhodopsin per-molecule pump rate (= $k_{\mathrm{cat,rho}}$) | $\approx$ 3.85 $\mathrm{H^+\,s^{-1}}$ (WT, 100 PPFD) |
| $d$ | Fixed-point damping factor | 0.5 (adaptive) |

### Supplemental Results

#### S9 Photophysics model validation against published experimental data

The quantum-yield and photocycle models were calibrated to wild-type bacteriorhodopsin observables and compared with three published datasets outside the likelihood (Table S12, Figure S14). These are qualitative checks, not an integrated validation: the studies use different temperatures, variants and observables, while the quantum-yield posterior is anchored by one value ($\Phi_{\text{iso}} = 0.64 \pm 0.04$ at 568 nm[11]).

##### Excitation-wavelength dependence at 110 K (Xie 1990)

Xie measured the absolute forward photoreaction QY ($\varphi_1$, bR→K) for wild-type bR at 110 K from 420 to 600 nm[63]. At cryogenic temperature the photocycle stalls after K, so $\varphi_1$ reports isomerisation alone. It declines from ≈0.61 at 420 nm to ≈0.54 at 600 nm. The comparison tests only whether the model reproduces that direction across wavelength.

The decline is qualitatively reproduced: the posterior median runs 0.71 at 420 nm to 0.58 at 600 nm (Figure S14a), the same direction as Xie's ≈0.61 to ≈0.54 but falling about twice as far and sitting above the cryogenic measurement at both ends. The 420 nm value comes from `PaperScripts/physics_data/generate_qy_420nm_point.py`, the 600 nm median from `Bin/PhysBin/QuantumYield/wavelength_raw_qy_samples.csv`. The comparison is indirect: the model is parameterised for room temperature, so only the trend is expected to match at 110 K. The WT calibration target of $0.64 \pm 0.04$[11] and the alternative room-temperature value of $0.58 \pm 0.05$[14] bracket the Xie values. The posterior median of ≈0.60 at 568 nm undershoots the target by ∼6% through Bayesian shrinkage, and the 95% credible interval (0.50–0.70) encompasses the Xie values (0.54–0.61) and both WT determinations (0.58 and 0.64).

##### Mutant absorption maxima at room temperature (Logunov 1996)

Logunov et al. measured $\Phi_{\text{iso}}$ for wild-type bR and four spectral variants by femtosecond transient absorption and photoacoustic spectroscopy[14]. Values span WT $0.58 \pm 0.05$ at 568 nm to D85N $0.75 \pm 0.15$ at 610 nm. The point estimates rise modestly with red shift, opposite to the model median, although the mutant uncertainties overlap.

The model median declines from ≈0.64 at 520 nm to ≈0.58 at 600 nm (`Bin/PhysBin/QuantumYield/wavelength_raw_qy_samples.csv`; Figure S14b). Its one-dimensional coordinate cannot represent residue-specific changes caused by D85, D212 or R82 substitution. The comparison therefore exposes a disagreement in mutant ordering rather than validating the predicted trend. The two WT measurements also differ (0.64[11] against $0.58 \pm 0.05$[14]).

##### Proton translocation QY and photocycle kinetics (Marinetti 1989)

Marinetti et al. measured proton translocation QY ($\varphi_t$) and proton release/uptake kinetics for bR variants in *E. coli*[60]. Unlike $\Phi_{\mathrm{iso}}$, $\varphi_t$ counts charge translocation events per absorbed photon, so a mutant can carry high $\Phi_{\mathrm{iso}}$ with low $\varphi_t$ if downstream proton machinery is disrupted. D85N has $\varphi_t = 0.03$ at pH 6 (Figure S14c), eliminating vectorial translocation despite $\Phi_{\mathrm{iso}} = 0.75 \pm 0.15$; this confirms Asp85 as the primary extracellular acceptor during the M transition, uninvolved in the chromophore torsional coordinate. The model couples pump flux through $\varphi_t \approx \Phi_{\mathrm{iso}}$ (near-unity coupling under physiological conditions[6]), appropriate for intact WT-like variants; acceptor-deficient mutants would need an explicit decoupling term and are outside the validated design space.

D96N has $\varphi_t = 0.78$, close to WT, but its cytoplasmic uptake half-time extends to 500–1000 ms (Figure S14d) against the WT ∼5 ms, because Asp96 is the internal donor and its absence forces far slower reprotonation by diffusion. The photocycle model, calibrated to WT targets ($\tau_M \approx 5$ ms, $\tau_{\mathrm{total}} \approx 15$ ms)[6,16], keeps D212N on the barrier-driven trend at ∼70 ms for a +20 nm shift (`PaperScripts/physics_data/physics_sweep.csv`), a few-fold above WT rather than two orders above it. It cannot reproduce the D96N anomaly (750 ms at $\Delta\lambda \approx 0$), because that timescale arises from losing a specific proton-shuttling residue rather than from a barrier perturbation.

##### Assessment

The Xie comparison supports the sign of the wild-type wavelength trend but not its magnitude. The Logunov variants disagree in ordering, and the Marinetti mutants show failure modes that the reduced coordinate cannot represent. No dataset tests the complete sequence from wavelength through pump abundance and formate saving. The 95% QY band in Figure S14 is propagated downstream, but it does not cover this structural mismatch, the unresolved pump scale or antenna assembly.

#### S10 Shadow prices across enzyme-capacity policies

Shadow prices are the LP duals on the metabolite balances $\mathbf{Sv} = \mathbf{0}$. Under fixed-growth formate minimisation a negative price on an energy cofactor means that balance is binding: one additional mmol gDW$^{-1}$ h$^{-1}$ of that cofactor would sustain the same growth on fewer mmol of formate. Prices on non-binding balances are zero by complementary slackness, so the active set, comprising constraints with non-zero duals, encodes which capacity limits bind simultaneously at the optimum and is the mechanistic readout of Fig. 4.

All analyses pin $v_{\mathrm{bio}}$ at 0.126 h$^{-1}$ and minimise formate while sweeping per-cell PPFD, isolating substrate efficiency at equal productivity. The rGlyP advantage appears as a lower dark minimum-formate floor, a steeper saving slope as light displaces respiratory ATP, and an earlier crossing of the minimum-feasible-formate boundary.

#### Invariance of the active set across saturation policies

Four policies were evaluated (Section S4.8): unconstrained; uniformly saturated; uniform 50% occupancy; and EC-class-specific multipliers from 0.35 for $O_2$-dependent oxidoreductases to 0.75 for NAD(P)H-linked ones. The last is primary throughout, for the reasons given in Section S4.

Across all four the rGlyP advantage is preserved and the active set is qualitatively identical: NADPH and NADH are the most limiting cofactors in dark and low light, ATP prices are smaller but non-negligible, and the relative ranking holds. The active set is fixed by the stoichiometric structure of the two pathways and the biomass equation, not by the magnitude of the capacity bounds: changing a policy rescales the effective pool without altering which reactions are necessary to close the carbon and energy balances.

What does change is the absolute magnitude of prices and the PPFD at which capacity constraints bind (unconstrained $\rightarrow$ smaller prices; fully saturated $\rightarrow$ earlier binding; EC-specific in between). The formate-demand ranking is invariant: rGlyP starts lower and saturates earlier, while the marginal formate displaced per unit pump-derived ATP is approximately equal for both pathways. In both, pump-derived ATP substitutes for respiratory formate oxidation at nearly the same rate.

#### Why rGlyP saves relatively more

The per-ATP saving rate is approximately equal, but rGlyP starts with lower dark formate demand because it uses formate carbon and reductant more efficiently. Pump-derived ATP therefore meets its residual respiratory requirement at a lower photon dose, and rGlyP reaches the minimum-feasible-formate threshold before CBB. In Fig. 4 the rGlyP curve is shifted down and to the left; the associated proteome saturation ceiling and photon-utilisation diagnostic are in Figure S2c.

Free-growth duals give a complementary diagnostic (proximal growth-rate bottleneck rather than substrate-saving value). The free-growth ATP price ($\approx -0.17$ mmol gDW per mmol formate equivalent) is more negative than the fixed-growth value ($\approx -0.14$), confirming ATP supply is more limiting when growth is unconstrained. The free-growth active set additionally includes the proteome-pool and membrane-area constraints. NADPH and ATP bind simultaneously, with rGlyP less constrained than CBB at equal formate, under both objectives. The fixed-growth topology therefore does not arise from the objective alone.

#### Respiratory $O_2$ decline

As pump-derived ATP displaces respiratory ATP with rising PPFD, formate oxidation through FDH falls, directly reducing electron flow to the terminal oxidases and hence $O_2$ consumption (Figure S3a). The decline is approximately proportional to the formate saving up to full respiratory substitution, so the $O_2$ curve tracks the saving curve with a coefficient set by the respiratory quotient of formate oxidation. This matters for reactor design: aeration demand falls at higher PPFD, partially relieving the gas–liquid $O_2$ transfer ceiling, so solar-light delivery sets the optimal operating point instead of aeration. The profiles are monotone functions of PPFD across all four policies with the same shape and rGlyP$<$CBB ordering, confirming insensitivity to enzyme-capacity parameterisation.

#### S11 Proteome allocation validation against published *C. necator* data

Figure S2b compares model-predicted dark (PPFD = 0, $\mu = 0.126\ \text{h}^{-1}$) GECKO enzyme-mass fractions against quantitative proteomics for *C. necator* H16 and, where no native data exist, the closest heterologous analogue[23,64,65]. The basis is g enzyme $\text{gDW}^{-1}$ using the Jahn-calibrated total protein of 0.68 g $\text{gDW}^{-1}$ (Figure S6). Plotted Jahn points use the nearest measured formate growth rate ($\mu = 0.15\ \text{h}^{-1}$); interpolating to the setpoint gives the same conclusion (ribosome 0.0607, CBB 0.0274, FDH 0.0151 g $\text{gDW}^{-1}$). This is a sector-scale plausibility check, not a full quantitative proteome fit.

##### Calibration basis

The 0.68 g $\text{gDW}^{-1}$ total-protein fraction comes from Jahn et al. proteomics of formate-grown H16[23] and sets the mass denominator for every sector comparison. Calibration across five growth rates by ordinary least squares (Figure S6) gives the ribosome regression $f_{\text{ribo}} = 0.068 + 0.157\,\mu$ (capped at 0.40, $r^2 = 0.95$) and a metabolic-enzyme pool bound of $\approx$0.280 g $\text{gDW}^{-1}$ at $\mu = 0.126\ \text{h}^{-1}$. These quantities are anchored by whole-proteome regression before any sector-level prediction is evaluated, so they are not free parameters fitted per sector.

##### Calvin–Benson–Bassham cycle

The CBB sector has the strongest experimental anchor: Schwartz et al. found the RuBisCO CbbLS complex at 3–7% of soluble protein and the full CbbLSXFPTKBA operon at ∼5–10% of total cell protein[66]; Kohlmann et al. confirmed CbbL dominance by iTRAQ[65], and Bowien & Kusian review CbbR-regulated induction[67]. The model predicts 0.0179 g $\text{gDW}^{-1}$ for the CBB strain against 0.0301 at the nearest Jahn point. As ∼2.6% of total protein the prediction sits just below the Schwartz CbbLS range, supporting the order of magnitude without validating the exact allocation.

In the rGlyP strain CBB enzymes are not expressed, and the model predicts Alloc_CBB $\approx 3 \times 10^{-4}$ g $\text{gDW}^{-1}$, confirming no spurious CBB mass is allocated through incomplete GPR exclusion or residual isozyme routes.

##### Formate dehydrogenase and central metabolism

Cramm reviewed the H16 formate dehydrogenases (FDH-S, FDH-O, FDH-I)[64], and quantitative surveys place total FDH at ∼2–4% of cell protein under formate growth[65]; the Jahn formate sector gives 0.0157 g $\text{gDW}^{-1}$. Jugder et al. report 3–12× FDH-I and 10–50× CBB operon induction on formate relative to fructose[68], supporting the partition direction but not comparable on a mass basis for multi-subunit complexes.

The model charges FDH as the NAD-dependent FdsABGD complex via a reaction-level complex mass and turnover override. With that correction the CBB-strain allocation is 0.0180 g $\text{gDW}^{-1}$ against the Jahn point at 0.0157 and the interpolated setpoint at 0.0151; the rGlyP-strain value is lower (0.0133) as expected from its smaller respiratory oxidation flux. The corrected accounting supports the FDH cost scale; residual uncertainty remains in isoform composition and in vivo saturation. (A degenerate-objective bug previously let this pair vary between reruns; see `PaperScripts/Supplementary/plots/plot_supp_proteome_validation.py` for the fix.)

#### Reductive glycine pathway and heterologous photosystem

rGlyP has been engineered in *C. necator* and supplies the organism-level yield anchor[24,25], but no absolute proteome fraction for the heterologous enzyme block at this operating point has been reported. The closest reference is heterologous rGlyP in *E. coli*, where the glycine cleavage system, SHMT and formate–THF ligase block reaches ∼3–8% of cell protein from medium-strength promoters[69,70]. The model predicts 0.0169 g $gDW^{-1}$ (≈2.5% of total protein), just below that band. This is an indirect comparison: it assumes the *E. coli* burden transfers and that those promoter strengths carry the modelled flux, neither of which is guaranteed given the different cell volume and translational machinery. Directional, same-organism evidence is available even without an absolute figure: Dronsella et al. report that moving the pathway from a plasmid-based construct (CRG4) to genome integration (CRG6, the strain behind the yield anchor above) freed ∼13% of the rGlyP proteome allocation, with the C3 module falling more than tenfold[24]. This confirms rGlyP carries a non-trivial, construct-dependent proteome cost in *C. necator* itself, consistent with though not numerically matched to the *E. coli*-derived band. Direct *C. necator* rGlyP proteomics at this operating point remains the highest-priority follow-up.

Heterologous bacteriorhodopsin expression in *C. necator* has not been quantified. As an upper bound, native *Halobacterium salinarum* purple membrane occupies ∼30–50% of cytoplasmic-membrane area and ∼5–12% of cell protein when fully induced[71]; this is an engineering ceiling, not a target. Outside rhodopsin biology, mitochondrial inner membranes independently reach 40–50% areal protein density against $\sim 24 \pm 6\%$ for native *E. coli*[72], so the 30–45% reference band is within a range biological membranes generally tolerate, even though it remains unconfirmed for rhodopsin specifically in this host. The model's dark photosystem allocation falls below this band because useful pump flux is zero at zero PPFD. The pump is a GECKO-costed reaction drawing on the shared protein and membrane pools; the reactor-optics reference abundance $n_{\text{rho}} = 5 \times 10^{-6}$ mol $gDW^{-1}$ is not imposed as an expression target on the dark optimisation. The model captures the direct protein and membrane opportunity cost of productive pumping but not retinal loading, folding-stress proteostasis or other heterologous-expression overheads, leaving the photosystem-sector burden as the largest validation gap.

#### Verdict and required follow-up

Ribosome mass agrees closely with the Jahn formate proteome, but this is a fit diagnostic ($f_{\text{ribo}}$ was regressed against that data), not a prediction. The CBB sector is the right order of magnitude but below the nearest Jahn point; FDH matches after the complex-level correction. Neither discrepancy overturns the pathway ranking, which is set primarily by stoichiometry with both pathways drawing on the same calibrated pool, but they identify CBB-sector enzyme costing and heterologous-sector measurement as priorities for reaction-level proteomic calibration.

The two weakest anchors, rGlyP block allocation and photosystem burden, carry cross-organism transfer uncertainty. The qualitative ranking and constraint-topology sign are insensitive to absolute sector magnitudes so long as both pathways share the pool constraint and the dominant formate-metabolism sector is correctly anchored.

#### S12 Dark formatotrophic baseline benchmark against published yields

AI-designed rhodopsins expressed in *C. necator* increase growth rate and cellular ATP fraction under illumination[73], but formate demand at fixed growth, the quantity predicted here, was not measured. We therefore compare dark biomass-on-formate yields and the rGlyP-over-CBB ordering with published measurements (Figure S10a). The model uses literature maintenance values, an aggregate proteome calibration to formate-grown wild-type *C. necator*, and assigned enzyme-saturation factors. Maintenance and saturation were not refitted to the yield measurements in this analysis. These comparisons assess the model conditional on those inputs; the shared physiological basis limits their independence.

##### Biomass yield on formate

Yield is an emergent LP output (formate minimised at fixed growth), not a fitted quantity. At the setpoint ($\mu = 0.126$ h$^{-1}$) dark formate demands are 28.887 and 34.060 mmol gDW$^{-1}$ h$^{-1}$ for rGlyP and CBB, giving yields of 4.362 and 3.699 g CDW mol$^{-1}$ formate; on a common basis (g CDW per g formic acid; 46.03 g mol$^{-1}$, biomass 1 Cmol = 24.6 g) that is 0.095 and 0.080 g g$^{-1}$. Dronsella et al. measured chemostat yields of $4.52 \pm 0.17$ and $3.88 \pm 0.19$ g CDW mol$^{-1}$ formate (0.098 and 0.084 g g$^{-1}$)[24]. The model sits 3.5% below for rGlyP and 4.7% below for CBB. These deviations are 0.93 and 0.95 standard deviations, within measurement error.

Grunwald et al. reported a formate-limited continuous-culture wild-type yield of 0.16 Cmol Cmol$^{-1}$ (0.086 g g$^{-1}$, 3.94 g CDW mol$^{-1}$)[74]. GAM is the literature value of Jahn et al.[23], adopted unchanged as Dronsella et al. did on this reconstruction[24]; it was not refitted for this comparison. Their culture ran at $D = 0.05$ h$^{-1}$; the model gives 0.1397 Cmol Cmol$^{-1}$ at $\mu = 0.05$ h$^{-1}$ (12.7% below measurement) and 0.1504 Cmol Cmol$^{-1}$ at 0.126 h$^{-1}$ (6.0% below). The first comparison matches the measured dilution rate; the second reports the paper's design setpoint.

##### Relative pathway yield

Both pathways share the proteome pool and biomass composition, which reduces some sources of variation in the comparison but does not guarantee cancellation of parameter errors. The model predicts +17.9% rGlyP yield advantage (15.2% formate-demand reduction); Dronsella et al. measured +16.5% (4.52 vs. 3.88 g CDW mol$^{-1}$, 14.2% formate-demand reduction), reported as "17% higher than that of the wild type"[24]. The predicted advantage was not fitted to this ratio and keeps its sign across all four enzyme-cost scenarios, ranging 10.4–20.3% (Section S14). This supports the predicted dark pathway ordering within the tested assumptions.

##### The residual gap, and why GAM was not refitted

The model under-predicts yield against both datasets and at both growth rates. We retain this residual because the data do not isolate its source. The respiratory-budget correction (Methods) improved every comparison: under the earlier formulation, where Complex I and two further proton-pumping reactions were not credited to the ATP-synthase proton budget and respiration was capped at 6 $H^+$ per formate, yields were 3.951 and 3.322 g CDW $mol^{-1}$ (12.6% and 14.4% below Dronsella et al.), with a predicted advantage of +18.9%. After correction, the predicted advantage moved towards the measured +16.5%.

Other assumptions affect yield in opposing directions. Ideal proton stoichiometry with no leak or slip can over-predict respiratory efficiency, whereas enzyme and maintenance costs reduce yield, so a small net residual neither identifies one cause nor excludes compensating errors. The gap is also largest where maintenance dominates. Model yield rises from 0.1397 to 0.1504 Cmol C per Cmol substrate between $\mu = 0.05$ and $\mu = 0.126$ $h^{-1}$, the expected Pirt behaviour as the fixed NGAM drain is diluted by faster growth (Section S4). Grunwald et al. measured at the low rate, where that share and the disagreement are largest.

GAM was not refitted because the dataset cannot identify it. Grunwald et al. ran a single dilution rate with no Pirt analysis, so only the GAM+NGAM combination at $D = 0.05$ $h^{-1}$ is constrained. Fitting GAM to that point (NGAM held at 3 mmol ATP $gDW^{-1}$ $h^{-1}$) returns GAM = 70.4 mmol ATP $gDW^{-1}$, about half the literature value. It lies as far below 135 as the ≈260 mmol ATP $gDW^{-1}$ demanded by a 9 $H^+$ chain lies above it. The literature value sits between those fits and is the value Dronsella et al. used on this reconstruction[24], so it is kept unchanged.

##### Growth rate and light coupling

The setpoint ($\mu = 0.126$ $h^{-1}$) and dark reference (0.18 $h^{-1}$) follow the $\mu_{max} = 0.18$ $h^{-1}$ measured for fed-batch wild-type cultures[74]. The enzyme-constrained maximum dark growth rate lands near this cap (consistency check, not a prediction). The CBB proteome-limited dark optimum is $\mu \approx 0.21$ $h^{-1}$ (17% above measured 0.18) and rGlyP reaches $\approx 0.27$ $h^{-1}$; these endpoints are enzyme-cost sensitive and no headline claim rests on them. This deviation runs *opposite* to the yields (which sit below measurement), so the two are not one systematic bias. Experimentally the engineered rGlyP strain grows much more slowly than wild type (batch doubling ∼11 h, ≈0.063 $h^{-1}$, against ∼6 h[24]) owing to heterologous-expression kinetics, which is orthogonal to the stoichiometric yield advantage examined here.

#### S13 RuBisCO engineering does not rescue CBB formatotrophy

CBB carries its carbon through RuBisCO, whose slow turnover and oxygenase side reaction are the classical engineering targets, so it is natural to ask whether a faster or more $CO_2$-specific enzyme would close the gap to rGlyP. Testing both axes showed that neither helps because, in formatotrophy, the RuBisCO burden is not the enzyme but the cycle it drives.

#### Turnover number is non-binding

Sweeping the carboxylase turnover from the wild-type Form II value ($k_{\text{cat}} = 3.0\ \text{s}^{-1}$) up to 50 $\text{s}^{-1}$ spans and exceeds the fastest RuBisCO on record, the *Gallionellaceae* GWS1B enzyme at 25.8 $\text{s}^{-1}$[75]. Dark fixed-growth formate demand remained *identical* to thirteen significant figures (34.06 mmol $\text{gDW}^{-1}\ \text{h}^{-1}$) across the whole range, with the proteome-limited maximum dark growth rate unchanged at $\mu \approx 0.21\ \text{h}^{-1}$.

RuBisCO is far from its catalytic ceiling here because FDH oxidises essentially all incoming formate to $CO_2$ for energy (∼34 mmol $\text{gDW}^{-1}\ \text{h}^{-1}$), of which only the biomass-carbon fraction (∼5.2) is re-fixed. Carrying that modest flux needs ∼0.009 g $\text{gDW}^{-1}$ of RuBisCO. This constraint does not bind, so a faster enzyme frees protein for which nothing competes. The cost that *does* scale is stoichiometric: fixing $N$ $CO_2$ obligately consumes $N$ RuBP and hence $N$ ATP via phosphoribulokinase, however fast each active site turns over.

#### Specificity is suppressed by $CO_2$ flooding

We added the oxygenase reaction (RBPO) and routed 2-phosphoglycolate through the host's native glycerate salvage (PGLYCP → GLYCTO1 → GLXCL → TRSARr → GLYCK), which loses one carbon as $CO_2$ per two oxygenations and consumes ATP and NAD(P)H, coupling oxygenation competitively to carboxylation at $\phi = v_{\text{oxy}}/v_{\text{carb}} = ([O_2]/[CO_2])/S_{c/o}$. The salvage introduces no free-energy artefact (the energy-generating-cycle test passes), and $\phi \to 0$ recovers the photorespiration-free baseline exactly.

Photorespiration is costly per event (+0.7% formate per percentage point of oxygenase flux; Figure S10c) but barely occurs here. FDH floods the cytoplasm with $CO_2$ (intracellular $[CO_2] > 500\ \mu\text{M}$), so against air-saturated $O_2$ (∼250 $\mu$M) the ratio is ≈0.5 or lower, making formatotrophy a built-in $CO_2$-concentrating mechanism. At that ratio the poor-specificity Form II enzyme *C. necator* carries ($S_{c/o} \approx$ 10–15) loses only ∼2–3.5% of formate, and engineering to a plant-like Form I ($S_{c/o} \approx$ 80–100) recovers only ∼3%. Specificity matters only under $CO_2$-limited conditions ($[O_2]/[CO_2] \gtrsim 1$), where the penalty exceeds 10% and low $S_{c/o}$ becomes infeasible, a regime avoided by formatotrophic $CO_2$ generation.

The record-turnover variant is strictly worse than the native enzyme for this application: it achieves its $k_{\text{cat}}$ by trading away specificity and is extremely $O_2$-sensitive[75], so its turnover advantage is worthless where turnover is non-binding while its degraded specificity would *add* photorespiratory cost.

#### Implication

RuBisCO turnover and specificity are therefore poor engineering levers here because the CBB disadvantage is energetic: the extra ATP and NADPH consumed per $CO_2$ fixed forces more formate to be oxidised for energy. This pathway stoichiometry cannot be engineered away at the catalyst, reinforcing the finding that pathway choice, not enzyme kinetics, sets the formate economy.

#### S14 Enzyme-cost accounting sensitivity of the pathway comparison

The relative performance of rGlyP against CBB in enzyme-constrained simulations depends on two enzyme costs not directly measured in *C. necator*: the subunit-mass charge for the glycine-cleavage system (GCS), and the reverse-turnover treatment for the rGlyP $C_1$/tetrahydrofolate reductive block. Both pathways were re-solved with each correction using `SupraModel.optimize()` (`PaperScripts/Supplementary/data/generate_rglyp_cost_sensitivity_rigorous.py`; outputs in `Results/rglyp_cost_sensitivity/rigorous_min_formate.csv` and `rigorous_max_growth.csv`).

##### The contested costs

*Glycine-cleavage system mass.* The GCS is a multi-subunit complex (P, T, H proteins) required by rGlyP. The baseline charges only one representative subunit; charging the full P+T+H complex raises the cost per unit GCS flux in proportion to the summed subunit mass, tightening the rGlyP proteome constraint without affecting CBB.

*Haldane reverse-$k_{cat}$ exemption.* Reversible reactions are split into half-reactions drawing independently on the pool, with reverse turnover set to $0.1 \times k_{cat,fwd}$ as a uniform Haldane approximation[30]. The baseline exempts five biosynthetic reactions of the rGlyP reductive $C_1$/THF block: formate activation, methenyl-cyclohydrolase, methylene-THF dehydrogenase, glycine cleavage and serine hydroxymethyltransferase. They run strongly in the thermodynamically favoured direction under formatotrophic conditions, supported by the thermodynamic and concentration analyses (Section S8). Removing the exemption forces each to carry a nominal reverse draw, penalising rGlyP exclusively since CBB has no analogous exempted block.

*Expression derate and CBB counter-correction.* A further scenario derates the $k_{cat}$ of the three heterologous rGlyP enzymes (Fch, GART, Ftl) by $\times 0.3$, reflecting suboptimal expression in a non-native host; this too penalises only rGlyP. A CBB-penalising counter-correction charges the full $L_8S_8$ RuBisCO holoenzyme rather than the large-subunit mass, tightening the CBB pool because RuBisCO is its largest single allocation.

##### Free growth: no superiority claim

At maximum-$\mu$ the baseline predicts rGlyP growing +29% faster than CBB on the same formate uptake. This is *not* used as a claim in the main text, and the sensitivity analysis shows why. Removing the Haldane exemption alone reverses +29% to −37%, because the rGlyP $C_1$ block then draws forward and nominal-reverse enzyme simultaneously. The $\times 0.3$ derate collapses the advantage to −0.5%; the full-complex GCS mass alone reduces it to +10%; stacking all three rGlyP penalisations gives −77%. Only the CBB-penalising RuBisCO correction increases it. The free-growth result is therefore determined by whether the $C_1$ block is exempted, a modelling assumption rather than a measured property, so no maximum-growth superiority claim is made.

##### Fixed growth: the formate saving persists

At $\mu = 0.126$ h$^{-1}$ with formate minimised, the baseline predicts a +15.2% saving for rGlyP (28.9 against 34.1 mmol gDW$^{-1}$ h$^{-1}$). None of the four individual corrections (Haldane penalty, full-complex GCS mass, ×0.3 derate or the CBB-penalising RuBisCO correction) makes rGlyP infeasible or reverses the saving, which ranges from 10.4% to 20.3%.

The underlying stoichiometric advantage (lower per-carbon ATP and NADPH burden) persists as a floor: enzyme-cost corrections can force the LP onto less efficient routes, but so long as rGlyP remains feasible no single correction erases it.

##### Adversarial stacked limit

Only when full-complex GCS mass, the Haldane penalty and the expression derate are imposed *simultaneously* does rGlyP become infeasible: combined pool demand exceeds the enzyme budget. This worst case requires three independent conservative assumptions to hold at once, so it is diagnostic rather than predictive. It identifies full-complex GCS mass combined with expression derate as the primary sensitivity knob, making direct *C. necator* rGlyP proteomics and in vivo $k_{\mathrm{cat}}$ measurement for the heterologous enzymes the most useful follow-ups.

GAM (135.0 mmol ATP gDW$^{-1}$) and NGAM (3.0 mmol ATP gDW$^{-1}$ h$^{-1}$) are held identical between pathways, so no observed difference reflects a maintenance assumption. Individual corrections, maintenance parity and saturation policy (Section S4.8) preserve the sign of the +15.2% advantage; only the stacked limit removes it. The separate folate-pool turnover stress test (Figure S8) identifies a small-pool, low-turnover corner where rGlyP's higher C1 throughput would become limiting.

##### Setpoint independence of the fixed-growth advantage

Re-solving from $\mu = 0.05$ to 0.15 h$^{-1}$ shows that the fixed-growth result does not depend on the 0.126 h$^{-1}$ setpoint (70% of the 0.18 h$^{-1}$ wild-type batch ceiling): the saving stays within 14.1–15.8% (Table S8), including 14.6% at $\mu \approx 0.07$ h$^{-1}$, the rate the current engineered rGlyP strain can reach. In this supplementary mechanistic sweep only, 100 PPFD uses the full nominal pump budget rather than the main text's 45% occupancy reference. The resulting rhodopsin saving is larger for rGlyP at every setpoint (52.9% against 42.6% at $\mu = 0.07$ h$^{-1}$; 26.3% against 16.8% at $\mu = 0.126$ h$^{-1}$); the absolute fraction falls with $\mu$ because a fixed ATP supply covers a smaller share of a larger demand. Reproduced by `PaperScripts/metabolic_audits/generate_setpoint_transhydrogenase_audit.py`; outputs in `Results/metabolic_audits/setpoint_advantage_sweep.csv`.

##### Soluble transhydrogenase sensitivity

Soluble NAD(P) transhydrogenase (`NADTRHD`) is blocked in the primary model (rhodopsin supplies ATP/PMF, not NADPH) while proton-coupled transhydrogenase remains available. Re-solving at $\mu = 0.126$ h$^{-1}$ with `NADTRHD` opened (±1000 mmol gDW$^{-1}$ h$^{-1}$) leaves dark demands unchanged

Table S8: **Setpoint robustness of the dark fixed-growth formate advantage.** (Related to Figure 2.) Enzyme-constrained minimum-formate demand (mmol $gDW^{-1}$ $h^{-1}$) for both pathways across the fixed-growth setpoint, with the relative rGlyP advantage (1 − rGlyP/CBB).

| $\mu$ ($h^{-1}$) | rGlyP | CBB | rGlyP advantage |
|---|---|---|---|
| 0.05 | 12.50 | 14.55 | 14.1% |
| 0.07 | 16.81 | 19.68 | 14.6% |
| 0.10 | 23.28 | 27.39 | 15.0% |
| 0.126 | 28.89 | 34.06 | 15.2% |
| 0.15 | 34.06 | 40.45 | 15.8% |

(rGlyP 28.89, CBB 34.06 mmol $gDW^{-1}$ $h^{-1}$; advantage 15.19% in both cases). Under the same full-pump-budget supplementary setting, the rhodopsin saving at exactly 100 PPFD is likewise essentially unchanged (rGlyP 26.3% under both; CBB 16.8% closed against 17.2% open). The differentiation is governed by absolute ATP demand, not by soluble transhydrogenation. Outputs in `Results/metabolic_audits/transhydrogenase_open_sensitivity.csv`.

#### S15 Sensitivity to the respiratory membrane footprints

The inner-membrane area constraint (Section S4) prices two sectors: light-harvesting modules, whose footprints follow from the rhodopsin monomer and photosystem cross-sections, and integral-membrane respiratory complexes, whose footprints are estimated from published structural dimensions. The second set is the weaker input: a cryo-EM structure fixes the transmembrane cross-section to within a factor below two, but converting that to an area *cost per unit flux* also invokes the turnover number, and neither the lipid annulus nor supercomplex organisation is represented. The respiratory draw is therefore an order-of-magnitude estimate.

A single multiplicative factor $s$ scales all respiratory footprints together, leaving the light-harvesting footprints and the 20 $m^2$ $gDW^{-1}$ budget untouched. At each level the current model is re-run: fixed-growth minimum-formate sweep, then five-variable design optimisation for the external cylinder with rGlyP. At $s = 0$ the respiratory complexes are unpriced; $s = 1$ is the structural estimate adopted throughout. The sweep spans to $s = 4$, covering more than a decade in the respiratory draw.

Table S9: **Respiratory footprint level against fixed-growth formate demand and optimised route efficiency.** (Related to Fig. 4.) Every row is regenerated from the current model. Formate demand is shown in the dark and at exactly 100 PPFD. Efficiencies are capture-free route-boundary external-cylinder rGlyP optima at the full-membrane ceiling, used here to isolate the footprint effect; the main text instead uses 45% occupancy on the closed, capture-charged boundary. The thermodynamic branch is formate-feed-bound at the 50-PPFD lower limit throughout, so its 600-nm boundary coordinate is not reported as a spectral preference.

| $s$ | Dark formate | Formate at 100 PPFD | $\eta$, SOTA | $\eta$, therm. limit | SOTA $\lambda^\star$ |
|---|---|---|---|---|---|
| | (mmol gDW$^{-1}$ h$^{-1}$) | | (%) | (%) | (nm) |
| 0 | 28.89 | 19.49 | 7.33 | 11.93 | 556.9 |
| 0.5 | 28.89 | 20.33 | 7.18 | 11.93 | 556.8 |
| 1 | 28.89 | 21.29 | 7.09 | 11.92 | 556.3 |
| 2 | 28.89 | 23.74 | 7.03 | 11.91 | 556.2 |
| 4 | 30.17 | 27.78 | 6.60 | 11.38 | 555.2 |

Machine-readable output: `Results/reactor_geometry_wavelength/respiratory_footprint_sensitivity.csv`.

*Membrane competition.* A larger respiratory draw leaves less area for the pump, so more formate must be oxidised to supply ATP. Dark demand is unchanged through $s = 2$, when the respiratory sector still fits without a competing pump. At the deliberately extreme $s = 4$ level, respiratory occupancy itself becomes restrictive and dark formate demand rises by 4.4%. The effect enters $\eta$ through the fixed-growth yield, not through a separate optical term.

*Efficiency effect.* The eight-fold range $s = 0.5$–$4$ lowers the state-of-the-art optimum by 0.58 percentage points and the thermodynamic-limit optimum by 0.55. Around the adopted $s = 1$ estimate, halving the footprint returns 0.10 points and doubling it costs 0.05 at the state of the art; the corresponding thermodynamic-limit changes are at most 0.02 points. Most of the remaining loss occurs at $s = 4$, where the dark state also becomes membrane-limited.

*Spectral effect.* The state-of-the-art optimum moves only 1.7 nm over the full range, from 556.9 to 555.2 nm. The wavelength is set mainly by photon energy against pump yield (Section S6.12); membrane pricing changes the amount of pump that can be used much more than the preferred band. The thermodynamic branch is not evidence for a red preference: it remains at the minimum PPFD and the wavelength search boundary while volumetric formate feed binds.

*Unpriced-respiration comparison.* Setting $s = 0$ raises the current state-of-the-art route optimum from 7.09% to 7.33%; the thermodynamic-limit optimum changes by less than 0.01 percentage points. The occupancy condition is a separate axis: the main text reports 45% occupancy on the closed boundary, where the state-of-the-art retinal-only optimum is 3.29% (Table S27b). This sensitivity therefore supports the direction and scale of the membrane-pricing effect without substituting its route-boundary values for the headline process result.

The priced set includes FDH2, Complex I, the terminal oxidases and ATP synthase, but the reconstruction does not resolve aggregate membrane occupancy for the formate, gas, water and ion transporters that also carry flux at the setpoint. Pricing those carriers would reduce the area available to rhodopsin and push in the same direction as increasing $s$. This unresolved occupancy is why the main text uses 45% as the reference allocation and treats the full membrane budget as a ceiling.

### Tables

**Efficiency definitions and main boundary accounting.** Moved here from the main text to keep the main-text figure-plus-table count within the journal's limit of seven combined; both are cited from the Results as Table S10 and Table S11.

Table S10: **Energy efficiencies analysed.** (Related to Figs. 1 and 7.) All three divide biomass chemical power by the specified solar input. Electrical demand per mole of formate comprises electrosynthesis energy $E_{synth}$ and $CO_2$ capture energy $E_{cap}$ (Methods). Direct-air-capture demand is 419.9 kJ mol$^{-1}$ at the state of the art and 19.16 kJ mol$^{-1}$ at the thermodynamic limit; alternative $CO_2$ sources are sensitivities (Table S19). The headline process efficiency includes sunlight incident on the whole collector aperture, with no export credit for unused energy. This is termed the closed energy-accounting boundary; it does not denote a physically closed reactor or imply an "open boundary" scenario. Cell-scale and fixed-geometry diagnostics include only the solar inputs specified in their rows. Crop comparisons use the whole-collector efficiency at the design irradiance, with the temporal and site-area limitations described in the Discussion.

| Quantity | Optimised over | Denominator charges | $CO_2$ capture, $E_{cap}$ | Reported in |
|---|---|---|---|---|
| $\eta_{cell}$ | cell flux and wavelength for reported maxima | absorbed in-band light, corrected for dichroic loss, plus solar input for formate ($E_{synth}$ only) | not charged | Sec. S6.12 |
| $\eta$, diagnostic | nothing; fixed $V/A$, 568 nm | in-band photons delivered to the culture, plus the photovoltaic branch ($E_{synth}$ + $E_{cap}$), at one fixed bioreactor geometry | charged | Fig. 6 |
| $\eta$, closed boundary | design variables jointly (Methods) | sunlight incident on the whole collector aperture, including any unused spectrum | charged | Fig. 7, Table S11 |

Table S11: **Process-efficiency accounting for the rGlyP optimum.** (Related to Figure 7.) Deterministic efficiencies include all incident collector sunlight and $CO_2$ capture energy. The illuminated reference caps rhodopsin occupancy at 45%. The dark comparators either omit the optical collector and use standard photovoltaics, or retain its aperture. The auxiliary-load case re-optimises the design with mixing energy included. The final row allows the full membrane budget and is a model ceiling rather than a demonstrated allocation.

| Accounting case | SOTA $\eta$ (%) | Theoretical $\eta$ (%) | Interpretation |
|---|---|---|---|
| Whole collector, retinal-only, $CO_2$ capture included | 3.29 | 10.89 | Propagated intervals: 2.73–3.87 (median 3.23) and 9.09–12.24 (median 10.53); heating value and dichroic transmission included (Methods) |
| dark comparator, standard PV (no aperture built) | 2.56 | 11.15 | Light adds 0.73 points at the state of the art; at the limit the dark branch is better |
| dark comparator, aperture PV (aperture still charged) | 2.14 | 10.86 | Light gains 1.16 and 0.03 points |
| Whole collector, joint carotenoid–retinal | 3.52 | 10.70 | Paired antenna $\Delta\eta$: +0.107 pp (−0.034 to +0.284; 93.2% positive) and −0.178 pp (−0.216 to −0.140; 0% positive); the light-enabled optimum selects dark at the limit |
| Closed boundary plus low-head auxiliary load ($\sim$0.35 W $L^{-1}$) | 3.26 | 10.44 | Duty inside the objective, so the design thins to pay it; no photovoltaic surplus exists to absorb it |
| Full-membrane ceiling, closed boundary | 3.74 | 10.90 | Rhodopsin occupancy about twice the purple-membrane reference; Table S27 |

Table S12: **Experimental data for bacteriorhodopsin photophysics model calibration.** Sources: Xie[63] ($\varphi_1$, 110 K); Logunov[14] ($\Phi_{iso}$, RT); Marinetti[60] ($\varphi_t$, RT, E. coli); wavelengths from Karasuyama et al.[76] (Table MOESM3). † = wavelength estimated from corresponding Asn variant. Marinetti WT $\lambda_{max} = 560$ nm (E. coli lipid). Predictions in Figure S14.

| Study | WT? | Variant | QY | QY type | $\lambda_{max}$ (nm) | Condition |
|---|---|---|---|---|---|---|
| *Xie (1990) – WT bR, 110 K* | | | | | | |
| [63] | Yes | WT | $0.609 \pm 0.022$ | $\varphi_1$ | 420 ($\lambda_{exc}$) | 110 K |
| [63] | Yes | WT | $0.553 \pm 0.020$ | $\varphi_1$ | 562 ($\lambda_{exc}$) | 110 K |
| [63] | Yes | WT | $0.544 \pm 0.036$ | $\varphi_1$ | 600 ($\lambda_{exc}$) | 110 K |
| *Logunov et al. (1996) – $\Phi_{iso}$, RT* | | | | | | |
| [14] | Yes | wbR | $0.58 \pm 0.05$ | $\Phi_{iso}$ | 568 | RT, pH 7 |
| [14] | No | D212N | $0.70 \pm 0.05$ | $\Phi_{iso}$ | 570 | RT, pH 7 |
| [14] | No | R82Q | $0.65 \pm 0.15$ | $\Phi_{iso}$ | 590 | RT, pH 7 |
| [14] | No | blue bR | $0.70 \pm 0.15$ | $\Phi_{iso}$ | 605 | RT, pH 7 |
| [14] | No | D85N | $0.75 \pm 0.15$ | $\Phi_{iso}$ | 610 | RT, pH 7 |
| *Marinetti et al. (1989) – $\varphi_t$, RT, E. coli* | | | | | | |
| [60] | Yes | wt | 0.63 / 0.67 | $\varphi_t$ | 560 | RT, pH 6 / 7 |
| [60] | No | D85N | 0.03 / 0.21 | $\varphi_t$ | 613 | RT, pH 6 / 7 |
| [60] | No | D85E | 0.23 / 0.54 | $\varphi_t$ | 615 | RT, pH 6 / 7 |
| [60] | No | D96N | 0.78 / 0.57 | $\varphi_t$ | 560 | RT, pH 6 / 7 |
| [60] | No | D96E | 0.61 / 0.58 | $\varphi_t$ | $\sim 560^{\dagger}$ | RT, pH 6 / 7 |
| [60] | No | D115N | 0.35 | $\varphi_t$ | 566 | RT, pH 6 |
| [60] | No | D115E | 0.18 / 0.30 | $\varphi_t$ | $\sim 566^{\dagger}$ | RT, pH 6 / 7 |
| [60] | No | D212N | 0.14 | $\varphi_t$ | 580 | RT, pH 6 |
| [60] | No | D212E | 0.36 | $\varphi_t$ | $\sim 580^{\dagger}$ | RT, pH 6 |

$\varphi_t$=proton translocation; $\Phi_{iso}$=photoisomerisation; $\varphi_1$=forward (bR→K).
/ separates pH 6 and pH 7. † wavelength from corresponding Asn variant.

Table S13: **Glossary of model terms and symbols.** Symbols grouped by model layer. Default flux units: mmol $gDW^{-1}$ $h^{-1}$ (normalised to biomass *present*; related to per-gDW-formed by $\mu$, e.g. 31.5 mmol $h^{-1}$ = 250 mmol per gDW formed at $\mu = 0.126$).

| **Term / symbol** | **Layer** | **Definition** |
|---|---|---|
| PPFD | Light input | Incident photon flux density ($\mu$mol photons $m^{-2}$ $s^{-1}$). |
| $\lambda_{max}$ | Photophysics | Rhodopsin absorption maximum. |
| $\Phi_{QY}$, $\Phi_{iso}$ | Photophysics | Quantum yield / photoisomerisation yield. |
| $\tau$ | Photocycle | Mean productive photocycle time (Poisson pump model). |
| $\sigma_0$ | Optics | Molecular absorption cross-section (WT extinction coefficient). |
| $\sigma_{eff}$ | Optics–metabolism | Effective cross-section ($4.5\sigma_0$); maps PPFD to pump flux. |
| $k_{abs}$ | Optics | Photon absorption rate per molecule ($\sigma_{eff} \times$ local flux). |
| $P_{act}$ | Pump model | Pump probability per photocycle, $1 - \exp(-k_{abs}\Phi_{QY}\tau)$. |
| $R_{pump}$, $r_{pump}$ | Pump model | Proton-pump capacity (one $H^+$ per photocycle). |
| $G(z)$, $G(r)$ | Reactor optics | Local scalar irradiance (flat-panel depth $z$, cylindrical radius $r$). |
| $a(\lambda)$, $s(\lambda)$ | Reactor optics | Absorption and scattering coefficients (two-flux model). |
| $\delta$ | Reactor optics | Effective attenuation; $\delta = \sqrt{a(a + 2bs)}$, $b$=backscatter fraction. |
| $\mu$ | Metabolism | Specific growth rate ($h^{-1}$; 0.126 in fixed-growth). |
| $v_{bio}$ | Metabolism | Biomass flux ($\equiv \mu$). |
| $Y$ | Yield | Biomass yield (g $mol^{-1}$ formate or g $g^{-1}$; basis varies by table). |
| rGlyP | Pathway | Reductive glycine pathway (direct C1 formate assimilation). |
| CBB | Pathway | Calvin–Benson–Bassham cycle ($CO_2$ fixation after formate oxidation). |
| EC-FBA / RBA | Metabolism | Enzyme-constrained flux balance / resource-balance analysis. |
| $P_{total}$, $f_{enzyme}$ | Proteome | Total protein and catalytic enzyme fractions. |
| $\sigma_j$ | Proteome | Enzyme saturation for reaction $j$ (not optical $\sigma_0$). |
| $\pi_i$ | LP duals | Shadow price for constraint $i$; meaning depends on objective. |

Table S14: **Principal model parameters.** Values are the main-analysis settings unless otherwise noted.

| Parameter | Value | Use / note | Source |
|---|---|---|---|
| WT $\lambda_{max}$ | 568 nm | Calibration anchor. | [14,63] |
| Main wavelength range | 450–600 nm | Main text; 450–700 nm in supplement. | Operating choice |
| WT QY calibration target | $0.64 \pm 0.04$ | Likelihood target. | [5,11] |
| QY posterior median | ∼0.60 at WT | Propagated to pump rate. | MCMC |
| Photocycle calibration | $\tau_M \approx 5$ ms, $\tau_{total} \approx 15$ ms | Seven-state timing. | [16] |
| Molecular cross-section $\sigma_0$ | $2.41 \times 10^{-20}$ $m^2$ | From WT extinction coefficient. | [77] |
| Effective cross-section $\sigma_{eff}$ | $4.5\sigma_0$ | Lumped calibration factor; sensitivity parameter. | This work |
| Pump activation | $1 - e^{-k_{abs}\Phi_{QY}\tau}$ | Poisson model. | This work |
| $H^+$ per pump event | 1 | Per productive photocycle. | [6] |
| ATP synthase coupling | 4 $H^+$ $ATP^{-1}$ | Rhodopsin $H^+$ to ATP. | [78] |
| Fixed-growth setpoint | 0.126 $h^{-1}$ | 70% of 0.18 $h^{-1}$ reference. | Operating choice |
| Envelope sweep | Formate 2–50, PPFD 0–700 | Growth–yield frontiers. | Operating choice |
| Fixed-growth sweep | Fixed $\mu = 0.126$; min formate | Main analysis mode. | Operating choice |
| Fixed-dark-formate sweep | Fixed dark formate; max growth | Light-response diagnostic. | Operating choice |
| GAM | 135.0 mmol ATP $gDW^{-1}$ | Growth-associated maintenance. | [23]; corroborated by[74] |
| NGAM | 3.0 mmol ATP $gDW^{-1}$ $h^{-1}$ | Maintenance ATP hydrolysis. | [22] |
| Passive proton leak | 1.0 mmol $H^+$ $gDW^{-1}$ $h^{-1}$ | PMF dissipation. | [22] |
| $P_{total}$ | 0.68 g $gDW^{-1}$ | Proteome budget. | [23] |
| $f_{enzyme}$ | 0.50 | Catalytic fraction (0.34 g $gDW^{-1}$). | [23] |

*Table S14 continued*

| Parameter | Value | Use / note | Source |
|---|---|---|---|
| Rhodopsin MW | 27 kDa | Proteome allocation. | Sequence-derived |
| Rhodopsin $k_{\mathrm{cat}}$ | Physics-derived | $\sim$3.85 $s^{-1}$ at WT, 100 PPFD. | Model-derived |
| Rhodopsin footprint | 7 $nm^2$ | Membrane occupancy. | Structural estimate |
| Membrane budget | 20 $m^2$ $gDW^{-1}$ | Inner-membrane occupancy limit. | [23,39] |
| Rhodopsin membrane cap | 1.0 of budget | **Upper bound**; native packing 0.4–0.6; cap sweep in Table S27. | Design choice |
| Baseline PPFD | 100 $\mu$mol $m^{-2}$ $s^{-1}$ | Flat-panel reference. | Operating choice |
| Path length | 5 cm | Flat-panel reference. | Operating choice |
| Biomass density | 0.5 gDW $L^{-1}$ | Optical reference. | Operating choice |
| Rhodopsin abundance | $5.0 \times 10^{-6}$ mol $gDW^{-1}$ | Self-shading input. | Model-derived |
| Spectral FWHM | 100 nm | Absorption-band width. | Assumed |
| Backscatter $b$ | 0.01 | Two-flux model. | Assumed; swept |
| Scattering coeff. | $0.8(\lambda/550)^{-1}$ | Power-law scattering. | [46,47] |
| QY MCMC | 32 walkers; 300 burn-in; 800 retained | 25,600 post-burn-in draws. | Operating choice |
| MCMC diagnostics | $\tau_{\mathrm{int}}$ = 61–69; ESS 371–423; $\hat{R}$ = 1.00–1.03; acc. 0.22 | Posterior convergence. | Model-derived |
| Solver tolerances | Feas. $10^{-6}$; opt. $10^{-7}$; RBA $10^{-4}$ | GLPK/RBA convergence. | |

Table S15: **Reactions constrained to zero or directionally capped.** Applied via `apply_standard_constraints()` before pathway modifications. $(0, 0)$=disabled; $(0, 1000)$=forward-only; $(-100, 0)$=reverse-only.

| **Reaction ID** | **Reaction / description** | **Constraint** | **Rationale** |
|---|---|---|---|
| *Transport – substrates absent from formate medium* | | | |
| THMDt2 | Thymidine transport | $(0, 0)$ | Absent |
| THRtr | Threonine transport | $(0, 0)$ | Absent |
| PROt4 | Proline transport | $(0, 0)$ | Absent |
| SERt4 | Serine transport | $(0, 0)$ | Absent |
| GLUt4 | Glutamate transport | $(0, 0)$ | Absent |
| INSt2 | Inosine transport | $(0, 0)$ | Absent |
| ADNt2 | Adenosine transport | $(0, 0)$ | Absent |
| URAt2 | Uracil transport | $(0, 0)$ | Absent |
| URIt2 | Uridine transport | $(0, 0)$ | Absent |
| CYTDtr | Cytidine transport | $(0, 0)$ | Absent |
| CYTDt2 | Cytidine transport (alt.) | $(0, 0)$ | Absent |
| ASO3t2 | Arsenite transport | $(0, 0)$ | Absent |
| BENZOTt | Benzoate transport | $(0, 0)$ | Absent |
| GLUABUTt7 | Glu/4-aminobutyrate antiport | $(0, 0)$ | Absent |
| CITt7 | Citrate transport | $(0, 0)$ | Absent |
| ASPALAt | Asp/Ala antiport | $(0, 0)$ | Absent |
| NAt3_1g | $Na^+$ transport (electrogenic) | $(0, 0)$ | Electroneutral exchange used |
| *Unidirectional reactions* | | | |
| PTAr | Phosphotransacetylase (rev.) | $(0, 1000)$ | Forward only; futile cycle |
| PPAKr | Polyphosphate kinase (rev.) | $(0, 1000)$ | Synthesis only |
| ALCDgl | Alcohol dehydrogenase (glycolaldehyde) | $(0, 100)$ | Forward only |
| ALCD19 | Alcohol dehydrogenase (alt.) | $(-100, 0)$ | Reverse only |
| *Inactive under formate* | | | |
| FOMETRi | Formate–methanol exchange | $(0, 0)$ | No methanol |
| SUCCtr | Succinate transport (efflux) | $(0, 0)$ | Not secreted |
| 3HBCDH | 3-HB-CoA dehydrogenase | $(0, 0)$ | PHB branch inactive |
| ICL | Isocitrate lyase (glyoxylate) | $(0, 0)$ | Inactive on C1 |
| HACD1 | 3-HA-CoA dehydrogenase | $(0, 0)$ | $\beta$-oxidation inactive |
| MDH2 | Malate DH (periplasmic) | $(0, 0)$ | Cytoplasmic MDH sufficient |
| POX | Pyruvate oxidase | $(0, 0)$ | Not expressed |
| MGSA | Methylglyoxal synthase | $(0, 0)$ | Bypass inactive |
| THRA | Threonine aldolase | $(0, 0)$ | No substrate |

*Table S15 continued*

| **Reaction ID** | **Reaction / description** | **Constraint** | **Rationale** |
|---|---|---|---|
| ALRTg | Aldehyde reductase | $(0,0)$ | Inactive |
| ALRTgp | Aldehyde reductase (alt.) | $(0,0)$ | Absent from proteome |
| G3PD2 | G3P dehydrogenase (FAD) | $(0,0)$ | No glycerol |
| HPYRI | Hydroxypyruvate isomerase | $(0,0)$ | No serine |
| ISOVC | Ascorbate cycle | $(0,0)$ | No ascorbate |
| P5CD4 | P5C dehydrogenase (4) | $(0,0)$ | Pro catabolism inactive |
| P5CD5 | P5C dehydrogenase (5) | $(0,0)$ | Pro catabolism inactive |
| *Adenylate kinase isoforms* | | | |
| ADK3 | Adenylate kinase 3 | $(0,0)$ | Prevents ATP–AMP cycling |
| ADK4 | Adenylate kinase 4 | $(0,0)$ | Inactive |
| *Nucleotide/cofactor* | | | |
| PTA2 | Phosphotransacetylase 2 | $(0,0)$ | Duplicate |
| ADPT | Adenine PRT | $(0,0)$ | No adenine |
| ADPRT3 | ADP-ribosyltransferase 3 | $(0,0)$ | Not expressed |
| ADPRT4 | ADP-ribosyltransferase 4 | $(0,0)$ | Not expressed |
| PCT1 | Pantothenate kinase 1 | $(0,0)$ | Prevents duplicate CoA |
| NADTRHD | NAD(P) transhydrogenase | $(0,0)$ | Isolates NADPH balance |
| CYTTS1 | Cyt $c$ oxidase step 1 | $(0,0)$ | Isoform inactive |
| CYTTS3 | Cyt $c$ oxidase step 3 | $(0,0)$ | See CYTTS1 |
| CYTTS5 | Cyt $c$ oxidase step 5 | $(0,0)$ | See CYTTS1 |
| *Exchange fluxes – not supplied/secreted* | | | |
| EX_acal_e | Acetaldehyde exchange | $(0,0)$ | Not secreted |
| EX_acac_e | Acetoacetate exchange | $(0,0)$ | Not secreted |
| EX_pyr_e | Pyruvate exchange | $(0,0)$ | Not secreted |
| EX_cit_e | Citrate exchange | $(0,0)$ | Not secreted |
| EX_icit_e | Isocitrate exchange | $(0,0)$ | Not secreted |
| EX_fum_e | Fumarate exchange | $(0,0)$ | Not secreted |
| EX_mlt_e | Malate exchange | $(0,0)$ | Not secreted |

**Constrained-optimal designs.** Five-variable optimisation ($\lambda$, CDW, $V/A$, PPFD, $f_{\text{mem}}$) subject to $O_2$, formate-feed and solar-concentration limits (Table S22; solver in Section S6). At external-reactor optima, solar concentration binds; $O_2$ and formate are slack. CDW* and $V/A^*$ sit on an iso-efficiency ridge ($\eta$ nearly degenerate at fixed areal optical depth $\propto$ CDW $\times$ $V/A$).

**Energy-conversion benchmark boundaries.** Table S18 defines the quantities displayed in Fig. 7f and includes their time and area bases. Values from this work use the capture-charged whole-collector

Table S16: **Enzyme turnover numbers ($k_{cat}$) used in the GECKO proteome-pool constraint.** Per-EC-class distribution of the 1317 enzyme/reaction entries in the kinetic database. Turnover numbers and molecular weights are TurNuP deep-learning predictions[31] keyed to *C. necator* H16 genes, except the heterologous rGlyP enzymes and reaction-level curated overrides listed below. The complete per-enzyme/reaction table is provided as supplementary data (`enzyme_kcat_table.csv`).

| EC class | $n$ | median $k_{cat}$ ($s^{-1}$) | min | max |
|---|---|---|---|---|
| 1 Oxidoreductases | 408 | 10.6 | 4.11 | 371 |
| 2 Transferases | 270 | 7.2 | 5.00 | 200 |
| 3 Hydrolases | 190 | 13.6 | 3.60 | 764 |
| 4 Lyases | 199 | 9.4 | 2.25 | 213 |
| 5 Isomerases | 55 | 18.5 | 5.10 | 4300 |
| 6 Ligases | 106 | 4.6 | 2.10 | 116 |
| 7 Translocases | 19 | 60.0 | 60.00 | 300 |

**Provenance:** 1306 TurNuP deep-learning prediction; 4 curated_complex_level; 2 literature_Bifidobacterium_Xfpk; 1 Marx2003_Mextorquens_AM1_purified_SA117Umg_derived; 1 Etzold_1997_EurJBiochem_243_336; 1 Pomper1999_Vmax_derived__Claassens2020_pathway__UniProt_Q49135; 1 Marolewski1997_Ecoli_PurT_homolog_transfer__UniProt_Q0KAH8; 1 shares_rubisco_holoenzyme_with_RBPC.

**Heterologous / manually-curated entries:**

| Reaction | gene/UniProt | $k_{cat}$ ($s^{-1}$) | MW (kDa) | source |
|---|---|---|---|---|
| Ftl | Ftl | 115.8 | 59.4 | Marx2003_Mextorquens_AM1_purified_SA117Umg_derived |
| ATPS4m | reaction:ATPS4m | 270.0 | 530.0 | Etzold_1997_EurJBiochem_243_336 |
| FDH | reaction:FDH | 200.0 | 190.2 | curated_complex_level |
| Fch | Q49135 | 120.0 | 21.7 | Pomper1999_Vmax_derived__Claassens2020_pathway__UniProt_Q49135 |
| GART | Q0KAH8 | 37.6 | 42.8 | Marolewski1997_Ecoli_PurT_homolog_transfer__UniProt_Q0KAH8 |
| NADH16 | reaction:NADH16 | 100.0 | 550.0 | curated_complex_level |
| NADH5 | reaction:NADH5 | 100.0 | 550.0 | curated_complex_level |
| NADHDH | reaction:NADHDH | 100.0 | 550.0 | curated_complex_level |
| PKETF | reaction:PKETF | 10.0 | 185.0 | literature_Bifidobacterium_Xfpk |
| PKETX | reaction:PKETX | 10.0 | 185.0 | literature_Bifidobacterium_Xfpk |
| RBPO | reaction:RBPO | 3.4 | 560.0 | shares_rubisco_holoenzyme_with_RBPC |

calculation. Canopy-intercepted and brief-peak crop values are identified separately.

Table S18: **System boundaries for energy-conversion benchmarks.** (Related to Fig. 7f.)

| Comparator | Counted boundary | Benchmark metric | Interpretation / exclusions |
|---|---|---|---|
| $C_3$ crops | Incident sunlight over a growing season to biomass, with biomass fractions dependent on source | Seasonal incident: 2.2%/3.6% (present/projected)[79]. Ceiling 4.6%[80,81] | Biomass recovery is excluded. |
| $C_4$ crops | Incident sunlight over a growing season to biomass, with biomass fractions dependent on source | Seasonal incident: 3.0%/4.1% (present/projected)[79]. Ceiling 6%[80,81] | The 4.3% value is brief-peak and canopy-intercepted. |
| Microalgae | Sunlight to algal biomass | 1.5–5.0%[47,81] | PBR benchmark; harvest excluded. |

| Comparator | Counted boundary | Benchmark metric | Interpretation / exclusions |
|---|---|---|---|
| PV–electrocatalytic $CO_2$ reduction | Solar PV to product | 10–15%[51] | Excludes biology. |
| PV–$H_2$ microbial protein | Solar PV to $H_2$ microbial protein | >10× protein vs soybean[82]; DAC-$H_2$ protein ~18.3 kWh $kg^{-1}$[83] | Closest power-to-biomass class; areal output metric. |
| PV–$H_2$ plus *C. necator* biosynthesis | $H_2$ to *C. necator* products | $CO_2$ reduction exceeding photosynthesis[84] | Host-lineage neighbour; product-specific allocation prevents direct ranking. |
| Electro-acetate-to-food / biomass | Electrochemical acetate to growth | Electrogenerated acetate[85] | $C_2$ route; organism-specific ranking needed. |
| Dark PV–formate bioconversion | All incident PV solar input, including $CO_2$ capture; no rhodopsin | 2.56% SOTA and 11.15% thermodynamic limit (rGlyP), this model | Standard-PV dark comparator; nutrients, mixing, capital and seasonal operation excluded. |
| Carotenoid–rhodopsin PV–formate bioconversion | Steady design irradiance over the whole collector aperture, including $CO_2$ capture | 3.52% SOTA and 10.70% imposed-light thermodynamic result at 45% occupancy and 70% antenna transfer | Optimised model predictions conditional on high pump expression and antenna assembly; the light-enabled optimum selects dark at the limit. |
| Microbial electrosynthesis / direct electrofermentation | Electrical input to cells/electrodes | No solar-to-biomass metric | Boundary context; TEA/LCA needed before ranking. |

**Carbon-capture basis.** Table S19 reports every capture basis (headline rows are maturity-matched). Both the CSV and the LaTeX rows are generated by `PaperScripts/reactor_geometry/data/generate_capture_basis_table.py`; dichroic $t \in [0, 1]$ is free (Section S6).

Table S19: **Closed-boundary efficiency against the carbon-capture basis.** (Related to Fig. 7.) Capture is charged as PV electrical work; dichroic transmission is free. The dark column uses standard PV with no light-delivery aperture. The retinal and joint columns include this dark branch as a fallback, so enabling light cannot lower the optimum. The joint column re-optimises the model-fit 70% Kin4B8 case for the two SOTA supplies compared in the text. Bold, maturity-matched; dashes, not evaluated. †Search edge. rGlyP, external cylinder, 45% occupancy.

| Scenario | Capture basis | $E_{synth}$ | $E_{cap}$ (kJ $mol^{-1}$) | $E_{formate}$ | $\eta_{dark}$ | $\eta_{ret}$ (%) | $\eta_{ant70}$ | $\Delta\eta_{ant}$ (pp) | $\lambda^{\star}_{ret/ant}$ (nm) |
|---|---|---|---|---|---|---|---|---|---|
| SOTA | None ($CO_2$ unpriced) | 500.00 | 0.00 | 500.00 | 4.71 | 5.03 | n/a | n/a | 552/n/a |
| Thermo. | None ($CO_2$ unpriced) | 270.00 | 0.00 | 270.00 | 11.94 | 11.94 | n/a | n/a | 600†/n/a |
| SOTA | Flue gas, 14 wt% | 500.00 | 29.32 | 529.32 | 4.45 | 4.81 | 4.82 | +0.002 | 551/550 |
| Thermo. | Flue gas, 14 wt% | 270.00 | 29.32 | 299.32 | 10.77 | 10.77 | n/a | n/a | 600†/n/a |

| Scenario | Capture basis | $E_{\mathrm{synth}}$ | $E_{\mathrm{cap}}$ (kJ mol$^{-1}$) | $E_{\mathrm{formate}}$ | $\eta_{\mathrm{dark}}$ | $\eta_{\mathrm{ret}}$ (%) | $\eta_{\mathrm{ant70}}$ | $\Delta\eta_{\mathrm{ant}}$ (pp) | $\lambda^{\star}_{\mathrm{ret/ant}}$ (nm) |
|---|---|---|---|---|---|---|---|---|---|
| SOTA | DAC, $W_{\mathrm{min}}/\eta_{\mathrm{II}}$ | 500.00 | 95.79 | 595.79 | 3.95 | 4.41 | n/a | n/a | 550/n/a |
| Thermo. | DAC, $W_{\mathrm{min}}/\eta_{\mathrm{II}}$ | 270.00 | 95.79 | 365.79 | 8.81 | 8.81 | n/a | n/a | 600†/n/a |
| **SOTA** | **DAC, deployed** | **500.00** | **419.90** | **919.90** | **2.56** | **3.29** | **3.52** | **+0.235** | **550/552** |
| **Thermo.** | **DAC,** $W_{\mathrm{min}}$ ($\eta_{\mathrm{II}}=1$) | **270.00** | **19.16** | **289.16** | **11.15** | **11.15** | **n/a** | **n/a** | **600†/n/a** |

Charging deployed DAC reduces the retinal-only SOTA prediction from 5.03% to 3.29%. On a like-for-like $W_{\mathrm{min}}/\eta_{\mathrm{II}}$ basis, the 14 wt% point-source duty is 69% below the air-separation value. Under point-source supply, light raises efficiency from the no-aperture dark baseline of 4.45% to 4.82%, or 0.37 percentage points (8%); the antenna increment is less than 0.01 points. Point-source use requires co-location with a suitable stream and is not atmospheric removal. At the thermodynamic limit, the standard-PV dark branch is better than either imposed light-harvesting design.

**Boundary stress tests.** Table S20 re-optimises the rGlyP external-cylinder design at 45% occupancy with additional electrical loads. All rows use the headline whole-collector boundary with $CO_2$ capture charged. Values are from `Results/reactor_geometry_wavelength/hybrid_dichroic_led_dac.csv`.

Table S17: Constrained-optimal designs for every electrolyser × pathway × geometry. (Related to Figure 7.) Route boundary with $CO_2$ capture; closed-boundary counterparts in Table S27b.

| Electrolyser | Pathway | Geometry | $\lambda^\star$ (nm) | $f^*_{\mathrm{mem}}$ | CDW* (g $L^{-1}$) | PPFD* ($\mu$mol $m^{-2}$ $s^{-1}$) | $V/A^*$ (m) | $\eta_{\mathrm{route}}$ (%) |
|---|---|---|---|---|---|---|---|---|
| SOTA | rGlyP | Flat panel | 556.2 | 1.00 | 9.17 | 2056 | 0.0112 | 4.80 |
| SOTA | rGlyP | Cyl ext. | 556.3 | 1.00 | 1.56 | 2056 | 0.0659 | 4.80 |
| SOTA | rGlyP | Annular | 556.3 | 1.00 | 1.44 | 2056 | 0.0710 | 4.80 |
| SOTA | rGlyP | Bag (int.) | 550.0 | 1.00 | 3.19 | 2048 | 0.0301 | 3.17 |
| SOTA | CBB | Flat panel | 552.9 | 1.00 | 9.11 | 2052 | 0.0087 | 4.39 |
| SOTA | CBB | Cyl ext. | 552.9 | 1.00 | 1.11 | 2051 | 0.0711 | 4.39 |
| SOTA | CBB | Annular | 552.9 | 1.00 | 5.62 | 2052 | 0.0141 | 4.39 |
| SOTA | CBB | Bag (int.) | 550.0 | 1.00 | 1.70 | 2048 | 0.0574 | 2.49 |
| Theoretical | rGlyP | Flat panel | 557.0 | 1.00 | 8.30 | 2057 | 0.0136 | 10.75 |
| Theoretical | rGlyP | Cyl ext. | 557.0 | 1.00 | 1.17 | 2057 | 0.0959 | 10.76 |
| Theoretical | rGlyP | Annular | 557.0 | 1.00 | 5.20 | 2057 | 0.0216 | 10.75 |
| Theoretical | rGlyP | Bag (int.) | 563.1 | 1.00 | 3.80 | 2064 | 0.1000 | 10.21 |
| Theoretical | CBB | Flat panel | 553.3 | 1.00 | 8.66 | 2052 | 0.0096 | 8.98 |
| Theoretical | CBB | Cyl ext. | 553.3 | 1.00 | 8.66 | 2052 | 0.0096 | 8.98 |
| Theoretical | CBB | Annular | 553.3 | 1.00 | 6.72 | 2052 | 0.0124 | 8.98 |
| Theoretical | CBB | Bag (int.) | 585.2 | 1.00 | 2.65 | 50 | 0.1000 | 8.49 |

Table S20: **Auxiliary-load stress tests for the rGlyP external-light optimum.** Each electrical load is included in the objective and supplied from photovoltaics on the collector aperture. $CO_2$ capture and the 45% rhodopsin occupancy cap are retained in every case.

| Accounting case | Counted denominator / stress imposed | SOTA $\eta$ (%) | Theoretical $\eta$ (%) | Interpretation |
|---|---|---|---|---|
| Reference | Whole incident collector solar input, $CO_2$ capture included | 3.29 | 10.89 | No additional mixing or circulation load |
| Low-head auxiliary load | Reference plus 0.35 W $L^{-1}$ | 3.27 | 10.44 | Both scenarios re-optimised with the load |
| Pumped long-loop stress | Reference plus 1.7 W $L^{-1}$ | 3.18 | 8.99 | Both scenarios re-optimised with the load |

**Optical-hardware stress tests.** Table S21 retains a separate optical sensitivity calculation for the rGlyP external cylinder, using the full membrane budget and a capture-free route denominator. It does not quantify sensitivity of the headline 45%-cap, capture-charged whole-collector optimum. Values are from `Results/reactor_geometry_wavelength/optical_hardware_stress.csv`.

Table S21: **Optical-hardware diagnostics (re-optimised, capture-free route accounting, full membrane budget).** These values are not the headline process efficiency and cannot be used for its crop comparison. (Related to Figure 7.)

| Stress case | Dichroic | Band (nm) | Max conc. | SOTA $\eta$ (%) | Theoretical $\eta$ (%) | Interpretation |
|---|---|---|---|---|---|---|
| Baseline | 0.90 | 100 | 3× | 7.10 | 11.04 | Baseline |
| Dichroic 80% | 0.80 | 100 | 3× | 7.10 | 11.04 | Moderate routing loss |
| Dichroic 70% | 0.70 | 100 | 3× | 7.10 | 11.07 | Conservative routing loss |
| Narrow band | 0.90 | 50 | 3× | 7.10 | 11.07 | Narrower collection |
| Wide band | 0.90 | 150 | 3× | 7.11 | 11.08 | Broader collection |
| 2× concentration | 0.90 | 100 | 2× | 7.10 | 11.07 | Lower concentration |
| 1× concentration | 0.90 | 100 | 1× | 7.10 | 11.07 | No concentration |
| Diffuse/no-tracking | 0.70 | 100 | 1× | 7.09 | 11.01 | Conservative: low routing, no concentration |

**Headline assumptions.** Every $\eta$ in the main text is conditional on Table S22; sensitivity in Table S23.

Table S22: Headline assumptions for the process-efficiency calculation. OTR: oxygen transfer rate.

| Parameter | Value | Source/justification |
|---|---|---|
| $\eta_{PV}$ (SOTA) | 0.241 | High-efficiency PV[51] |
| $\eta_{PV}$ (Theoretical) | 0.330 | Shockley–Queisser |
| Electrolyser demand (SOTA) | 500 kJ mol$^{-1}$ | ≈2.6 V, ≈51% HHV |
| Electrolyser demand (Theoretical) | 270 kJ mol$^{-1}$ | Thermodynamic min (HHV ≈ 254.6 + overpotential) |
| Dichroic efficiency | 0.90 | Typical mirror |
| Absorption bandwidth | 100 nm FWHM | Dichroic photon-supply cap |
| Biomass HHV | 22.4 kJ g$^{-1}$ | Microbial energy density |
| $\mu$ | 0.126 h$^{-1}$ | Dark formatotrophic upper bound |
| CDW bound | $[0.05, 10]$ g L$^{-1}$ | Sustained autotrophic density |
| $V/A$ bound | $[0.005, 0.10]$ m | Thin-film to deep-tank |
| $O_2$ transfer | $\leq 100$ mmol L$^{-1}$ h$^{-1}$ | OTR range[52] |
| Formate feed | $\leq 100$ mmol L$^{-1}$ h$^{-1}$ | Substrate supply |
| Solar conc. | $\leq 3\times$ band PPFD | vs unconcentrated AM1.5G |
| Photon supply | dichroic solar | AM1.5G band via dichroic; no LED |
| Heat removal | ∼49 W L$^{-1}$ (non-binding) | Metabolic + solar at 45% optimum; within PBR cooling |

*Continued on next page*

*Table S22 continued*

| Parameter | Value | Source/justification |
|---|---|---|
| $CO_2/NH_4^+$ transfer | not modelled | $O_2$/formate limits above; others unmodelled |

**Process-sensitivity accounting.** Table S23 separates propagated from outside-boundary perturbations.

**Global optimisation audit.** Table S24 audits the headline whole-collector calculation itself. Twelve draws spanning each scenario's calculated distribution were re-optimised over all six continuous variables and every reactor geometry; $\Delta\eta$ is the full-space result minus the per-draw propagated-search result.

**Evidential standing.** Table S25 states each claim's evidential basis and the measurement that would test it; module-level falsification in Table S26.

**Biological module feasibility audit.** Table S26 consolidates biological quantities requiring experimental verification. Both columns use the 45% membrane cap. The reference is generated by `PaperScripts/pump_divergence/data/generate_membrane_feasible_reference.py`; the optimum uses the closed, capture-charged boundary.

**Membrane-capped stress sweep.** Table S27: rGlyP external cylinder re-optimised under rhodopsin-abundance caps. (a) Route boundary; (b) closed boundary (main text). $CO_2$ capture charged; each panel re-optimised independently. Sources: (a) `Results/reactor_geometry_wavelength/membrane_cap_optima_dac.csv`, (b) `Results/reactor_geometry_wavelength/hybrid_dichroic_led_dac.csv`. Panel (b) optimises dichroic $t \in [0, 1]$ and includes the no-aperture standard-PV dark branch.

**Cofactor-policy and PMF/ATP audit.** Table S28 checks whether the ATP-only interpretation depends on NADTRHD closure. Reopening to 100 mmol gDW$^{-1}$ h$^{-1}$ leaves formate demand unchanged at every operating point. The slack uses the same complete set of native respiratory proton sources as the LP constraint. It is zero when ATP synthase uses all available proton supply and positive when pumping exceeds its demand.

**Three-objective Pareto optima.** Three process objectives under the same constraints; no single point maximises all three.

### Figures

Table S23: **Sensitivity terms in the propagated uncertainty interval and risks outside it.**

| Perturbation | Tested range or status | Effect on optimised $\eta$ | In Monte Carlo? | Interpretation |
|---|---|---|---|---|
| $O_2$ transfer ceiling | 20–200 mmol $L^{-1}$ $h^{-1}$ | < 0.1 pp | Yes, bounded shift | Slack; rhodopsin displaces $O_2$ |
| Formate-feed ceiling | 50–300 mmol $L^{-1}$ $h^{-1}$ | < 0.1 pp | Yes, bounded shift | Slack |
| Solar-concentration / optical hardware | Concentration and solar-band delivery varied; dichroic transmission re-optimised per draw | Depends on denominator and occupancy | Yes, selected terms | Construction and ageing remain outside the interval |
| Rhodopsin absorption calibration $\sigma_{eff}$ | 0.5–2× forward pump scale | SOTA 2.69–4.63% at 45% occupancy | Yes, pump multiplier | Whole-collector calibration sensitivity; Table S5 |
| Quantum-yield uncertainty | Represented by an effective pump multiplier | Via pump input | Yes, approximate | No full draw-wise photophysics refit in process propagation |
| Biomass heating value | Normal around 22.4 kJ $g^{-1}$, truncated to 20.0–24.8 | Included in numerator | Yes | Sampled in every headline draw |
| $CO_2$ capture energy | SOTA triangular 300–550 kJ $mol^{-1}$; thermodynamic limit 19.2 | Included in formate electrical demand | Yes (SOTA); fixed (limit) | Included in every headline design |
| GAM and metabolic demand | ±10% GAM + structure multiplier | Dominant SOTA source | Yes, response slope | Main propagated uncertainty |
| Kin4B8 transfer, delivery and bandwidth | 0.50–0.70; 0.75–1.0; 0.8–1.2× width | Paired SOTA $\Delta\eta$: median +0.107 pp; 93.2% positive | Yes, joint antenna interval | Evidence-bounded epistemic ranges; not empirical confidence limits |
| Antenna assembly burden | No measured distribution; 6.7% break-even bound | Can erase the central optical gain | No | Must be measured in the constructed strain |
| Transhydrogenase and cofactor policy | NADTRHD closed; Table S28 | No change | No | Isolates NADPH balance; zero flux when reopened |
| Structural photophysics / metabolic-model error | Single-WT QY, unmeasured mutants, EC-FBA structure | No distribution assigned | No | Structural uncertainty; motivates region-level claims |
| Biological construction terms | Expression, folding, loading, membrane burden | Outside boundary | No | Experimental go/no-go |
| Balance-of-plant / TEA terms | Nutrients, most mixing/circulation, downstream recovery and drying, retinal supply, capital and downtime | Outside headline interval; two mixing duties charged separately | No | $CO_2$ capture is included; full deployment and economics are not |

Table S24: **Global-optimisation audit for the headline whole-collector uncertainty calculation.**

| Mode | Audit draws | Median $\Delta\eta$ (pp) | Max $\Delta\eta$ (pp) | Interpretation |
|---|---|---|---|---|
| SOTA | 12 | −0.000012 | 0.000001 | Maximum absolute difference 0.000047 pp |
| Theoretical | 12 | −0.000003 | 0.000020 | Maximum absolute difference 0.000040 pp |

Table S25: **Evidential basis and falsification test for each claim.**

| Claim | Rests on | What would test it |
|---|---|---|
| Dark pathway ordering | *C. necator* chemostat data[24,74] | Met. Unfitted advantage reproduces measured chemostat ratio |
| Rhodopsin supplies usable ATP | Light-dependent formate growth[73] | Formate demand under light/dark switching |
| rGlyP and rhodopsin combined | Components built separately[24,25,86] | Combined strain; rGlyP predicted to save more formate |
| Blue/blue-green design region | Integrated three-scale model | Multi-wavelength QY screen |
| Contextual crop values | Published seasonal conversions shown beside a steady-design model | No performance ordering is claimed; seasonal operation, site footprint and compatible biomass recovery would be required |

Table S26: **Feasibility audit for the rhodopsin ATP-supply module.** Reference: 100 PPFD, 568 nm and fixed growth. Optimum: state-of-the-art rGlyP external cylinder on the closed, capture-charged boundary. Both use the 45% membrane cap.

| Quantity | 100-PPFD reference | SOTA closed-boundary optimum | Interpretation / required check |
|---|---|---|---|
| Rho abundance | 2.13 $\mu$mol gDW$^{-1}$ | 2.13 $\mu$mol gDW$^{-1}$ (4.74 at ceiling) | Proteomics/absorbance calibration needed |
| Rho molecules | $1.3 \times 10^{18}$ gDW$^{-1}$ | $1.3 \times 10^{18}$ gDW$^{-1}$ ($2.9 \times 10^{18}$ at ceiling) | Folding/activity must be retained |
| Retinal demand | 0.60 mg gDW$^{-1}$ | 0.60 mg gDW$^{-1}$ (1.35 at ceiling) | Supply/loading must not erase gain |
| Membrane footprint | 9.0 of 20 m$^2$ gDW$^{-1}$ | 9.0 of 20 m$^2$ gDW$^{-1}$ | Design variable; requires folding and retained activity |
| Cap sweep | 100–0% nominal caps | Route SOTA: 4.80–2.32%; closed retinal-only (100/45/30%): 3.74/3.29/3.05%; no-aperture dark: 2.56% | Table S27; low caps collapse to dark |
| $H^+$ flux | 29.6 mmol gDW$^{-1}$ h$^{-1}$ | 87.3 mmol gDW$^{-1}$ h$^{-1}$ | Measure directly or infer from a calibrated abundance |
| ATP equiv. flux | 7.4 mmol gDW$^{-1}$ h$^{-1}$ (4 $H^+$/ATP) | 21.8 mmol gDW$^{-1}$ h$^{-1}$ | Depends on ATP-synthase coupling |
| Formate effect | 28.9 $\rightarrow$ 24.7 mmol gDW$^{-1}$ h$^{-1}$ (14.6% cut) | 18.7 mmol gDW$^{-1}$ h$^{-1}$ | Direct fixed-growth test |
| Pump-capacity screen | 0–100% nominal caps | 0–35.5 mmol gDW$^{-1}$ h$^{-1}$ | Measure before reactor-scale claims |

Table S27: **Re-optimised rGlyP external-cylinder designs under membrane-capped rhodopsin abundance.** (Related to Figure 7.) **(a)** Route boundary with $CO_2$ capture charged. **(b)** Closed boundary (main-text basis), $CO_2$ capture charged, dichroic transmission free; the 0% rows are the no-aperture standard-PV branch. Each panel is optimised independently. "Pump fraction": relative to uncapped inventory; $\alpha$: volume-averaged/surface PPFD; ATP flux: 4 $H^+$ $ATP^{-1}$; "Aperture": collector area per unit reactor footprint (panel (a) units as Table S17; $H^+$/ATP fluxes in mmol $gDW^{-1}$ $h^{-1}$). †Panel-(b) Theoretical $\lambda^\star$ lands on the 600-nm search edge because the dichroic routes almost all light through photovoltaics; it is a boundary corner, not a resolved preference (Section S6.12). At 45% occupancy this row also presses $V/A$ onto its upper bound and PPFD onto its lower bound (Section S6).

**(a) Route boundary, capture charged**

| Electrolyser | Nominal membrane cap | Pump fraction | $\lambda^\star$ (nm) | PPFD* | CDW* | $V/A^*$ | Areal CDW | $\alpha$ | $\eta_{route}$ (%) | $H^+$ flux | ATP equiv. |
|---|---|---|---|---|---|---|---|---|---|---|---|
| SOTA | 100% | 0.948 | 556.2 | 2056 | 1.06 | 0.0968 | 102.5 | 0.105 | 4.80 | 142.0 | 35.5 |
| SOTA | 80% | 0.758 | 553.3 | 2052 | 9.16 | 0.0105 | 95.9 | 0.132 | 4.69 | 141.8 | 35.5 |
| SOTA | 45% | 0.427 | 551.1 | 2049 | 9.14 | 0.0083 | 75.7 | 0.242 | 4.33 | 141.1 | 35.3 |
| SOTA | 40% | 0.379 | 550.9 | 2049 | 9.13 | 0.0078 | 71.2 | 0.274 | 4.23 | 140.9 | 35.2 |
| SOTA | 30% | 0.284 | 550.5 | 2049 | 0.61 | 0.0993 | 60.1 | 0.374 | 3.94 | 139.9 | 35.0 |
| SOTA | 25% | 0.237 | 550.3 | 2049 | 0.53 | 0.0996 | 53.1 | 0.457 | 3.72 | 138.8 | 34.7 |
| SOTA | 10% | 0.095 | 550.2 | 2049 | 0.64 | 0.0996 | 63.6 | 0.509 | 2.45 | 61.0 | 15.2 |
| SOTA | 5% | 0.047 | 585.8 | 50 | 3.14 | 0.0999 | 313.6 | 0.119 | 2.32 | 0.2 | 0.0 |
| SOTA | 0% | 0.000 | 568.0 | 50 | 3.14 | 0.1000 | 313.5 | 0.137 | 2.32 | 0.0 | 0.0 |
| Theoretical | 100% | 0.948 | 557.0 | 2057 | 1.40 | 0.0803 | 112.6 | 0.096 | 10.76 | 129.6 | 32.4 |
| Theoretical | 80% | 0.758 | 555.4 | 2055 | 1.14 | 0.0933 | 105.9 | 0.120 | 10.31 | 128.6 | 32.2 |
| Theoretical | 45% | 0.427 | 600.0 | 50 | 3.47 | 0.1000 | 347.0 | 0.053 | 11.13 | 0.8 | 0.2 |
| Theoretical | 40% | 0.379 | 593.0 | 50 | 3.15 | 0.1000 | 314.7 | 0.063 | 10.09 | 0.8 | 0.2 |
| Theoretical | 30% | 0.284 | 599.7 | 50 | 3.15 | 0.1000 | 314.5 | 0.072 | 10.09 | 0.7 | 0.2 |
| Theoretical | 25% | 0.237 | 600.0 | 50 | 3.14 | 0.1000 | 314.4 | 0.079 | 10.08 | 0.6 | 0.2 |
| Theoretical | 10% | 0.095 | 600.0 | 50 | 3.14 | 0.1000 | 314.0 | 0.106 | 10.07 | 0.3 | 0.1 |
| Theoretical | 5% | 0.047 | 600.0 | 50 | 3.14 | 0.1000 | 313.8 | 0.120 | 10.06 | 0.2 | 0.0 |
| Theoretical | 0% | 0.000 | 568.0 | 50 | 3.14 | 0.1000 | 313.5 | 0.137 | 10.05 | 0.0 | 0.0 |

**(b) Closed boundary (main text), capture charged**

| Electrolyser | Nominal membrane cap | $\lambda^\star$ (nm) | PPFD* | CDW* | $V/A^*$ | Aperture ($m^2$ $m^{-2}$) | $\eta_{closed}$ (%) |
|---|---|---|---|---|---|---|---|
| SOTA | 100% | 550.0 | 2048 | 4.96 | 0.0289 | 3.00 | 3.74 |
| SOTA | 45% | 550.0 | 2048 | 4.61 | 0.0273 | 3.00 | 3.29 |
| SOTA | 30% | 550.0 | 2047 | 4.71 | 0.0248 | 3.00 | 3.05 |
| SOTA | 0% | n/a | 0 | n/a | n/a | n/a | 2.56 |
| Theoretical | 100% | 600.0† | 50 | 3.47 | 0.1000 | 2.50 | 10.90 |
| Theoretical | 45% | 600.0† | 50 | 3.47 | 0.1000 | 2.50 | 10.89 |
| Theoretical | 30% | 600.0† | 50 | 3.47 | 0.1000 | 2.50 | 10.88 |
| Theoretical | 0% | n/a | 0 | n/a | n/a | n/a | 11.15 |

Table S28: **Cofactor-policy and PMF/ATP audit (fixed-growth formate min).** NADTRHD closed vs open (100 mmol $gDW^{-1}$ $h^{-1}$); other flux units as Table S17.

| Pathway | Policy | PPFD | Formate | $O_2$ | Rhodopsin $H^+$ | ATP synthase | Proton-source slack | NADTRHD | THD2 |
|---|---|---|---|---|---|---|---|---|---|
| rGlyP | Closed | 0 | 28.89 | 9.03 | 0.0 | 31.54 | 0.00 | 0.00 | 5.65 |
| rGlyP | Open | 0 | 28.89 | 9.03 | 0.0 | 31.54 | 0.00 | 0.00 | 5.65 |
| rGlyP | Closed | 100 | 21.29 | 5.23 | 53.2 | 31.54 | 0.00 | 0.00 | 5.65 |
| rGlyP | Open | 100 | 21.29 | 5.23 | 53.2 | 31.54 | 0.00 | 0.00 | 5.65 |
| rGlyP | Closed | 1915 | 10.87 | 0.01 | 127.9 | 31.55 | 1.75 | 0.00 | 5.65 |
| rGlyP | Open | 1915 | 10.87 | 0.01 | 127.9 | 31.55 | 1.75 | 0.00 | 5.65 |
| CBB | Closed | 0 | 34.06 | 11.61 | 0.0 | 40.59 | 0.00 | 0.00 | 2.72 |
| CBB | Open | 0 | 34.06 | 11.61 | 0.0 | 40.59 | 0.00 | 0.00 | 2.72 |
| CBB | Closed | 100 | 28.35 | 8.76 | 51.2 | 40.66 | 0.00 | 0.00 | 2.38 |
| CBB | Open | 100 | 28.35 | 8.76 | 51.2 | 40.66 | 0.00 | 0.00 | 2.38 |
| CBB | Closed | 1915 | 10.87 | 0.01 | 163.0 | 40.74 | 0.00 | 0.00 | 2.31 |
| CBB | Open | 1915 | 10.87 | 0.01 | 163.0 | 40.74 | 0.00 | 0.00 | 2.31 |

Table S29: Pareto-optimal designs, rGlyP external cylinder. (Related to Figure 7.) Bold = maximised KPI. Route boundary without capture (cf. Table S17). Max-material and max-productivity rows sit on degenerate ridges.

| Objective | Electrolyser | $\lambda^*$ (nm) | $f^*_{\mathrm{mem}}$ | CDW* (g $L^{-1}$) | $V/A^*$ (m) | PPFD* ($\mu$mol $m^{-2}$ $s^{-1}$) | $\eta_{\mathrm{route}}$ (%) | Yield (g biomass $mol^{-1}$ formate) | Productivity (g $L^{-1}$ $h^{-1}$) |
|---|---|---|---|---|---|---|---|---|---|
| Max energy | SOTA | 556.4 | 1.000 | 9.088 | 0.0114 | 2056 | **7.09** | 11.47 | 1.145 |
| Max energy | Theoretical | 557.1 | 1.000 | 8.212 | 0.0138 | 2057 | **11.05** | 10.36 | 1.035 |
| Max material | SOTA | 516.6 | 0.752 | 0.375 | 0.0103 | 484 | 2.03 | **11.60** | 0.047 |
| Max material | Theoretical | 516.6 | 0.752 | 0.375 | 0.0103 | 484 | 2.25 | **11.60** | 0.047 |
| Max productivity | SOTA | 546.7 | 0.893 | 9.204 | 0.0050 | 1133 | 6.42 | 11.60 | **1.160** |
| Max productivity | Theoretical | 546.7 | 0.893 | 9.204 | 0.0050 | 1133 | 9.31 | 11.60 | **1.160** |

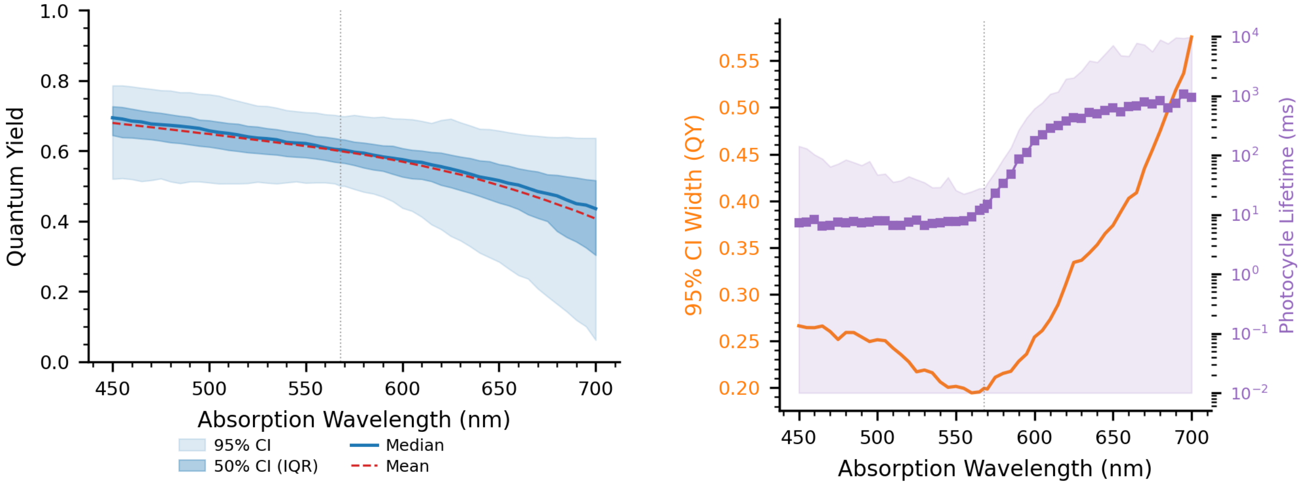


Figure S1: **Posterior-predictive quantum yield across the design sweep.** (Related to Figure 5.) Predictions over 450–700 nm, marginalised over the four-parameter posterior ($m$, $\gamma$, $c_s$, $\Delta E_{\rm CI}$) and the unconstrained $\beta \sim U(0,1)$ prior; shaded = 95% credible interval, solid = median. The second panel gives the width of that interval against wavelength, narrowest at the 568 nm calibration anchor and widening in both directions away from it, with the photocycle lifetime on the secondary axis (Figure S4 resolves the same quantities against the extended sweep). Ensemble: 800 post-burn-in steps $\times$ 32 walkers (25,600 draws).

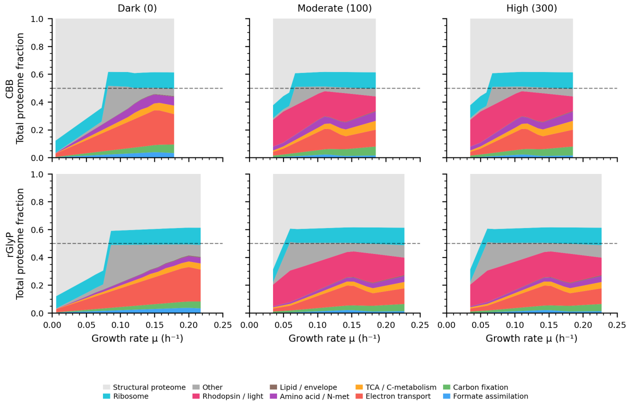


(a)

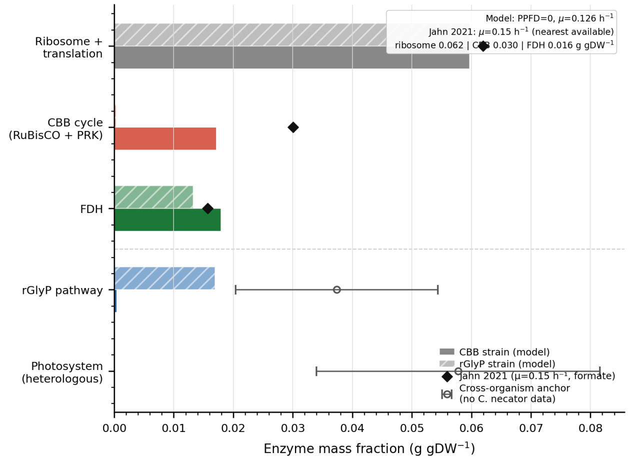


(b)

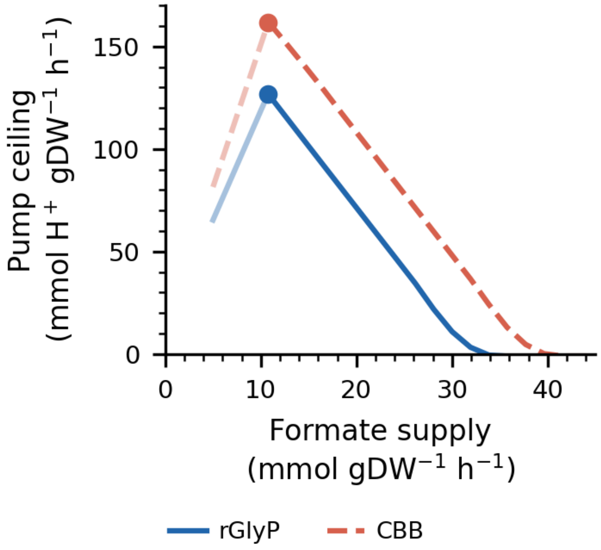


(c)

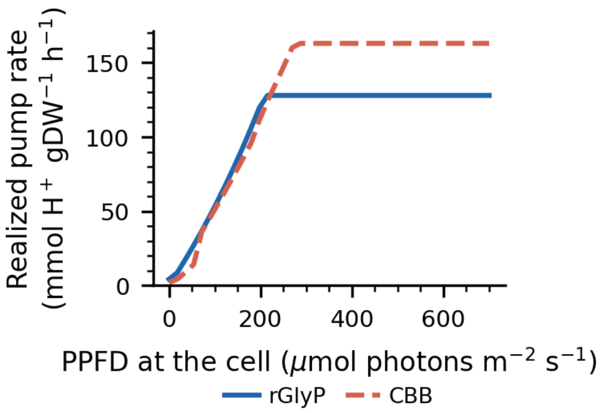


(d)

Figure S2: **GECKO proteome allocation, validation against *C. necator* proteomics, and the saturation ceiling.** (Related to Figure 4.) **(a)** Pool allocation across enzyme categories: stacked areas showing budget partitioning ($P_{\text{total}} = 0.68$ g gDW$^{-1}$, $f_{\text{enzyme}} = 0.50$) as growth rate varies. Top row CBB, bottom rGlyP; columns: no pump, moderate (100), high (300 mmol H$^+$ gDW$^{-1}$ h$^{-1}$). Dashed line: enzyme-accessible fraction. **(b)** Validation: dark (PPFD = 0, $\mu = 0.126$ h$^{-1}$) enzyme-mass fractions (bars; solid = CBB, hatched = rGlyP) against Jahn et al. proteomics (diamonds) and cross-organism bands (open circles), in g protein gDW$^{-1}$; sector-by-sector assessment in Section S11. **(c)** Proteome saturation flux against formate uptake: CBB's ceiling falls with formate (Calvin-cycle enzymes pre-load the pool), whereas rGlyP's is approximately flat (membrane-limited). **(d)** Photon utilisation efficiency at $\mu = 0.126$ h$^{-1}$: rGlyP converts more across the full range.

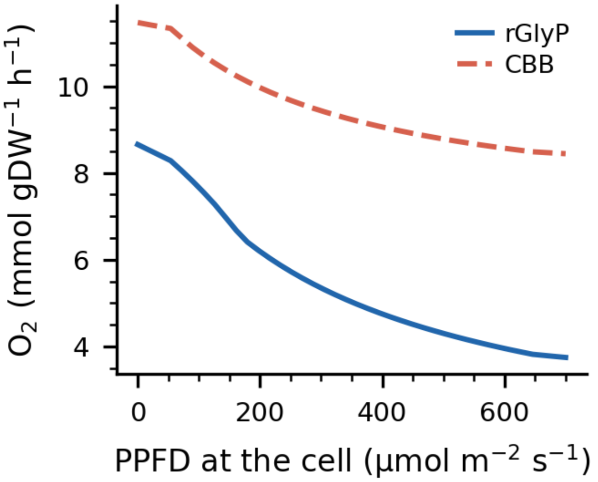


(a)

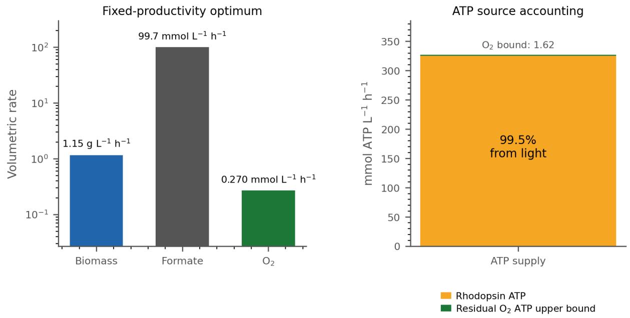


(b)

Figure S3: **Respiratory oxygen demand falls as light-driven ATP displaces formate oxidation.** (Related to Figures 3 and 7.) **(a)** $O_2$ consumption against pump rate at fixed dark formate. CBB requires ∼29% more $O_2$ than rGlyP at dark baseline and retains higher demand across the full pump range. **(b)** Stoichiometric sanity check at the constrained optimum (externally illuminated cylindrical rGlyP/SOTA): biomass production 1.153 g L$^{-1}$ h$^{-1}$, formate uptake 99.7, residual $O_2$ uptake 0.270 mmol L$^{-1}$ h$^{-1}$. At six ATP per $O_2$, residual respiration supplies only ∼1.6 mmol ATP L$^{-1}$ h$^{-1}$ against ∼325 from proton pumping, so the low $O_2$ flux in Fig. 7 reflects ATP displacement, not an aeration bottleneck.

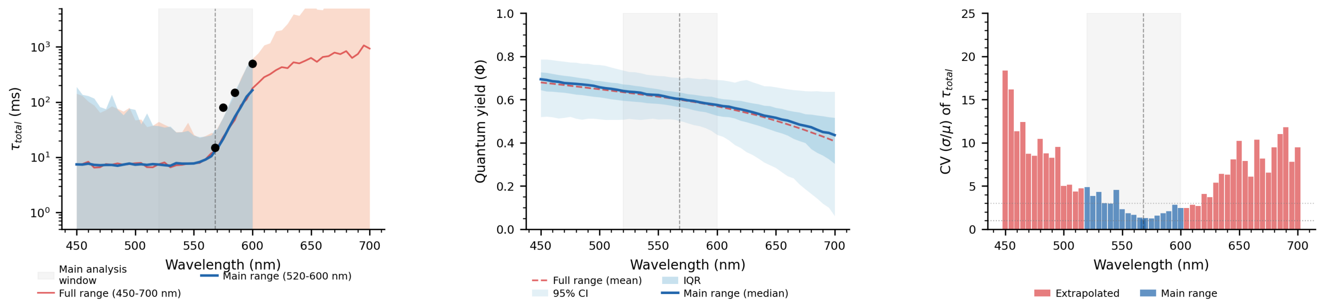


Figure S4: **How the predictive interval widens across the extended 450–700 nm sweep.** (Related to Figure 5.) **a,** $\tau_{\text{total}}$ with $\pm 1\sigma$ bands. **b,** Quantum yield: the main-range posterior median and credible bands against the extended-sweep mean (dashed). The extended sweep evaluates the simulator at every wavelength it covers, so the dashed mean is a prediction throughout rather than a value carried over from the main window. **c,** Coefficient of variation of $\tau_{\text{total}}$, smallest at the 568 nm wild-type anchor (1.16) and rising toward both ends of the range, to 18.4 at 450 nm (some 16× the anchor value) and 9.5 at 700 nm (`PaperScripts/physics_data/supplementary_full_range/physics_sweep.csv`). The main analysis uses 450–600 nm. Posterior as in Figure S1.

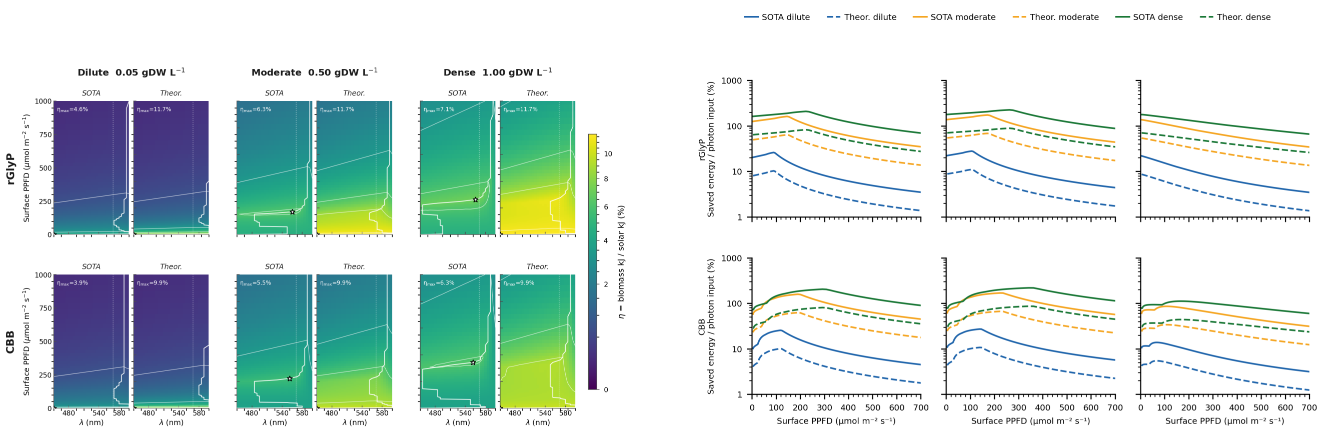


(a) (b)

Figure S5: **Reactor-aware process and formate-saving efficiency.** (Related to Figure 7.) Both panels use fixed-growth conditions ($\mu = 0.126\ \mathrm{h}^{-1}$, formate minimised, EC-class saturation) with surface PPFD propagated through the externally illuminated cylinder in the perfect-mixing limit. The LP receives no fixed pump-rate bound: turnover $r_{\mathrm{pump}} = [1 - \exp(-k_{\mathrm{abs}}\Phi_{\mathrm{QY}}\tau)]/\tau$ is computed per grid point, and realised pump flux is an LP output. **(a)** Process energy efficiency: biomass chemical energy divided by PV energy for formate plus incident photon energy per gDW. **(b)** Formate-saving efficiency: PV energy for formate saved (relative to dark baseline) divided by incident photon energy. Sub-panels show rGlyP and CBB at 450, 568 and 600 nm for two electrolysis-energy assumptions.

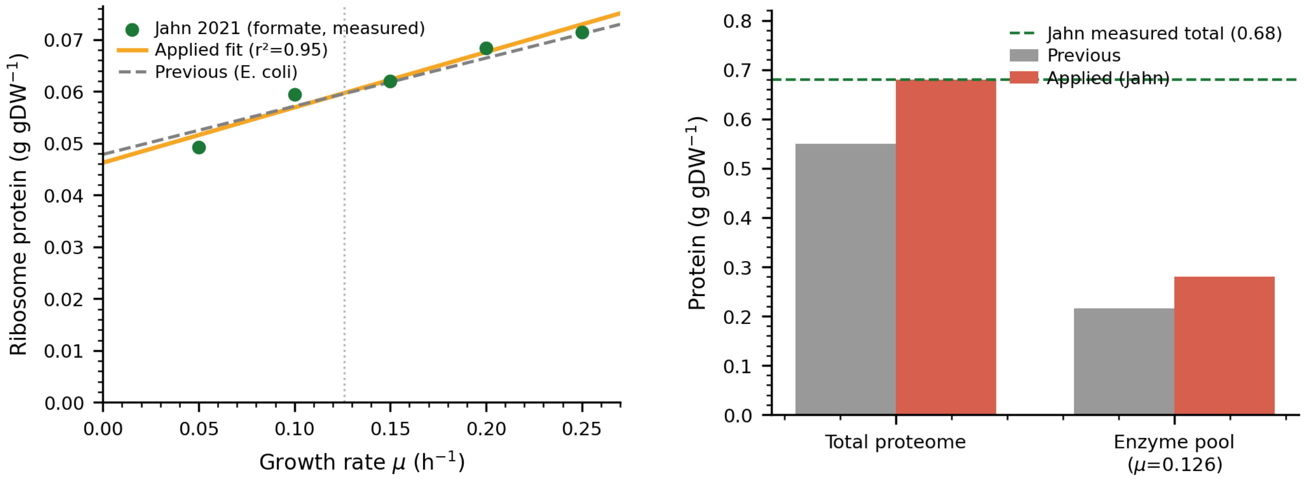


Figure S6: **Calibration of the GECKO proteome budget and ribosome sector to measured *C. necator* proteomics[23].** (Related to Figure 4.) **a,** Ribosomal protein mass against growth rate on formate: measured values (five chemostat dilution rates) with the applied ordinary-least-squares fit ($f_{\text{ribo}}(\mu) = 0.068+0.157\,\mu$, $r^2 = 0.95$) replacing the previous *E. coli* parameterisation (grey dashed); dotted line marks the setpoint. **b,** Total proteome budget and the resulting effective enzyme pool $P_{\text{total}}(f_{\text{enzyme}} - f_{\text{ribo}})$ at the setpoint, before and after calibration, with the measured total (0.68 g gDW$^{-1}$, $\mu$-invariant) as the dashed line. Calibration raises $P_{\text{total}}$ from 0.55 to 0.68 and the enzyme pool from 0.22 to 0.28 g gDW$^{-1}$. Mapping model genes onto the same measured proteome is consistent with retaining $f_{\text{enzyme}} = 0.50$, but is not an independent validation because the host proteomics also calibrate the envelope.

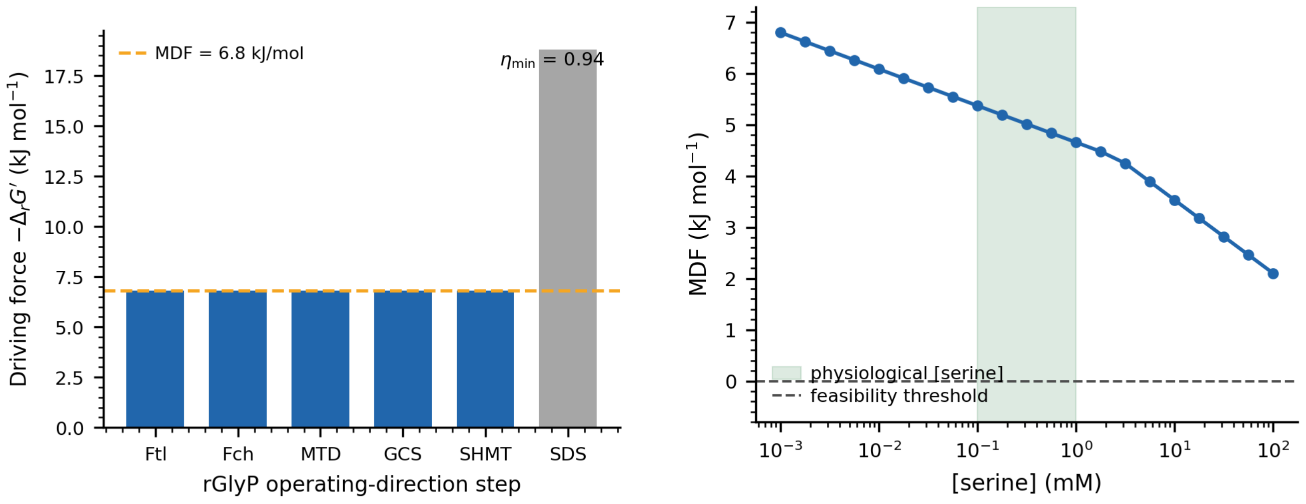


Figure S7: **The rGlyP route is thermodynamically feasible under physiological concentrations, justifying the reverse-$k_{cat}$ treatment (Section S8).** MDF analysis (eQuilibrator $\Delta_r G'^0$ at pH 7, $I = 0.25$ M, pMg 3[61,62]). **a,** Per-reaction driving force at the MDF optimum. The five C1/THF steps (Ftl, Fch, MTD, GCS, SHMT) co-limit at MDF = +6.8 kJ mol$^{-1}$ (dashed), with $\eta_{min} \approx 0.94$ ($\sim$7% thermodynamic enzyme surcharge); serine deaminase is strongly forward-driven. **b,** MDF against pinned serine concentration: feasibility holds from 1 $\mu$M to 100 mM, declining gently across the physiological window (shaded), refuting a standard-state analysis flagging SHMT as a thermodynamic wall. This does not replace genome-scale concentration-resolved analysis, and the reverse-$k_{cat}$ exemption remains a sensitivity assumption.

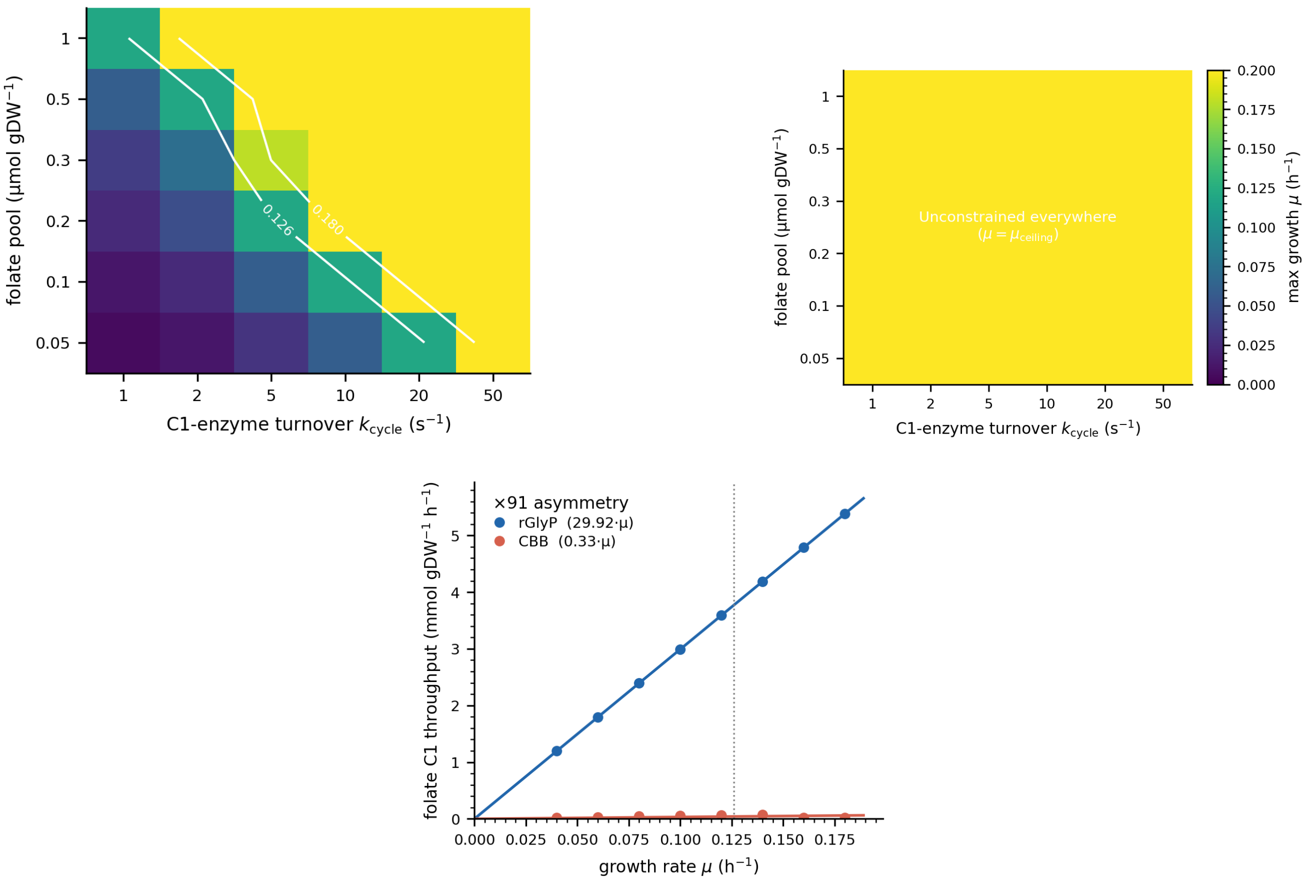


Figure S8: **The rGlyP advantage persists across folate C1-cycle turnover except in a small-pool, low-saturation corner.** The steady-state model enforces C1 *flux* but does not track folate pool size; a high C1 flux can exhaust a small pool. We impose $J_{C1} \leq F_{pool}\, k_{cycle}$ separately and map maximum growth across pool size and turnover. **a,** rGlyP maximum $\mu$; white contours: setpoint (0.126 $h^{-1}$) and batch ceiling (0.18 $h^{-1}$). The constraint binds only in the small-pool, low-turnover corner. **b,** CBB: C1 demand is so small the constraint never binds. **c,** Folate throughput versus $\mu$, exactly proportional (rGlyP $= 29.7\,\mu$, CBB $= 0.56\,\mu$ mmol gDW$^{-1}$ h$^{-1}$, $R^2 = 1$), a $\times 53$ asymmetry. At a central pool estimate ($\approx 0.3$ µmol gDW$^{-1}$) rGlyP needs $k_{cycle} \gtrsim 3.5$–$4.9$ s$^{-1}$, within reported turnover; the bottleneck bites only for pools $\lesssim 0.1$ µmol gDW$^{-1}$ or substantially undersaturated enzymes. Pool sizes follow *E. coli* ranges absent a *C. necator* measurement.

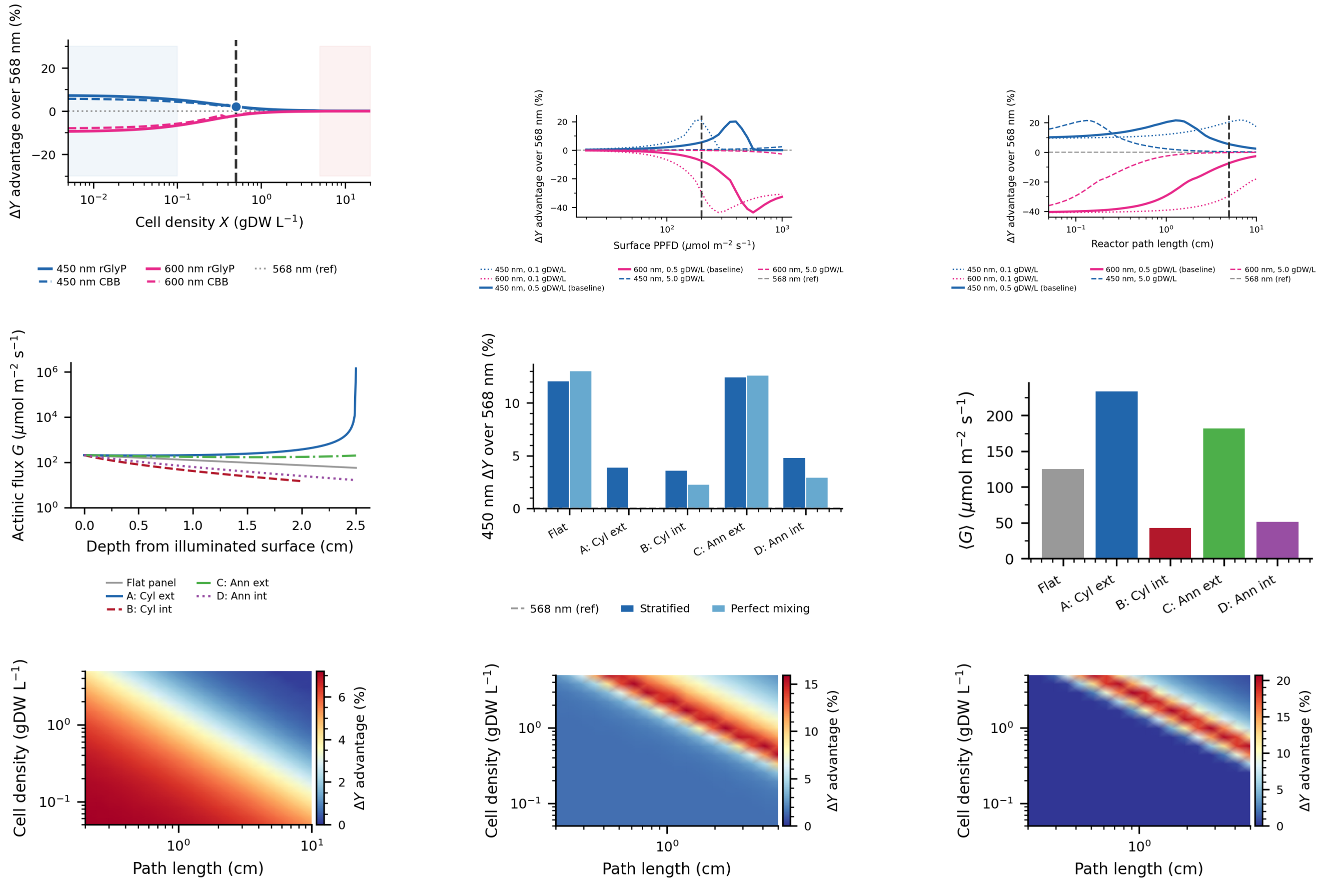


Figure S9: **Reactor-optics sensitivity.** (Related to Fig. 6.) Baseline: 100 PPFD, 0.5 gDW $L^{-1}$ and $5.0 \times 10^{-6}$ mol rhodopsin $gDW^{-1}$ at $\mu = 0.126$ $h^{-1}$. **a–c,** Flat-panel spectral advantage against density, PPFD and path length. **d–f,** Actinic flux, geometry and mixing. **g–i,** Operating windows for the flat panel and external cylinder. The production model uses background absorption of 0.02 $m^2$ $gDW^{-1}$; wavelength-dependent host absorption is tested in Section S5.

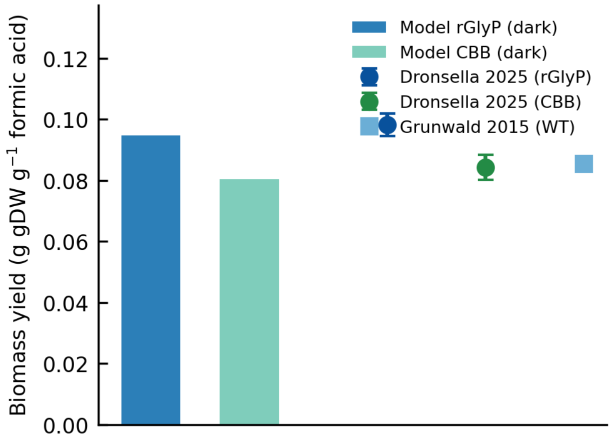


(a)

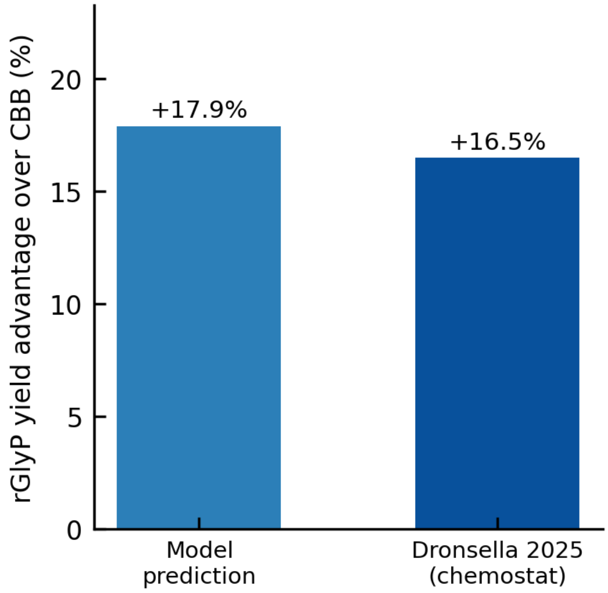


(b)

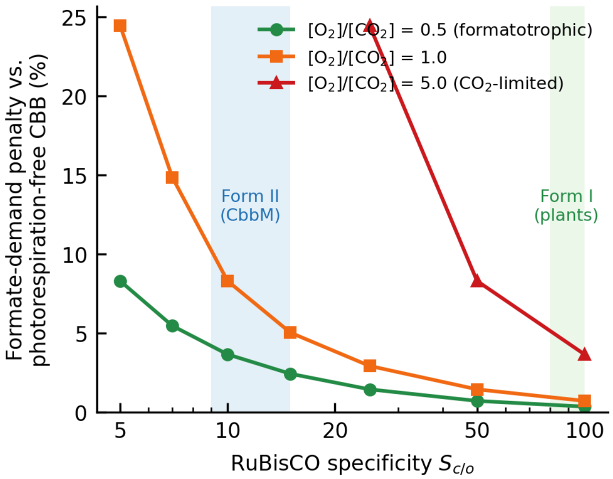


(c)

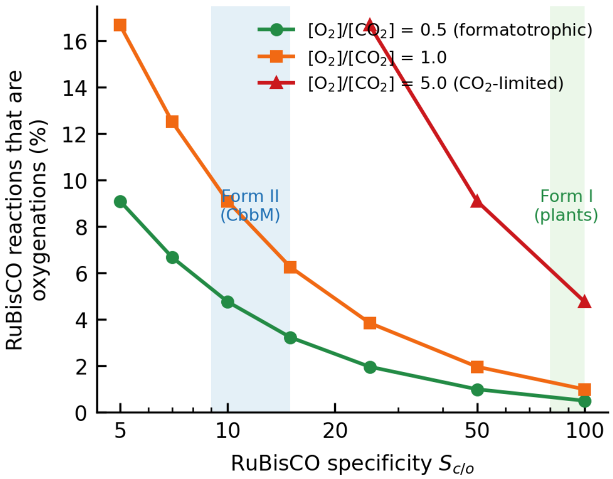


(d)

Figure S10: **Dark formatotrophic baseline benchmark and RuBisCO sensitivity.** (Related to Figure 2.) **(a,b)** Dark (PPFD = 0) fixed-growth optima against published *C. necator* chemostat yields[24,74] (g CDW per g formic acid). Yield is emergent, not fitted (Section S12). **(a)** Biomass yield: model sits 3.5–4.7% below measurement (rGlyP 4.362, CBB 3.699 g CDW $mol^{-1}$), i.e. conservative, preserving rGlyP > CBB. **(b)** rGlyP-over-CBB advantage matches measurement: +17.9% predicted vs. +16.5% observed[24]. **(c,d)** RuBisCO specificity barely affects CBB formatotrophy because $CO_2$ flooding suppresses photorespiration (Section S13). CBB solved with oxygenase and native salvage, sweeping $S_{c/o}$ at three $[O_2]/[CO_2]$ ratios ($\phi = ([O_2]/[CO_2])/S_{c/o}$); shaded bands: Form II ($S_{c/o} \approx 10$–15) and Form I ($\approx$80–100). **(c)** Formate penalty vs. photorespiration-free baseline. **(d)** Oxygenation fraction. See Section S13 for the non-binding turnover result and the $O_2$-sensitive variant of McDonald et al.[75].

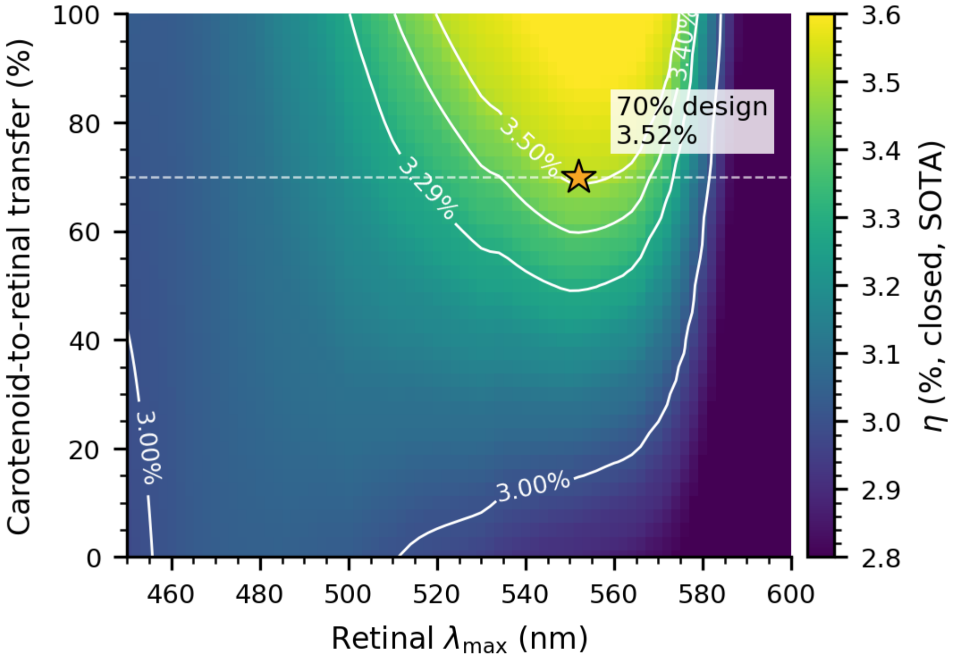


Figure S11: **Joint retinal–carotenoid equivalent-photon sensitivity at the** 45% **membrane-occupancy reference.** (Related to Figure 7.) Closed, capture-charged state-of-the-art efficiency is evaluated over retinal $\lambda_{\max}$ and carotenoid-to-retinal excitation-transfer efficiency for the measured Kin4B8 antenna envelope (432/457/487 nm). Surface PPFD follows the active 3× concentration boundary; density and depth are re-optimised along the model-fit 70% case's constant-areal-loading ridge; and dichroic transmission is re-optimised at every map point. The star marks that 70% sensitivity ($\lambda_{\max} = 552$ nm, $\eta = 3.52\%$).

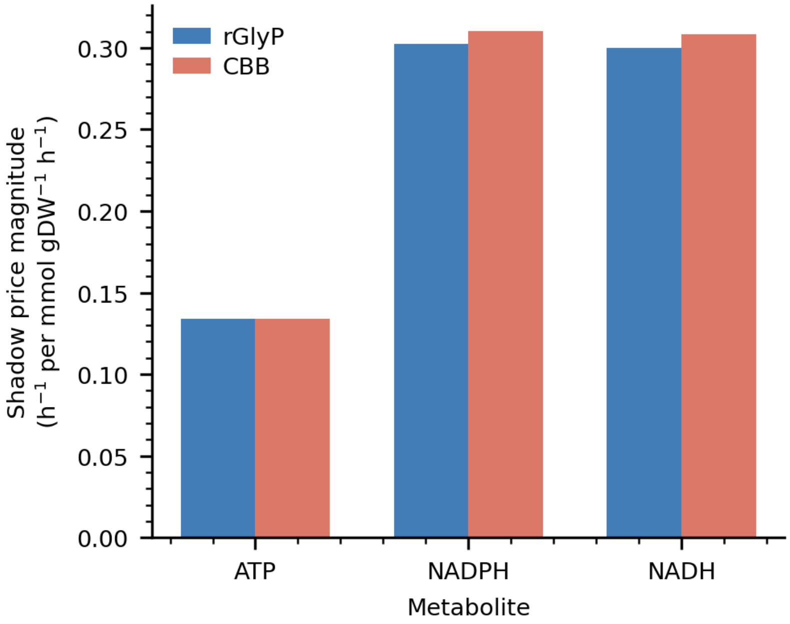


Figure S12: **EC-FBA shadow-price magnitudes at the fixed-growth reference.** (Related to Figure 4.) NADPH and NADH are approximately twice as constraining as ATP for both pathways (CBB slightly higher), so the dark baseline is limited by reducing power rather than ATP. Cofactor ratios and growth-associated ATP demand are given numerically in the main text.

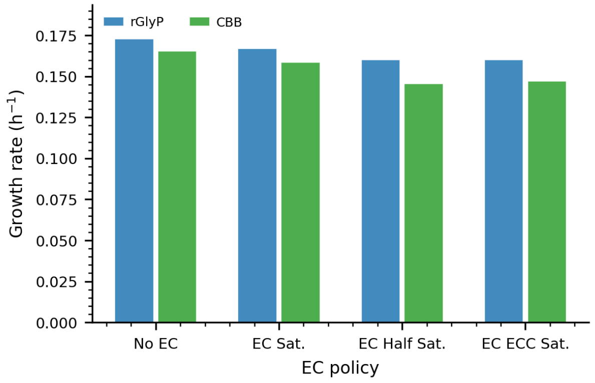


(a)

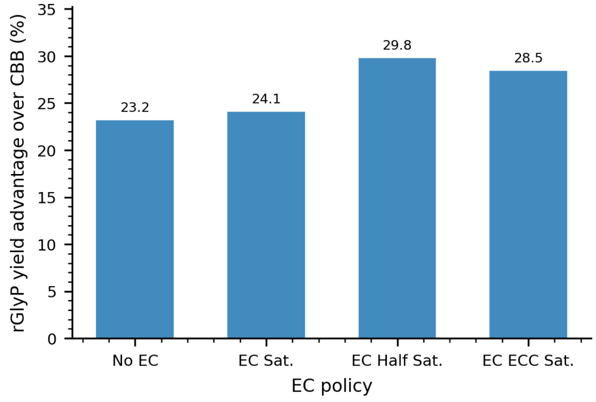


(b)

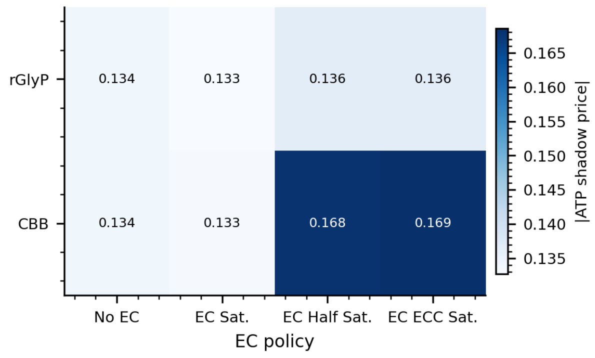


(c)

Figure S13: **Pathway comparison across enzyme-capacity policies.** (Related to Figure 2.) All panels evaluate the four EC saturation policies at the fixed pump-flux reference (deterministic LP optima). **(a)** Growth rate: rGlyP > CBB in every policy. **(b)** rGlyP yield advantage (%): positive throughout, largest under half-saturation and EC-class-specific saturation. **(c)** ATP shadow-price magnitude; darker cells mark stronger marginal formate-saving value. The active constraint structure is unchanged across policies.

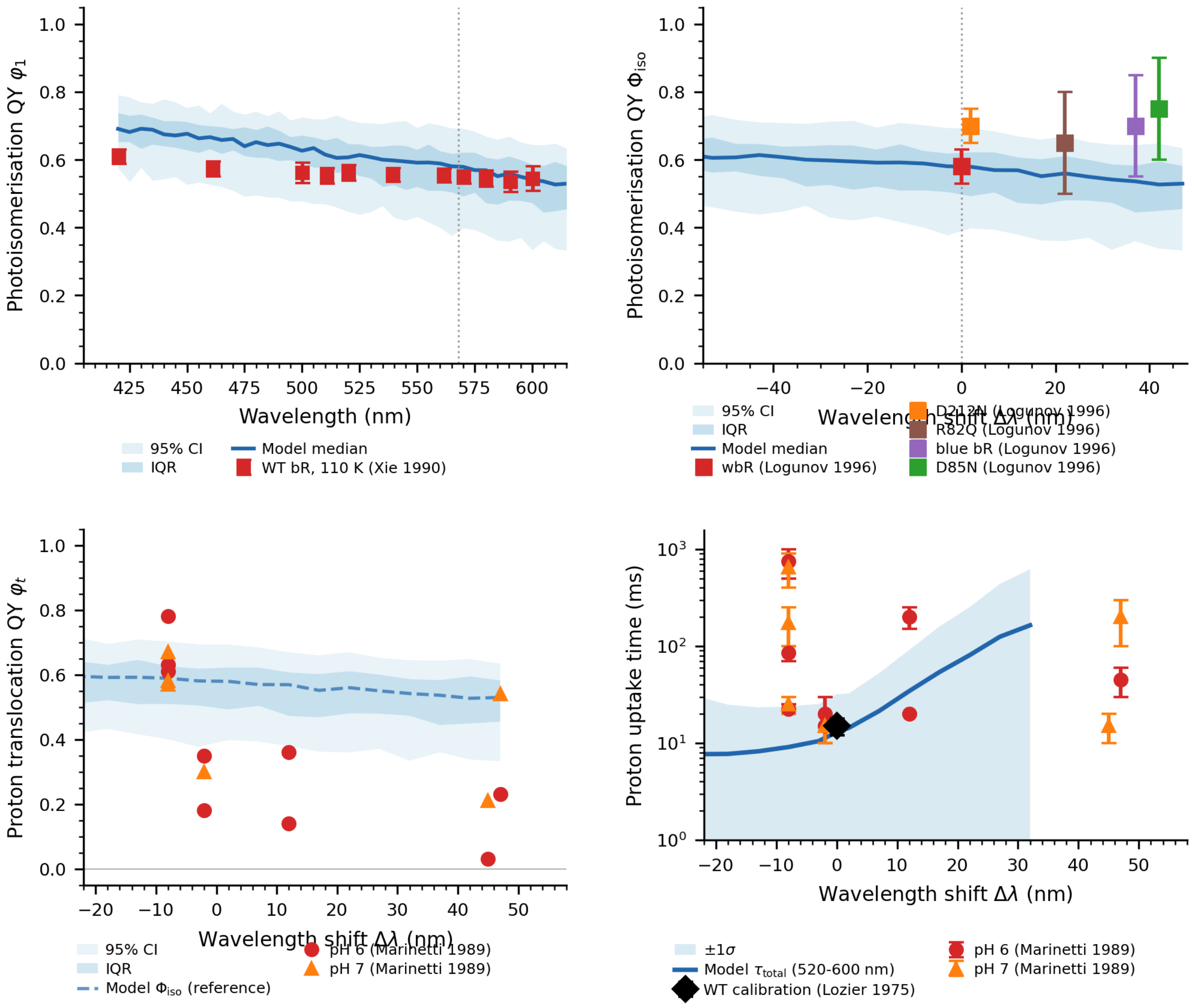


Figure S14: **Photophysics model calibration against published data.** (Related to Figure 5.) Blue shading = model posterior (95% CI, IQR, median). **a,** Xie[63]: WT bR quantum yield at 110 K against excitation wavelength; data fall within the model IQR. **b,** Logunov et al.[14]: $\Phi_{iso}$ for WT and four spectral variants at room temperature; the model captures WT but not the upward trend in red-shifted mutants, which reflects residue-specific effects. **c,** Marinetti et al.[60]: proton translocation quantum yield at pH 6 and pH 7, noting $\varphi_t \neq \Phi_{iso}$. **d,** Proton uptake time against $\Delta\lambda$ overlaid on model $\tau_{total}$ ($\pm1\sigma$); the D96N outlier reflects loss of the proton donor and lies outside the spectral-shift model's scope. Posterior as in Figure S1.

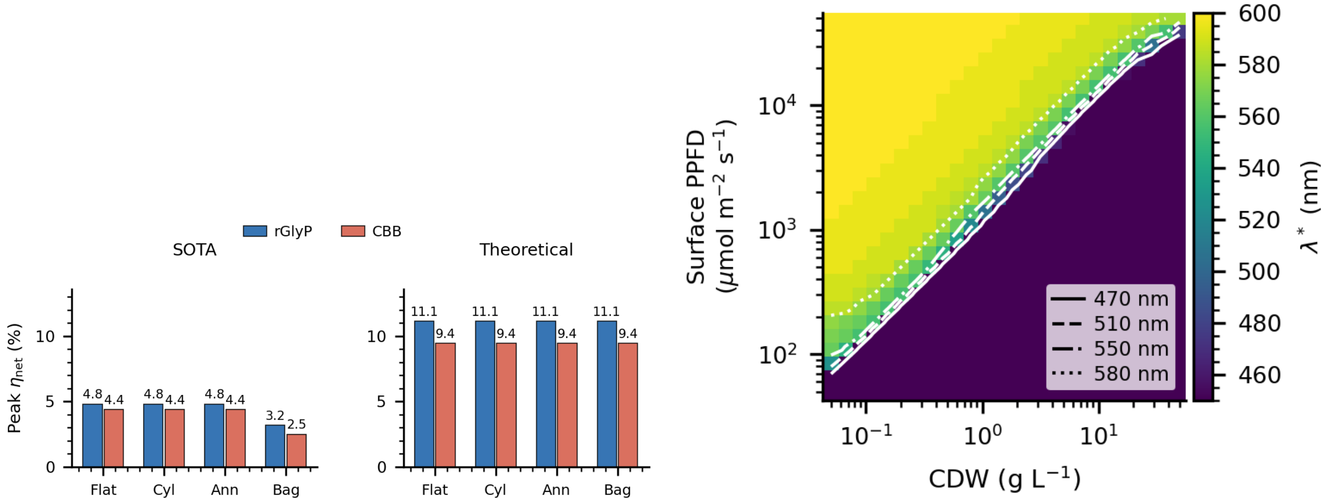


Figure S15: **Process-design details from the wavelength–reactor optimisation.** (Related to Fig. 7.) **a,** Constrained-optimal $\eta_{net}$ on the route boundary (Table S10) for each geometry, pathway and electrolyser scenario. This supplementary route diagnostic is separate from the closed, capture-charged, 45%-occupancy carotenoid–retinal result in Fig. 7. **b,** Optimal absorption maximum $\lambda^*$ for the externally illuminated cylinder (SOTA, closed boundary) across density and concentrated surface PPFD, with contours at 470, 510, 550 and 580 nm. The optimum is unconstrained in wavelength and therefore not pinned to WT bacteriorhodopsin or to the bluest design point.
